# Cold-mediated regulation of systemic retinol transport controls adipose tissue browning

**DOI:** 10.1101/2020.02.07.938688

**Authors:** Anna Fenzl, Oana Cristina Kulterer, Katrin Spirk, Goran Mitulović, Rodrig Marculescu, Martin Bilban, Sabina Baumgartner-Parzer, Alexandra Kautzky-Willer, Lukas Kenner, Jorge Plutzky, Loredana Quadro, Florian W. Kiefer

**Affiliations:** Clinical Division of Endocrinology and Metabolism, Department of Medicine III, Medical University of Vienna, Vienna, Austria; Clinical Institute of Laboratory Medicine, Medical University of Vienna, Vienna, Austria; Clinical Institute of Pathology, Medical University of Vienna, Vienna, Austria; Cardiovascular Medicine, Brigham and Women’s Hospital, Harvard Medical School, Boston, Massachusetts, USA; Department of Food Science and Rutgers Center for Lipid Research and New Jersey Institute of Food Nutrition and Health, Rutgers University, New Brunswick, NJ 08901, USA

## Abstract

Browning of white fat reduces obesity in many preclinical models. Vitamin A metabolites (retinoids) have been linked to thermogenic programming of adipose tissue (AT), however the physiologic importance of systemic retinoid metabolism for AT browning is unknown. Here we show that cold stimulation in mice and humans increases circulating retinol and its plasma transporter, retinol binding protein (RBP). Cold exposure shifts retinol abundance from liver towards subcutaneous white AT which correlates with enhanced thermogenic gene transcription. Cold-mediated retinoid flux is abrogated in Rbp deficient (Rbp^-/-^) mice and AT browning is dramatically impaired, which renders Rbp^-/-^ mice cold intolerant. Rbp deficiency attenuates cold-induced lipid clearance due to decreased oxidative capacity. In humans, cold-mediated retinol increase is associated with enhanced lipid utilization. Retinol stimulation in primary human adipocytes promotes thermogenic gene expression and mitochondrial respiration. In conclusion, coordinated retinol delivery is essential for cold-induced thermogenic programming of white fat.

## INTRODUCTION

During obesity development, excess energy intake is mainly stored in white adipose tissue in the form of triglycerides (Schutz, 1995). In contrast, brown adipose tissue oxidizes fatty acids and dissipates energy through uncoupled respiration and thermogenesis, a process mediated primarily by the key thermogenic factor uncoupling protein 1 (UCP1) (Cannon & Nedergaard, 2004; Seale, Kajimura, & Spiegelman, 2009). Promoting brown fat function *in vivo* has consistently demonstrated a decrease in adiposity and related metabolic complications in multiple preclinical models (Sellayah, Bharaj, & Sikder, 2011; Tseng et al., 2008; Whittle et al., 2012). In humans, BAT activity varies inversely with obesity and appears inducible by cold exposure. In addition to activating existing BAT, the induction of UCP1 expressing beige/brite adipocytes within white fat, a phenomenon termed *browning*, has been raised as another alternative increasing energy expenditure and promoting a lean phenotype (Kiefer, Orasanu, et al., 2012; Seale et al., 2011; Vegiopoulos et al., 2010). Thus, understanding the molecular mechanisms that lead to chronic BAT activation and/or browning of WAT represents a critical step in the development of novel therapeutic approaches for the treatment of obesity.

Retinoid metabolism (vitamin A and its derivatives) is known to be integral in regulating energy balance through actions in adipose tissue and the liver (Kiefer, Vernochet, et al., 2012; Mercader et al., 2006; Villarroya, Iglesias, & Giralt, 2004). Retinoids are diverse signaling molecules with distinct, essential biological actions, including the effect of specific retinoid species that can modulate the activity of the nuclear receptors retinoic acid receptor (Huang et al.) and retinoid X receptor (RXR) (Ross, 1993; Ziouzenkova & Plutzky, 2008). Physiologic retinoid concentrations depend on vitamin A intake, tissue storage and subsequent modification through a complex enzymatic network. Vitamin A must be obtained through dietary intake of either preformed vitamin A (retinol and/or retinyl esters) or provitamin A (carotenoids) (Frey & Vogel, 2011). Retinol is stored in form of esters predominantly in hepatic stellate cells (Blaner et al., 2009). In plasma, retinol is transported to target tissues bound to retinol binding protein (RBP or RBP4) (Noy, 2000; Ross, 1993).

Once taken up by target cells, retinol is reversibly oxidized to retinaldehyde by alcohol- and retinol dehydrogenases followed by irreversible oxidation to retinoic acid by retinaldehyde dehydrogenases (Frey & Vogel, 2011; Kiefer, Orasanu, et al., 2012). Most actions of the vitamin A pathway are considered to be mediated by the metabolite retinoic acid (Alvarez et al., 2000; Alvarez et al., 1995; Berry, DeSantis, Soltanian, Croniger, & Noy, 2012; Mercader, Palou, & Bonet, 2010; Noy, 2000; Puigserver, Vazquez, Bonet, Pico, & Palou, 1996; Rabelo, Reyes, Schifman, & Silva, 1996; Ribot, Felipe, Bonet, & Palou, 2001, 2004; Ross, 1993).

Previous reports in mice revealed an association between global vitamin A deficiency and obesity. Retinoids inhibit adipogenesis in vitro while retinoid administration in vivo can limit obesity (Mercader et al., 2006; Schwarz, Reginato, Shao, Krakow, & Lazar, 1997; Ziouzenkova et al., 2007). However, the molecular mechanisms underlying these responses and phenotypes have remained incompletely understood. Retinoic acid stimulation of adipocytes induces expression of BAT marker genes such as UCP1 (Mercader et al., 2010; Puigserver et al., 1996; Ribot et al., 2004). We reported that raising retinaldehyde levels by retinaldehyde dehydrogenase 1 deficiency or antisense induced a WAT thermogenic program increasing energy expenditure and protecting against obesity (Kiefer, Vernochet, et al., 2012). These findings combine with the extensive vitamin A stores in the liver to raise the hypothesis that hepatic retinoid metabolism helps regulate cold adaption, thermogenesis and systemic energy balance by retinoid formation and release to target tissues.

Here we show that cold exposure in mice and humans increased circulating vitamin A and RBP concentrations accompanying significant decreases in hepatic retinoid stores and increases in WAT and BAT retinol levels. Cold-mediated retinoid flux from the liver to WAT was absent in Rbp deficient (Rbp^-/-^) mice, a genetic mouse model with defective hepatic retinoid mobilization and systemic transport, resulting in impaired cold-induced WAT browning, perturbed mitochondrial function and fat oxidation, and significant cold intolerance. These findings establish that cold exposure is tightly coupled to systemic vitamin A levels and that intact hepatic retinoid shuttling to WAT is essential for executing the thermogenic program in adipose tissue. Understanding the biological role of this previously unknown liver-fat axis in adaptive thermogenesis and translating such pathways into human biology could help identify novel targets for treating obesity and its complications.

## RESULTS

### Cold exposure regulates systemic retinoid levels in mice and men

To study the effects of cold exposure on systemic vitamin A metabolism, we first investigated cold-mediated regulation of retinol and RBP levels in mice and humans. Cold exposure (24 hrs at 4°C) significantly increased circulating retinol and RBP concentrations in 129/Sv x C57BL/6J mice (Fig. 1a,b). Next, we tested if cold-induced alterations in vitamin A metabolism were also present in humans. 30 healthy lean subjects (baseline characteristics Table S1) were exposed to moderate cold (14-17°C) for 2.5 hrs using water-perfused cooling vests under a protocol which has been used successfully to activate BAT in humans (van Marken Lichtenbelt et al., 2009; Yoneshiro et al., 2013). Notably, only 2.5 hrs of moderate cold exposure increased circulating retinol and RBP concentrations significantly in these lean subjects (Fig. 1c,d).

**Fig. 1:**
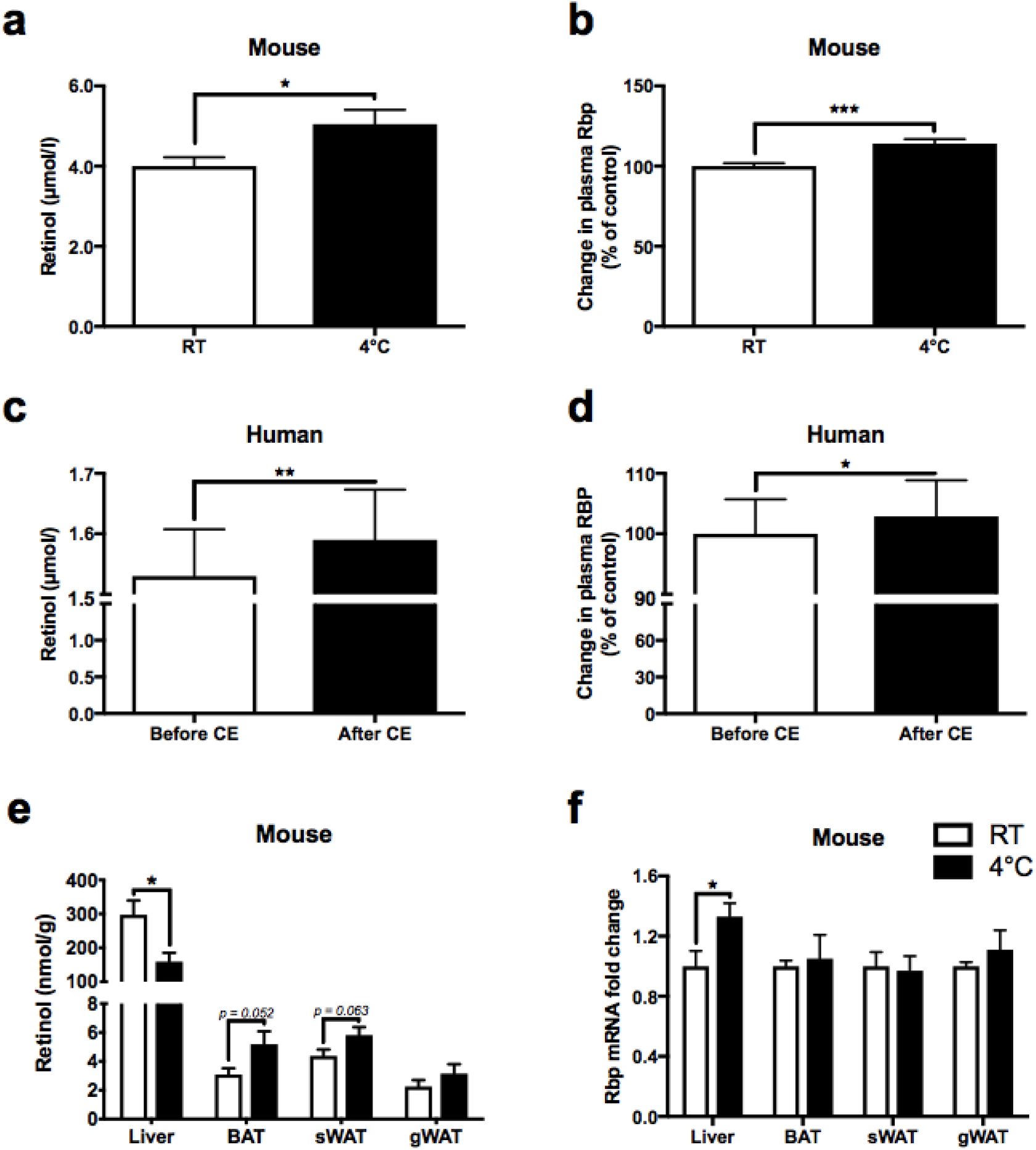
Cold exposure regulates systemic retinoid levels in mice and humans. Circulating retinol and Rbp concentrations in mice (n=9-10/group) housed at either room temperature or 4°C (a,b). Plasma retinol and RBP concentrations in humans (n=30) before and after cold exposure (Chondronikola et al.) for 2 hrs (c,d). Analysis of tissue retinol levels of mice (n=7-9/group) housed at either room temperature or 4°C (e). Rbp gene expression of WT mice (n=4-5/group) housed at RT or 4°C (f). Data are given as mean±SEM. * p≤0.05; ** p≤0.01; ***p≤0.001.

To identify the tissue-specific source of cold-mediated increase in plasma vitamin A, we analyzed liver, WAT and BAT retinol levels in mice. Hepatic retinol content was markedly decreased in response to cold exposure, whereas retinol levels increased in subcutaneous WAT (sWAT), and BAT but not gonadal WAT (gWAT) (Fig. 1e), strongly suggesting coordinated mobilization and redistribution of hepatic retinol to adipose tissue. In accordance with increased Rbp plasma concentrations, cold exposure induced *Rbp* mRNA expression significantly in the liver but not adipose depots (Fig. 1f).

These data establish that systemic vitamin A turnover, circulating plasma levels and ultimate adipose tissue distribution is tightly regulated by cold exposure in mice and possibly humans.

### Liver to fat retinoid redistribution is impaired in Rbp deficiency

To study the functional effects of disturbed cold-mediated retinoid redistribution, we used Rbp-deficient (Rbp^-/-^) mice, a model for impaired hepatic retinol mobilization and defective retinol transport. Rbp^-/-^ mice maintained on a vitamin A-rich diet do not manifest global vitamin A deficiency, however mobilization of hepatic retinoid stores is significantly impaired (Quadro et al., 1999). Both, WT and Rbp^-/-^ mice were maintained at room temperature (24°C) or exposed to cold (4°C) for 24 hrs. In contrast to WT mice, plasma retinol was barely detectable in Rbp^-/-^ mice, as expected (Quadro et al., 1999), and did not increase with cold exposure (Fig. 2a). Unlike WT controls, retinol concentrations in Rbp-deficient liver and sWAT were not affected by cold challenge (Fig. 2b,c). In contrast, Rbp deficiency did not prevent a cold-induced increase in retinol levels in BAT and gWAT (Fig. 2d,e). Together, these findings indicate that Rbp action is required for cold-induced retinol redistribution between liver and subcutaneous adipose tissue.

**Fig. 2:**
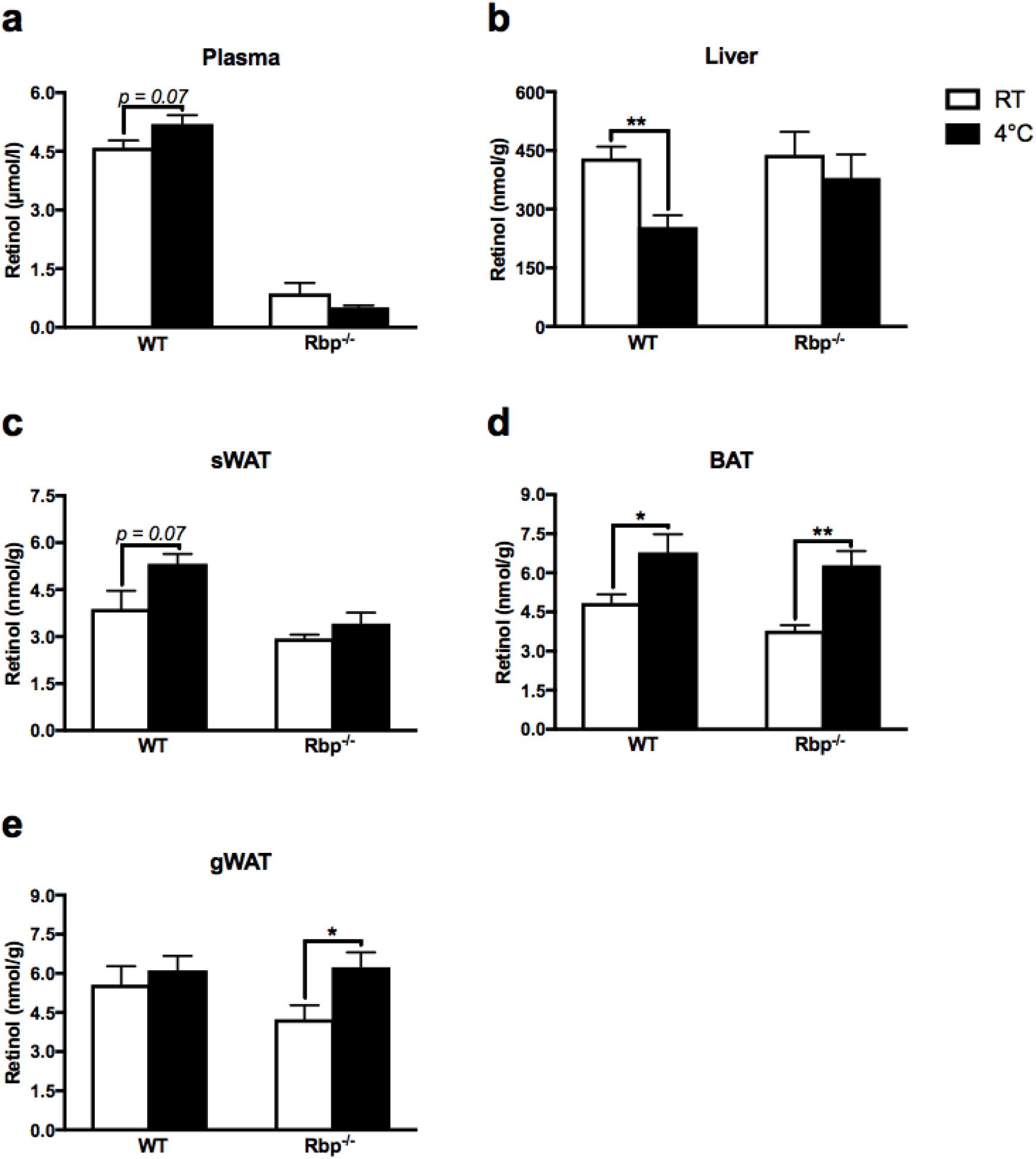
Liver to fat retinol redistribution is impaired in Rbp deficiency. Plasma retinol levels in WT and Rbp^-/-^ mice before and after cold exposure (**a**). Analysis of tissue retinol levels in WT and Rbp^-/-^ mice before and after cold exposure (**b-e**). Data are given as mean±SEM (n=4-5/group). * p≤0.05; ** p≤0.01.

### Intact retinol transport is vital for WAT browning

Using this Rbp^-/-^ mouse model provided a unique tool to study the effects of perturbed retinol delivery on cold-mediated WAT browning. Cold-induced expression of important thermogenic genes including *Ucp1, Cidea, Elovl3 and Pgc1α* was repressed in sWAT of Rbp^-/-^ as compared to WT mice (Fig. 3a) while Rbp deficiency had only modest or no effects on thermogenic gene expression in BAT (Fig. 3b). Interestingly, circulating retinol and Rbp correlated positively with Ucp1 expression in sWAT (Fig. 3c) and BAT (Fig. 3d), in keeping with a functional relationship between vitamin A levels and the thermogenic program. A hallmark of WAT browning is the emergence of cold-induced UCP1-positive multilocular beige adipocytes. Whereas beige adipocytes were abundantly induced in sWAT of cold-exposed WT mice, adipocyte morphology as well as UCP1 protein expression in Rbp^-/-^ sWAT were not affected by cold exposure (Fig. 3e,g). Despite blunted sWAT browning, Rbp deficiency did not interfere with the thermogenic program in BAT (Fig. 3b,f,h). Tissue norepinephrine (NE) concentrations did not differ between cold-challenged WT and Rbp^-/-^ mice (Fig. S1a) suggesting that altered sympathetic output did not contribute to this phenotype. These findings demonstrate that cold-mediated retinol flux towards white fat is critical for the induction of a browning program in sWAT.

**Fig. 3:**
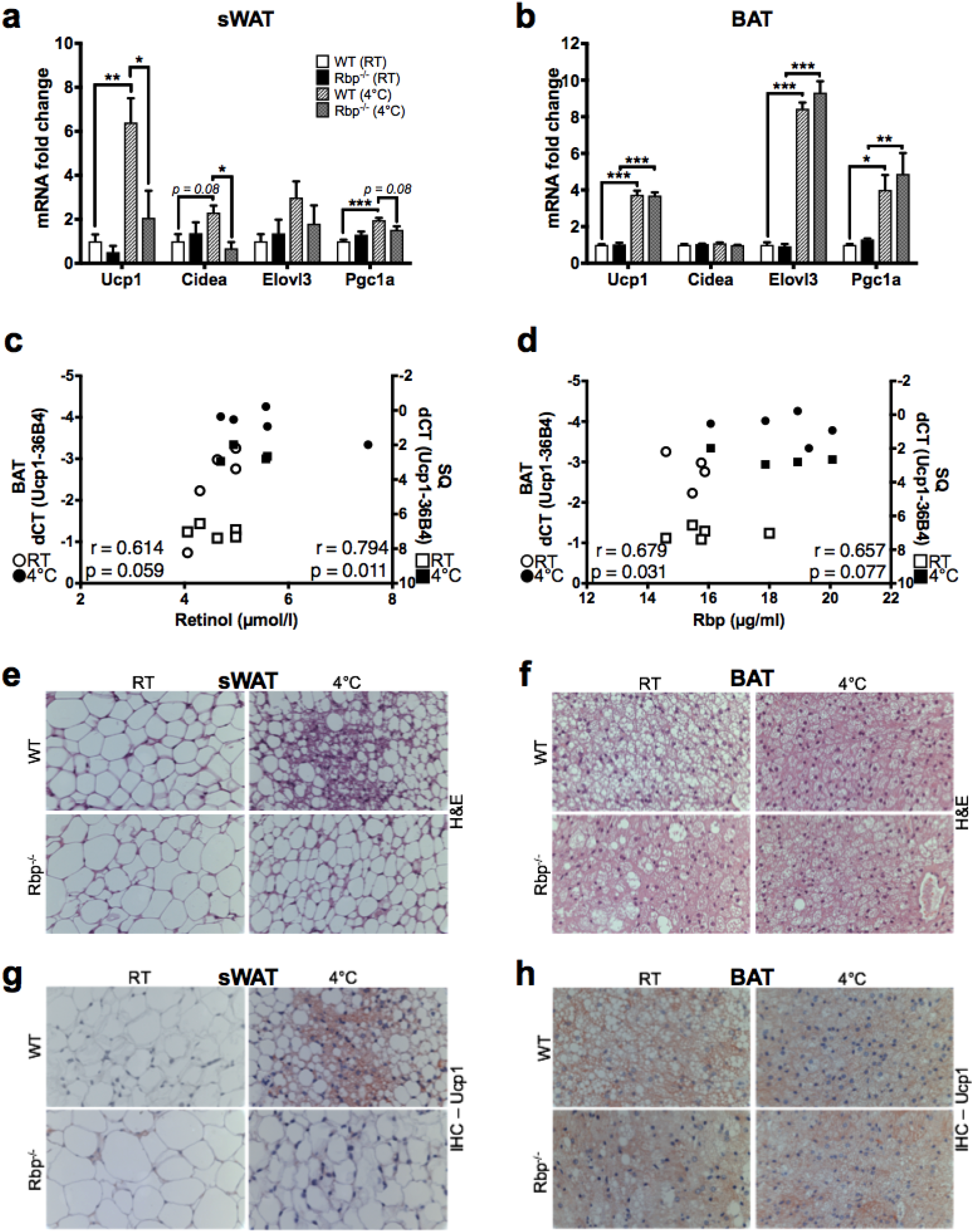
Intact retinol transport is vital for WAT browning. Expression of thermogenic genes in sWAT (a) and BAT (b) of WT and Rbp^-/-^ mice. Pearson’s correlation of circulating retinol (c) and Rbp (d) with Ucp1 mRNA expression in sWAT and BAT. Representative images of haematoxylin and eosin staining (H&E) (e, f) as well as Ucp1 immunohistochemistry (g, h) of sWAT and BAT from WT and Rbp^-/-^ mice exposed to RT or 4°C (n=5/group). * p≤0.05; ** p≤0.01; *** p≤0.001.

### Defective retinol transport perturbs the cold-induced TCA cycle and OXPHOS program in sWAT

Next, we studied the effects of Rbp deficiency on the genetic program of cold-regulated mitochondrial energy metabolism. Rbp deficiency repressed cold-induced induction of canonical participants in the tricarboxylic acid (TCA) cycle (*Cs, Aco2, Dlat, Sdha*) as well as the mitochondrial electron transport chain (ETC, *Cox4, Cox7a1, Cox8b*) in sWAT (Fig. 4a). In line with unaltered BAT thermogenesis, TCA and ETC gene expression was unaffected by Rbp deficiency in this depot (Fig. S1b). The transcriptional changes in Rbp-deficient sWAT were accompanied by lower protein levels of the mitochondrial ETC subunits I, II and V, the site for oxidative phosphorylation (OXPHOS) (Fig. 4b). The reduction of the OXPHOS protein complexes was not associated with decreased mitochondrial content since expression of the mitochondrial marker protein TOM20 was similar in WT and Rbp^-/-^ sWAT (Fig. 4c).

**Fig. 4:**
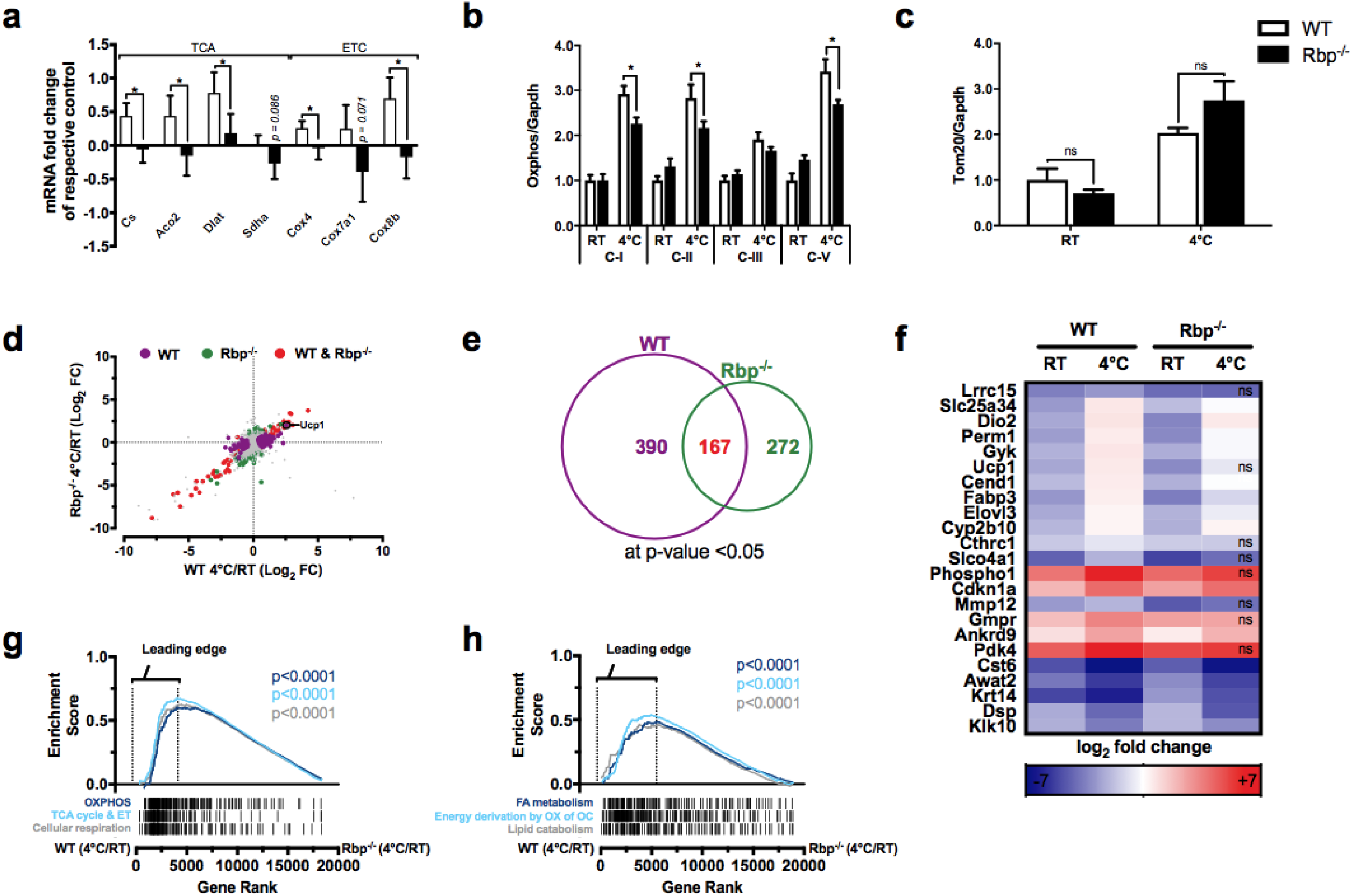
Defective retinol transport perturbs the cold-induced TCA cycle and OXPHOS program in sWAT. Gene expression of tricarboxylic acid (TCA) cycle genes and electron transport chain (ETC) genes (**a**). Protein expression of oxidative phosphorylation (OXPHOS) enzymes of the ETC (**b**) and TOM20 (**c**) in sWAT of WT and Rbp^-/-^ mice. mRNA sequencing data showing cold-induced changes in gene expression in sWAT of WT and Rbp^-/-^ mice. Purple, green and red spots represent genes significantly differentially expressed (p<0.05; n=5 mice per group). Grey dots indicate genes that are not statistically significantly regulated (**d**). Venn diagram illustrating the number of significantly cold-regulated genes in each genotype (**e**). Heatmap of the top 18 genes upregulated and the top 5 genes downregulated by cold exposure. Each column represents pooled data from sWAT of 5 mice (**f**). Gene set enrichment analysis (GSEA) of HALLMARK, GO and REACTOME gene sets in sWAT highlight browning pathways including “oxidative phosphorylation”, “TCA cycle and respiratory electron transport” and “cellular respiration” (**g**) as well as lipid metabolism including “fatty acid metabolism”, “energy derivation by oxidation of organic compounds” and “cellular lipid catabolic processes” (**h**). Gene expression is given as mRNA fold change as mean±SEM (n=5/group). * p≤0.05; ns = not significant.

To further characterize the molecular changes of impaired sWAT browning following cold exposure in the Rbp deficiency model, we performed unbiased mRNA sequencing (mRNAseq) analysis in sWAT. In WT mice, cold exposure significantly altered 390 genes, of which 318 were induced and 72 repressed (Fig. 4e, Table S2). In Rbp^-/-^ sWAT 272 genes were significantly regulated, of which only 72 were induced and 200 were repressed (Fig. 4e, Table S3). mRNAseq analysis also confirmed previous qPCR results showing the top regulated genes in sWAT of cold-exposed WT mice were thermogenic markers *Dio2, Elovl3 and Ucp1* among others (Fig. 4f). In contrast, many thermogenic genes including *Ucp1* were not significantly induced in cold-exposed Rbp^-/-^ mice, in line with our previous findings (Fig. 3a, 4f). Gene set enrichment analysis (GSEA) of Gene Ontology (GO), Reactome and Hallmark genes sets revealed marked induction of browning pathways involving *“oxidative phosphorylation”, “TCA cycle and respiratory electron transport”* and *“cellular respiration”* in cold-exposed WT versus Rbp ^-/-^ sWAT (Fig. 4g, Table S4). We also identified a strong shift towards increased cold-induced lipid metabolism in WT versus Rbp deficient sWAT, with significant gene set enrichment of *“fatty acid metabolism”, “energy derivation by oxidation of organic compounds”* and *“cellular lipid catabolic processes”* (Fig. 4h, Table S5). In summary, these data suggest that the absence of a cold-induced retinol increase in sWAT mitigates the browning capacity while also causing a qualitative defect in mitochondrial respiratory function and concordant changes in lipid metabolism.

### Rbp deficiency impairs cold-induced lipid turnover and *in vivo* thermogenesis

During cold adaption, fatty acids from plasma triglycerides (TGs) are essential energy substrates utilized for Ucp1 activation as well as mitochondrial ß-oxidation. In BAT, cold exposure accelerates plasma TG clearance through increased lipoprotein lipase activity and uptake into brown adipocytes (Broeders et al., 2015). Given the marked shift in lipid metabolism and fatty acid oxidation gene expression sets in sWAT of Rbp^-/-^ vs WT mice (Fig. 4h), we analyzed plasma TG and non-esterified fatty acids (NEFA) clearance as well as changes in cholesterol levels after cold exposure. Cold-induced triglyceride clearance was significantly decreased in Rbp^-/-^ mice (Fig. 5a) while NEFA clearance did not differ between the two genotypes (Fig. 5b). Total cholesterol levels were similarly but not-significantly increased in both genotypes following cold exposure (Fig. 5c). However, circulating triglyceride-rich VLDL-cholesterol levels were markedly reduced by cold exposure in WT but not in Rbp-deficient mice (Fig. 5d,e).

**Fig. 5:**
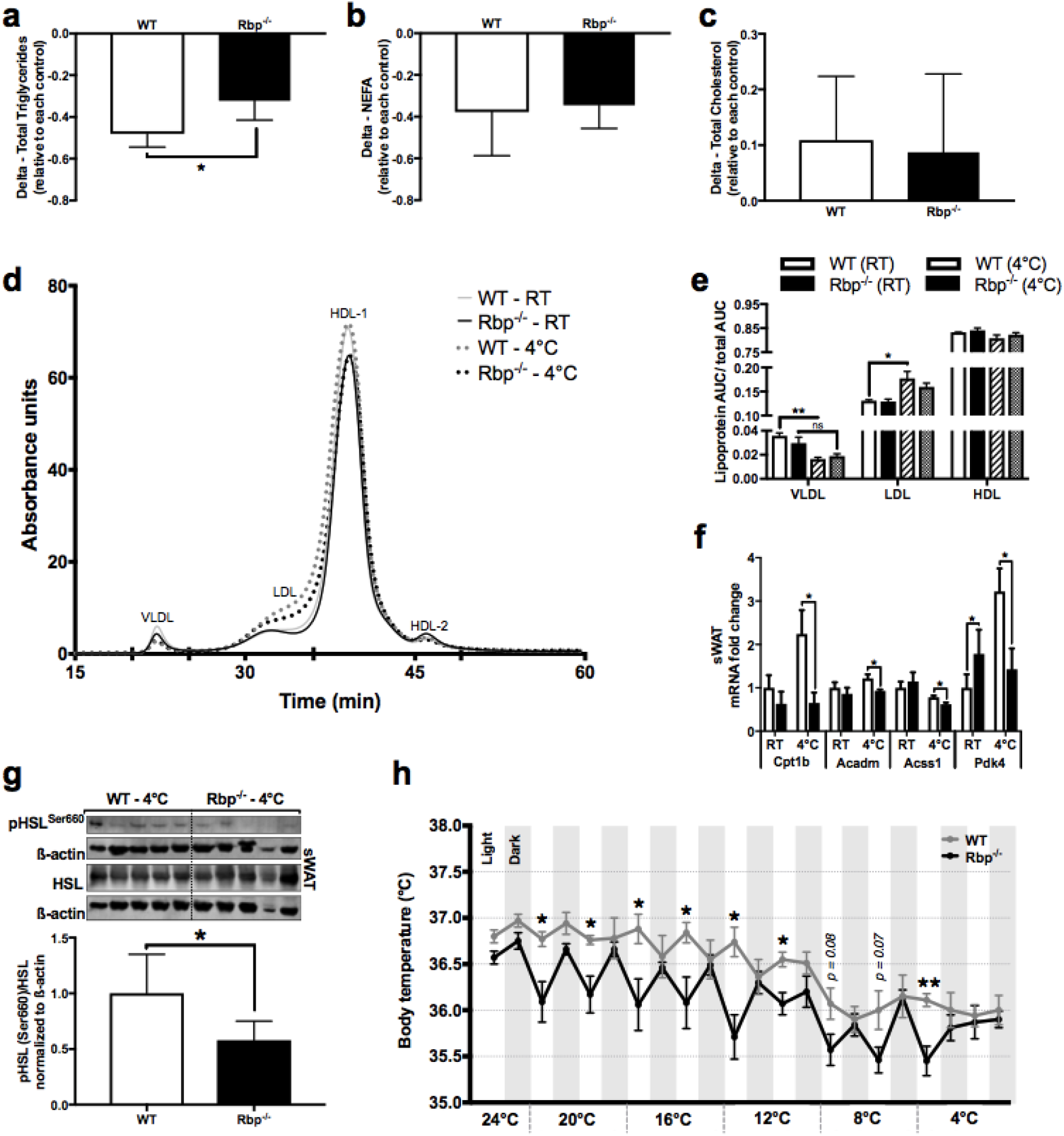
Rbp deficiency impairs cold-induced lipid turnover and in vivo thermogenesis. Cold-induced changes of circulating total triglycerides, NEFAs and total cholesterol (Calderon-Dominguez et al.). Plasma cholesterol distribution in WT and Rbp^-/-^ mice assessed by FPLC analysis in mice housed at RT or 4°C (**d**). Specific lipoprotein area under the curve (AUC) was normalized to total AUC of each group (**e**). mRNA fold change of fatty acid (Chondronikola et al.) transport and FA oxidation markers (**f**). Western blot analysis of pHSL/HSL expression in sWAT (**g**). Telemetric core body temperature measurements in WT and Rbp^-/-^ mice during a stepwise cooling challenge (**h**). n=5/group, * p≤0.05; ** p≤0.01; ns = not significant.

Next, we analyzed expression of genes involved in mitochondrial fatty acid transport and ß-oxidation in sWAT of WT and Rbp^-/-^ mice. Expression of several rate-limiting FA transporters and ß-oxidation enzymes including carnitine palmitoyltransferase 1b (*Cpt1b*), medium-chain acyl-CoA dehydrogenase (*Acadm*), acyl-CoA synthetase short chain family member 1 (*Acss1*) and pyruvate dehydrogenase kinase 4 (*Pdk4*) were significantly lower in cold-challenged Rbp^-/-^ vs WT sWAT (Fig. 5f). Minimal if any differences were observed in cold-exposed BAT between the two genotypes (Fig. S1c). Finally, we examined hormone sensitive lipase (HSL) activity, the rate limiting step in lipolysis. Ser660 phosphorylation of HSL was significantly decreased in sWAT of cooled Rbp^-/-^ compared to WT mice (Fig. 5g) in line with reduced cold-induced lipolytic activity. Together this data point towards impaired local and systemic cold-regulated lipid homeostasis in the absence of Rbp.

To test consequences of these profound molecular changes observed in Rbp deficient sWAT *in vivo*, we performed core body temperature measurements in cold-challenged animals. In line with impaired sWAT browning and perturbed lipid oxidation, Rbp^-/-^ mice were significantly more cold-sensitive as reflected by lower core body temperatures during cold stress (Fig. 5h).

### Retinol enhances oxidative capacity in primary human adipocytes and is associated with increased lipid utilization in humans

In order to study the effects of retinol on the thermogenic capacity in a cell autonomous manner we used human adipocyte precursor cells (hAPCs) isolated from abdominal subcutaneous fat specimens. hAPCs were differentiated for five days before stimulation with various retinol concentrations (1 nM to 10 μM) or vehicle for 24 hrs. 1 μM retinol resulted in the most robust increase in UCP1 mRNA expression (Fig. S1d) and was therefore used for all other experiments. Retinol stimulation significantly increased thermogenic gene expression and UCP1 protein levels in differentiated hAPCs from four different donors (healthy females, aged 35-45 years (Fig. 6a,b). Notably, retinol stimulation enhanced cellular respiration as demonstrated by increased maximum, mitochondrial and reserve capacity oxidative consumption rate using *Seahorse*^®^ analysis (Fig. 6c,d). Hence, these data suggest that retinol promotes browning in human adipocytes with coordinated increases in oxidative metabolism. In line with this finding, short-term cold exposure in 30 healthy lean subjects concomitantly increased circulating retinol concentrations (Fig. 1c) with concurrent significant decreased respiratory quotients (RQ, defined as the ratio between CO_2_ release and O_2_ consumption) by 6.0%±1.87% (Fig. 6e). A significant inverse correlation existed between the cold-mediated change in plasma retinol (delta retinol) and the change in the respiratory quotient (delta RQ) (Fig. 6f), suggesting that enhanced cold-induced retinol flux is associated with augmented lipid utilization in humans.

**Fig. 6:**
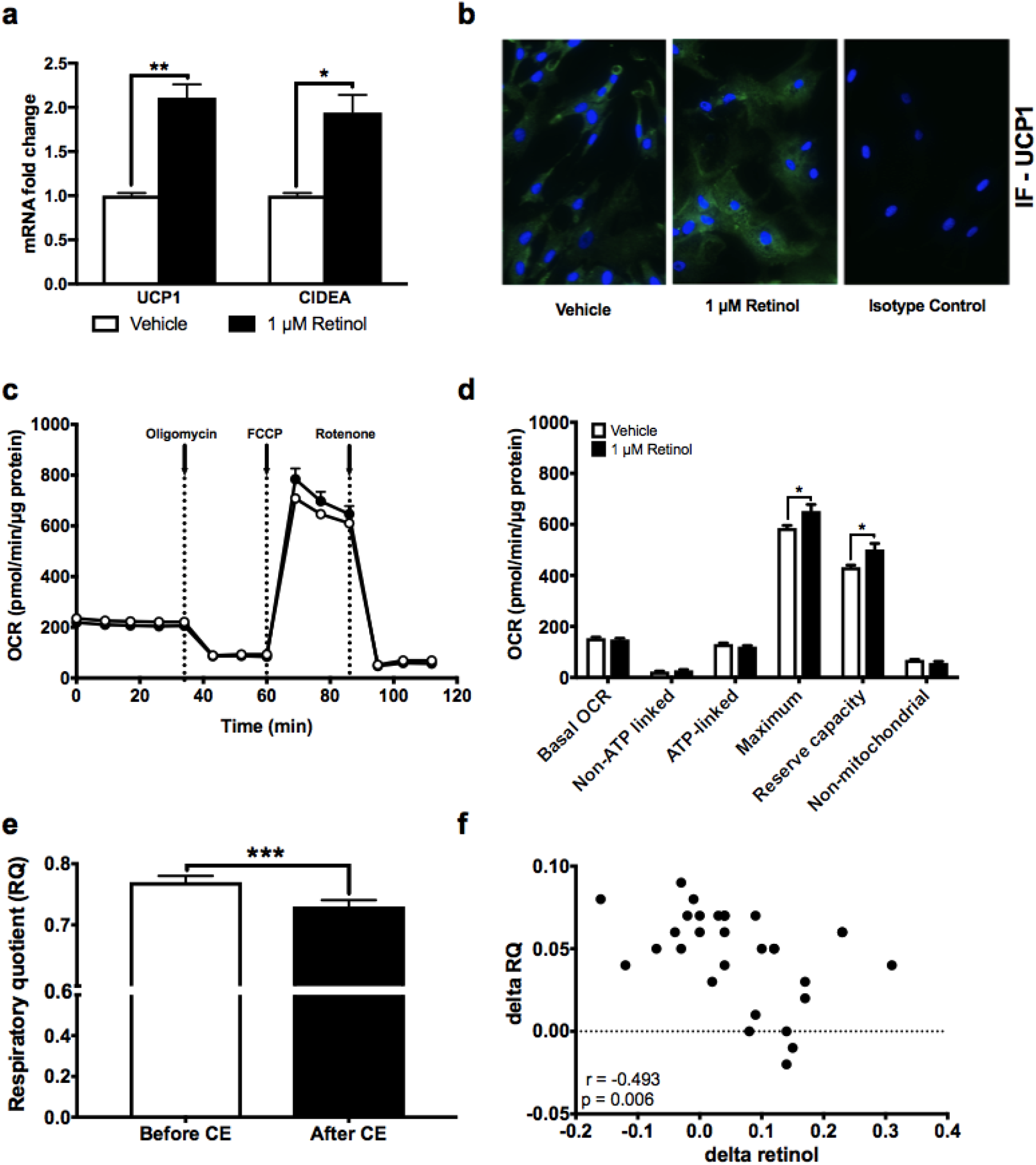
Retinol enhances oxidative capacity in primary human adipocytes. UCP1 gene expression (**a**) and immunofluorescence (**b**) in differentiated primary human adipocytes stimulated with retinol for 24 hrs (n=5). Oxygen consumption rate (OCR) of primary human adipocytes treated with 1 μM retinol for 24 hrs. Data are illustrated as real time replicate readings (**c**) or as the group average of different respiratory phases (basal, non-ATP linked, ATP-linked, maximum, reserve capacity, non-mitochondrial and mitochondrial respiration) (**d**). Respiratory quotient (RQ) assessed by indirect calorimetry in 30 healthy lean subjects before and after 2.5 hrs of cold exposure (**e**). Spearman’s correlation of cold-induced retinol changes with RQ changes in healthy lean humans (**f**). Values are given as mean ± SEM. * p≤0.05, ** p≤0.01, *** p≤0.001.

## DISCUSSION

Thermogenesis is an essential and at times adaptive biologic function required for survival among endothermic species that must generate heat internally. BAT, by uncoupling electron generation and storage, offers a source for this energy production (Cannon & Nedergaard, 2004). Given excess energy intake as a significant contributor to the epidemic obesity, the prospect of inducing WAT to take on the energy wasting characteristics of BAT has received considerable attention (Kiefer, Orasanu, et al., 2012; Seale et al., 2011; Vegiopoulos et al., 2010). Extensive prior work establishes retinoid signaling as a key regulator of adipogenesis, UCP1 expression and overall energy balance (Kiefer, Vernochet, et al., 2012; Mercader et al., 2006; Villarroya et al., 2004). Despite this, specific connections between the liver, as the primary site of retinoid formation and storage, retinol release, circulating plasma levels and delivery to tissues like fat are lacking. We establish here a novel liver-to-fat axis that controls WAT browning via integrated hepatic retinol release and Rbp-mediated delivery to distant adipose beds, with evident transcriptional coordination of TG metabolism, fatty acid oxidation and thermogenic mediators.

Retinoids exert most of their metabolic effects within liver and adipose tissue (Frey & Vogel, 2011; Ribot et al., 2004; Ross, 1993; Ziouzenkova & Plutzky, 2008). Global vitamin A deficiency in animals is linked to increased body weight and BAT adiposity (whitening of brown fat), whereas retinoic acid or retinaldehyde induce thermogenic gene expression in adipocytes and reduce fat stores (Alvarez et al., 1995; Berry et al., 2012; Berry & Noy, 2009; Bonet et al., 2000; Kiefer, Vernochet, et al., 2012; Mercader et al., 2006; Puigserver et al., 1996; Ribot et al., 2001, 2004; Ziouzenkova et al., 2007). Despite accumulating evidence implicating retinoids in adipocyte thermogenesis, fundamental issues remained unresolved, including a) the dependency on retinoids for cold-mediated adaptive thermogenesis, b) the physiologic regulation of tissue-specific retinoid availability in response to cold or c) the contribution of hepatic retinoid stores to browning of WAT or increased BAT activity. We demonstrate here striking cold-mediated shifts in retinol between liver, BAT, sWAT and plasma accompanied by increased circulating levels of Rbp as the primary retinol carrier.

This cold mediated retinoid flux from the liver towards adipose tissue evident in WT mice was absent in Rbp deficient mice, a model with defective hepatic retinoid mobilization and systemic transport (Quadro et al., 1999). These data argue that cold exposure stimulates mobilization of hepatic retinoid stores via Rbp (D’Ambrosio, Clugston, & Blaner, 2011; Shirakami, Lee, Clugston, & Blaner, 2012), thus shuttling retinol towards extrahepatic tissues. These cold mediated shifts also induce coordinated changes in gene expression programs involved in thermogenesis, TG metabolism and fatty acid beta oxidation. Thus, the extensive, primary vitamin A storage in the liver (Blaner et al., 2009) combines with our data to identify the liver as a massive but regulated thermogenic reservoir signaling to fat depots via retinoid release, as evident in cold adaption (Fig. 7).

**Fig. 7:**
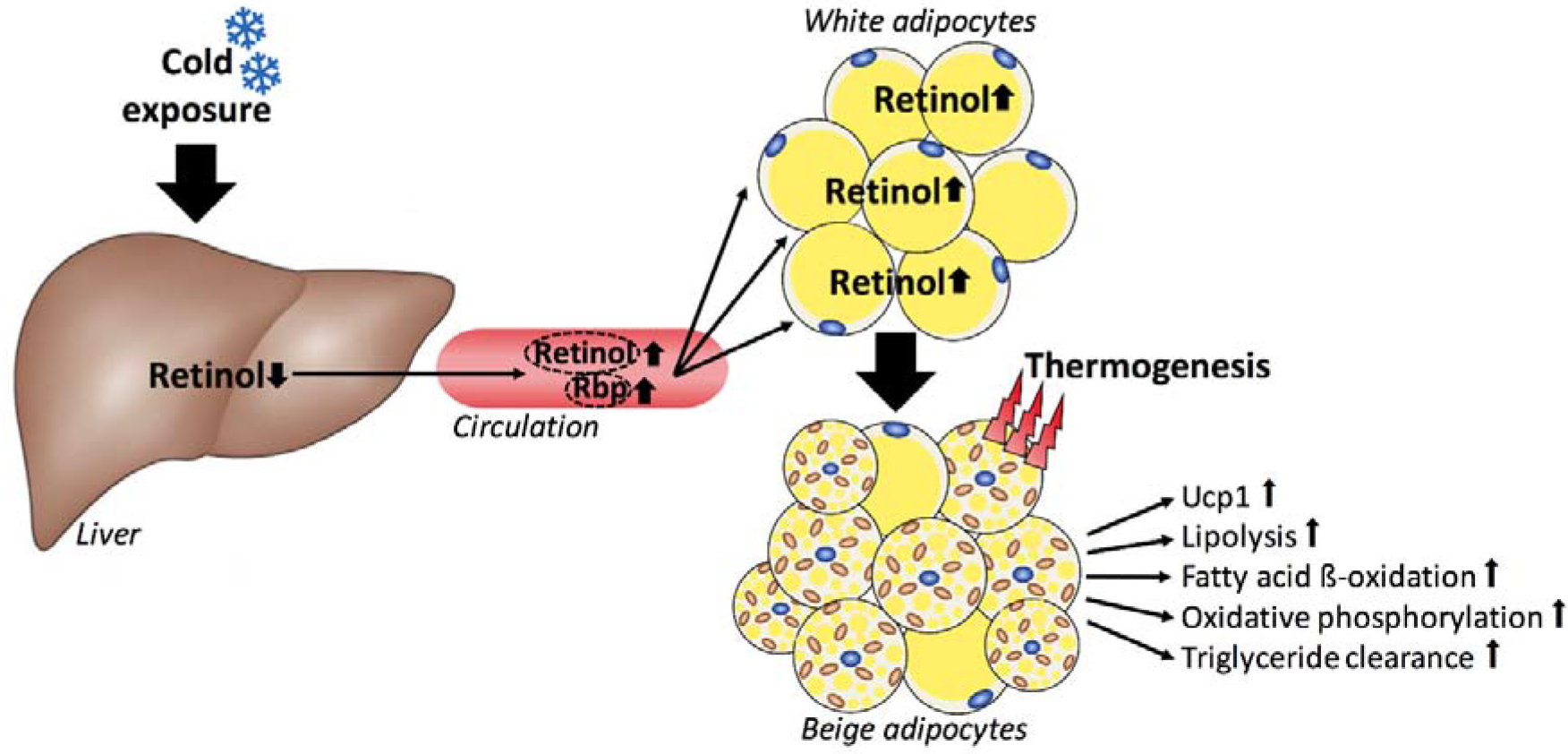
Graphical abstract. Highlights:

- Systemic vitamin A abundance is tightly regulated by cold exposure
- Cold-induced redistribution of vitamin A from liver towards white adipose tissue is critical for browning and mitochondrial function
- Defective retinol transport to target tissues impairs cold-induced thermogenesis
- Retinol mediates previously unknown interorgan crosstalk between liver and adipose tissue with important implications for global substrate metabolism

Rbp is known to be a key player in energy balance, including diabetes. Our data identifies Rbp as essential for executing the liver-fat axis in energy balance and thermogenesis. Prior studies found an association between increased circulating Rbp levels and insulin resistance in various genetic and diet-induced models of obesity (Norseen et al., 2012; Yang et al., 2005). Human studies have been more controversial with variable relationships between Rbp concentrations and different cohorts with obesity, insulin resistance or type 2 diabetes (Korek et al., 2018; Kotnik, Fischer-Posovszky, & Wabitsch, 2011; Mody, 2017), which could reflect dietary or genetic differences. DNA-arrays in the insulin resistant adipose-specific glucose transporter type 4 (GLUT4) knockout mice revealed elevated Rbp expression levels in adipose tissue but unaltered Rbp expression in the liver (Yang et al., 2005). Thus, elevated circulating Rbp levels in insulin resistant states were attributed to increased expression and secretion by adipose tissue (Yang et al., 2005). In our study, cold exposure increased Rbp mRNA expression in the liver accompanied with elevated circulating Rbp concentrations with no changes in adipose depots. In general, cold exposure improves insulin sensitivity and glucose homeostasis in mice and humans (Bartelt et al., 2011; Chondronikola et al., 2014; Hanssen et al., 2015), suggesting that increased circulating Rbp levels after cold stimulation do not cause insulin resistance. Recent evidence suggests that the Rbp tissue source may determine insulin resistance. Mice with hepatocyte-specific deletion of Rbp (LRKO) exhibit no detectable plasma Rbp levels on chow or high-fat diet despite intact adipose Rbp production, establishing hepatocytes as the principal source of circulating Rbp. More importantly, LRKO mice are not protected from diet-induced obesity or insulin resistance consistent with adipocyte-secreted Rbp is confined to autocrine or paracrine actions within adipose tissue, even in an insulin resistant state (Fedders et al., 2018; Thompson et al., 2017). In support of this notion, adipocyte-specific Rbp overexpression increases adipose tissue inflammation, lipolysis and circulating FA levels (Lee, Yuen, Jiang, Kahn, & Blaner, 2016). These findings may also explain why cold-mediated increases in systemic Rbp levels do not cause insulin resistance.

Rbp is well established as a key protein in retinoid metabolism (Quadro, Hamberger, Colantuoni, Gottesman, & Blaner, 2003; Soprano D. R., 1994) mediating retinol secretion and extrahepatic transport preferentially towards adipose tissue as previously demonstrated in Rbp^-/-^ mice challenged with dietary conjugated linoleic acid supplementation or chronic alcohol consumption (Clugston, Huang, & Blaner, 2015; Ortiz et al., 2009). Both studies showed that Rbp facilitates hepatic retinol redistribution towards fat depots although some residual adipose tissue retinol uptake may persist in Rbp deficiency (Clugston et al., 2015) as we observed in BAT of cold-challenged Rbp^-/-^ mice (Fig. 2e). In contrast to BAT, cold-mediated retinol increase was completely abrogated in sWAT of Rbp-deficient mice (Fig. 2d). These fat depot specific differences may stem from an alternative retinol uptake mechanism in BAT. Postprandial tissue retinol supply is enabled by lipoprotein lipase (LPL)-dependent hydrolysis of retinol-containing chylomicrons released by enterocytes after dietary intake (Blaner et al., 1994; Blomhoff, Green, Berg, & Norum, 1990). Retinoid provision to BAT during cold challenge via this mechanism, even in the absence of Rbp would be consistent with cold-mediated induction of LPL expression in both WT and Rbp^-/-^ BAT (data not shown). We show here that the induction of a thermogenic program in sWAT strongly depends on intact Rbp-mediated retinoid transport and tissue availability. Despite intact BAT function, blunted adipose tissue browning in Rbp deficiency has systemic consequences including impaired cold tolerance and perturbed triglyceride clearance. Defective BAT retinoid supplies may further aggravate this thermogenic phenotype. In fact, disrupted thermoregulation has previously been reported in alcohol-challenged mice with decreased BAT retinoic acid content (Clugston et al., 2015).

Given the importance of systemic energy balance and thermogenesis, a coordinated interorgan crosstalk that allows for integrated networks would be expected. Recently, several liver-derived factors have been identified that contribute to BAT activation or WAT browning including fibroblast growth factor 21 (Douris et al., 2015; Fisher et al., 2012), bile acids (Broeders et al., 2015; Worthmann et al., 2017) or acylcarnitines (Simcox et al., 2017). Our results establish hepatic retinoids as novel mediators of extended liver to adipose signaling. Control over thermogenesis involves regulation of sources of energy substrates through lipolysis and FA release as well as combustion of those resources through beta oxidation. In keeping with this, blunted WAT browning in Rbp deficiency is coupled to repressed expression of lipid catabolism and FA oxidation genes, with decreased mitochondrial oxidative phosphorylation capacity. Consequently, cold-induced TG clearance was impaired in Rbp^-/-^ mice. The importance of activated BAT for TG hydrolysis has been established in several mouse models and in human cold exposure studies (Bartelt et al., 2011; Berbee et al., 2015; Khedoe et al., 2015). We show here that the cold-induced increase of circulating retinol occurs in conjunction with a decreased respiratory quotient in humans (Fig. 6f), suggesting that enhanced retinol supply is associated with higher lipid oxidation. Indeed, retinol stimulation in primary human adipocytes augmented the mitochondrial oxidative capacity (Fig. 6c, d), suggesting that the retinol effects are cell-autonomous. Given that retinol is intracellularly converted to retinoic acid, which activates the nuclear receptors RAR and RXR, increased retinoic acid signaling may contribute to the observed thermogenic effects (Fig. 6a, as widely reported (Alvarez et al., 2000; Alvarez et al., 1995; Berry et al., 2012; Mercader et al., 2010; Mercader et al., 2006; Puigserver et al., 1996; Rabelo et al., 1996)). Accordingly, retinoic acid target gene expression was markedly induced in retinol stimulated human adipocytes (Fig. S1d).

Taken together, we demonstrate here that local hepatic retinoid stores and the systemic delivery of these molecules via hepatic Rbp are essential for cold-induced thermogenic responses in adipose tissue. Moreover, these data establish coordinated regulation of hepatic retinol shuttling to WAT by Rbp as essential for normal adaption to cold by enabling a transcriptional thermogenic program (summarized in Fig. 7). The evidence provided here for the presence of this liver-fat retinoid signaling in both mice and humans identifies novel mediators and pathways exerting control over energy balance and thermogenesis, with potential for new insights to obesity and its treatment.

## MATERIAL AND METHODS

### Animals

Rbp knockout mice (Rbp^-/-^) as well as their littermate controls (wildtype, WT) were kindly provided by Dr. Loredana Quadro (Rutgers University) and bred on a mixed background (129/Sv x C57BL/6J) (Quadro et al., 1999). Mice were kept on a standard chow diet with a vitamin A content of 25 IU g^-1^. All experiments were approved by the local ethics committee for animal studies and the Federal Ministry of Science, Research and Economy (BMBWF-66.009/0124-V/3b/2018) and followed the guidelines on accommodation and care of animals formulated by the European Convention for the Protection of Vertebrate Animals Used for experimental and other scientific purposes.

### Human cold exposure studies

Moderate cold exposure (14°C-17°C) for two hours is usually sufficient to activate human BAT (van Marken Lichtenbelt et al., 2009; Yoneshiro et al., 2013). We applied this protocol at 30 young healthy lean subjects (age: 20-45 years, BMI: 18.5 – 24.9 kg/m2) which was approved by the ethics committee of the Medical University of Vienna (EK-No. 1032/2013), registered at CinicalTrials.gov (NCT02381483), and performed in compliance with the Declaration of Helsinki and Good Clinical Practice. In detail, participants were examined in the morning after an overnight fasting period. A venous catheter was placed in order to perform blood sampling and after a 30 minutes resting phase an indirect calorimetry (Quark RMR, Cosmed, Italy) was performed. Subjects were then fitted with a water perfused cooling vest with adjustable temperature (CoolShirt, Stockbridge, Georgia, USA) and participants were cooled slightly above muscle shivering in order to avoid shivering thermogenesis. Electromyography (OT Bioelettronica, Turin, Italy) was used to detect subclinical shivering and thus allowed us to maintain the proper cooling temperature. After 2.5 hrs cold exposure another indirect calorimetry and blood sampling were performed. Serum vitamin A and RBP were determined by HPLC (Chromsystems Instruments & Chemicals GmbH) and nephelometry (Siemens BnII), respectively, by the Department of Laboratory Medicine, Medical University of Vienna using commercially available kits.

### Cold exposure and body temperature measurements in mice

To determine the vitamin A abundance as a critical determinant of adaptive thermogenesis 16-18 weeks old male body weight matched WT and Rbp^-/-^ mice were single housed, without cage enrichment, and cold exposed for 24 hrs in a climate chamber for keeping mice HPP750life (Memmert). In the stepwise cooling experiment, telemetric temperature probes (Anipill Capsule 0.1C Precision) were implanted intraperitoneally to control real time body temperature. After a 1 week acclimation phase, ambient temperature was dropped by 4°C every 48 hrs. Mice were fasted 2 hours prior to killing, EDTA-plasma was collected by heart puncture. The liver was perfused to exclude circulating retinol from the liver pool. Thus, 0.5 U heparin was injected in the inferior vena cava while the portal vein and the inferior vena cava were cannulated and perfused with 0.9% NaCl and tissues were dissected and analyzed.

### Reverse transcription and gene expression

Total RNA was extracted using TRIzol (Invitrogen), treated with DNase (Thermo Scientific) and reverse transcribed to cDNA (Applied Biosystems) according to manufacturer’s instructions. Gene expression, normalized to 36B4, was analyzed by quantitative real-time RT-PCR (SybrGreen, Roche or GoTaq, Promega) using a QuantStudio 6 cycler (Applied Biosystems). Primer sequences are available upon request.

### Immunoblotting

Tissue samples were homogenized in RIPA buffer supplemented with protease and phosphatase inhibitors (Roche). Protein concentration was determined with the PierceTM BCA protein assay kit (23225, Thermo Scientific). Standard western blotting was performed using mouse or rabbit polyclonal antibodies to Oxphos (Abcam), Tom20 (Sigma), Gapdh (Cell Signaling), ß-actin (Novus Biologicals). Blots were visualized using either chemiluminescence (Roche) and quantified on Fusion Fx Vilber Lourmat (FusionFx, PeqLab) or by using fluorescent secondary antibodies and visualization with the Odyssey imaging system (Licor). Protein bands were quantified using densitometry in ImageJ.

### Immunohistochemistry

Paraffin sections were prepared from mouse sWAT and BAT after fixation (4.5% (vol/vol) formaldehyde). Sections were stained using haematoxylin and eosin (H&E) and with rabbit polyclonal antibody to Ucp1 (U6382, Sigma, 1:1000) and biotinylated secondary goat antibody to rabbit (Vector Laboratories Inc., 1:300). Control staining was performed on selected sections with isotype control. Counterstaining was performed with haemalaun for 1 min. Samples were analyzed on an Axio Imager 2.

### HPLC analysis

Mouse BAT samples were homogenized in ddH_2_O at a ratio of 18 mg to 120 μl and liver samples at a ratio of 15 mg to 150 μl ddH2O. From the lysates 100 μl were used for extraction. For the determination of gonadal or subcutaneous tissue retinol content a minimum of 50 mg whole tissue was used for analysis. Plasma retinol was determined in 100 μl EDTA-Plasma. All samples were spiked with 100 μl internal standard. Retinyl acetate (Cat.No 46958, Sigma) was prepared at a stock concentration of 1.5 mM in methanol including 500 nM BHT (Cat.No W218405, Sigma) which resulted in a final concentration of 150 picomoles of internal standard per sample. Retinoids were extracted using a double hexane (including 500 nM BHT) method. The aqueous phase was evaporated using a cold trap SpeedVac concentrator. Samples were stored at −20°C and re-suspended in 200 μl methanol including 500 nM BHT in amber vials prior to injection. A user defined injection program was applied. Loading solvent (= mobile phase A) was used for flushing of the injection needle or the buffer loop. The autosampler was cooled to 10°C and samples were placed in the closed compartment only immediately prior to injection. Retinol was resolved by reverse phase chromatography using the Phenomenex Kinetex Biphenyl column (2.1mm ID x 150mm length, 2.6μm particle size, 100 Å pore size) on an UltiMate 3000 HPLC System (ThermoFisher, Vienna, Austria) consisting of a quaternary gradient pump, autosampler, thermal compartment, and an UV detector operated at 325nm. Analytes were separated using the following gradient protocol (flow, 150 μl/min; gradient, 0-9 min 40 % A and 60% B, 10-20 0 % A and 100 % B, 21-35 40 % A and 60% B). Mobile phase A 5 % acetonitrile, 95% water with 0.1% formic acid and mobile phase B 5% methanol, 95% acetonitrile with 0.1% formic acid. The separation column was operated at 60°C in the thermal compartment. The peak signals detected by UV were integrated using the Chromeleon V6.8 (Thermo Scientific, Vienna, Austria) and the obtained peak area of retinol was used for calculation of retinol’s concentration using the equation of the standard curve (Retinol, P/N 95144, Sigma) spanning the physiological range of the analyte. Finally, the retinol content was normalized to mg tissue used for analysis.

### Determination of circulating parameter

ELISA and colorimetric assays were performed according to manufacturer’s protocols using EDTA-plasma for the measurement of Rbp4 (MRBP40, R&D Systems), Triglyceride (TR0100, Sigma), NEFA (Standard: 270-77000, NEFA HRII R1: 434-91795, NEFA HR II R2: 436-91995, Wako Diagnostics) and Total Cholesterol (MAK043, Sigma). Lipoprotein distribution was determined with fast protein liquid chromatography (FPLC) measurement which was performed by Audric Moses (Lipidomics Core Facility, The Women and Children’s Health Research Institute and the Faculty of Medicine and Dentistry, University of Alberta, Canada).

### Norepinephrine (NE) measurement

A commercial available ELISA Kit (BA E-52000, LND) was modified and established for the use of determining NE in tissue homogenates.

### Gene expression profiling and GSEA

Sequencing libraries were prepared using the NEBnext Ultra II directional RNA library prep kit for Illumina according to manufacturer’s protocols (NEB) and sequenced in the 75bp single-read mode on the Illumina NextSeq 500 (Illumina) at the Core Facility Genomics, Medical University of Vienna. RNA-Seq data were processed with the Illumina Cufflinks Assembly & DE workflow implemented in BaseSpace. More than 98.5% of 26-30 Mio reads per sample were mapped to the Mus musculus/mmc10 assembly of the murine genome using the STAR Aligner (Dobin et al., 2013). Differential gene expression was analyzed using Cuffdiff (Trapnell et al., 2013).

GSEA (Gene Set Enrichment Analysis) is a computational pathway analysis tool that determines if a given set of manually curated genes show statistically significant, concordant differences between two biological states (http://www.broadinstitute.org/gsea/index.jsp).The analysis was performed with the GSEA Software (Subramanian et al., 2005) (version 2.1.0) using the c2 (version 5) gene set database. Genes were first ranked based on real value using the weighted signal-to noise metric. *p*-values and false discovery rates (FDR) for the enrichment score of each gene set were calculated based on 1000 gene set permutations.

### Isolation of human adipocyte precursor cells (hAPCs) and in vitro adipocyte differentiation

Cells were obtained from five healthy female donors, aged 31-45 years undergoing abdominoplasty. Briefly, adipose tissue was thoroughly minced with scissors using sterile techniques, and digested using 20 mg collagenase II (Worthington Biochemical Corporation) in a shaking water bath at 37°C. Cells were filtered through a 100 μM nylon filter (Greiner Bio-One GmbH) and centrifuged (200 g, 5 min). The floating section was discarded and the stromal-vascular fraction pellet was re-suspended in growth medium and centrifuged again (200 g, 5 min). Thereafter, the pellet was re-suspended in growth medium and filtered using a 30 μM nylon mesh (Sysmex Europe GmbH/Sysmex Partec GmbH). Filtered hAPCs were cultivated in growth medium at 37°C with 5 % CO2 till they reached 80 % confluency. At this point, cells were trypsinized, counted and seeded at a density of 150000 cells/well in a 24-well plate or 80000 cell/well in XF 24-well cell culture microplates in growth medium (DMEM/F12 (1:1), 10% (vol/vol) FBS and 1% (vol/vol) penicillin-streptomycin, 33 μM Biotin, 17 μM D-pantothenic acid). Once cells reached confluency, induction medium (growth medium, 0.5 mM isobutylmethylxanthine, 850 nM insulin, 2 nM T3, 1 μM dexamethasone, 5 μM rosiglitazone) was added for 2 days. Thereafter, the medium was changed to differentiation medium (growth medium, 850 nM insulin, 2 nM T3, 5 μM rosiglitazone) for another 4 days while changed every 48 hrs. On day 5 cells were stimulated with 1 μM Retinol (Sigma), whereas 0.1 % EtOH served as vehicle, for 24 hrs. This study was approved by the Medical University of Vienna’s ethics committee and the General Hospital of Vienna (EK 1032/2013).

### Immunofluorescence

Cells cultured in chamber slides were fixed in 2 % (vol/vol) paraformaldehyde, permeabilized in 1% Triton X (30 min) and blocked in 3 % BSA-PBS. UCP1 staining was performed over night with a rabbit polyclonal antibody against UCP1 (U6382 Sigma Aldrich, 1:1000). UCP1 was visualized with a fluorescence secondary antibody (Alexa Fluor 488 goat anti-rabbit IgG, Molecular Probes, Inc). Counterstaining was performed with DAPI (Sigma) for 5 min. UCP1 staining was analyzed on an Axio Imager 2 microscope (Zeiss). As a negative control, staining was performed on selected sections with isotype control.

### Measurement of oxygen consumption rate

Cellular respiration was measured using the Seahorse XF24 Analyzer (Agilent). On day 5 the instrument calibration was prepared following manufacturers protocol. After 23 hrs of stimulation with 1 μM retinol (Sigma), cells were washed with seahorse medium (Seahorse Biosience) and 630 μl assay medium (seahorse medium supplemented with 5 mM glucose (Sigma) and 1 mM sodium pyruvate (Gibco)) was added to all wells. Next, the plate was incubated for 1 hour in a CO_2_-free incubator at 37°C while the instrument was calibrated. Cellular oxygen consumption rates (OCR) was analyzed in an XF24 Flux Analyzer. Measurements were performed with repetitive cycles of 2 min mixture, 2 min wait and 4 min OCR measurement times. Injected compounds for the mitochondrial stress test were oligomycin (2□μM working concentration) to inhibit ATP synthase, followed by FCCP (1□μM working concentration) to induce mitochondrial uncoupling and rotenone (2□μM working concentration) to block the mitochondrial respiratory chain. Following the assay, medium was carefully removed and cells were harvested in RIPA buffer for the measurement of cellular protein content using the Pierce BCA Protein Assay Kit (Thermo Scientific) to adjust for potential differences in cell numbers.

### Statistics

The sample size is stated in the Figure and Table legends, respectively. Sample size calculation for mouse studies was performed by Java Applets for Power and Sample Size [Computer software]. Retrieved Mai 15, 2013, from http://www.stat.uiowa.edu/~rlenth/Power]. To evaluate differences of 50% with a standard deviation of 30%, a significance level of 5% and a power of 75%. The human data presented here were part of a cross sectional clinical study with the correlation between systemic retinol concentrations and cold-induced BAT activity as the primary outcome parameter. The approximate power to detect a given correlation at a level of 0.05 was calculated based on Fisher’s z transformation. In case of a true correlation of 0.4 a sample size of 30 was required for a power of 80%.

The statistical analysis was conducted using explorative data analysis and descriptive statistics. Results are given as mean ± standard error of the mean (SEM). Differences were analyzed by unpaired two-tailed Student’s t-test for two-group comparisons and with one-way ANOVA followed by Holm-Sidak’s *post-hoc* test for multiple group comparisons, respectively. Data were tested for Gaussian distribution using column statistics in GraphPad Prism 7. A p-value of ≤0.05 was considered statistically significant.

## Acknowledgements

This work was supported by the Vienna Science and Technology Fund, LS12-059 and Austrian Science Fund, P 27391 both to FWK. We thank Karin Strohmeier, Rita Lang, Sonja Seyfert and Helga Schachner for their excellent technical support as well as Audric Moses (Lipid Metabolite Analysis Core Facility, part of the Women and Children’s Health Research Institute and Faculty of Medicine and Dentistry at the University of Alberta, Edmonton, Canada) for performing FPLC analysis of plasma lipoproteins.

