## Supplemental Material for "Cold-mediated regulation of systemic retinol transport controls adipose tissue browning"

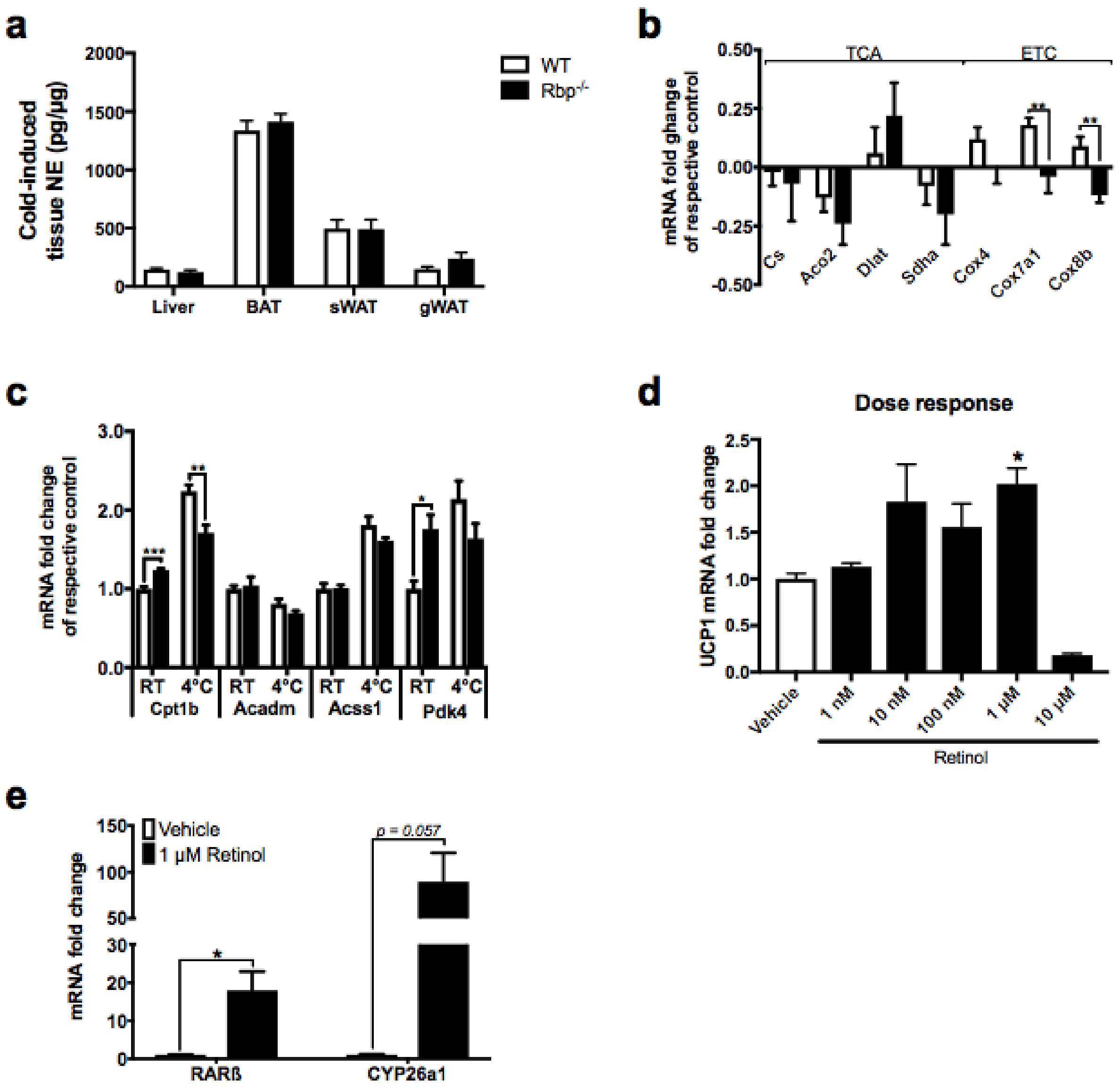


**Fig. S1**

Tissue NE content after cold exposure (**a**). Gene expression of TCA cycle genes and ETC genes in BAT (**b**). mRNA fold change of FA transport and FA oxidation markers in BAT (**c**). Retinol dose response (**d**). mRNA expression of RA target genes in hAPCs stimulated with 1 µM Retinol (**e**). * $p \leq 0.05$; ** $p \leq 0.01$; *** $p \leq 0.001$.

**Table S1**: Baseline characteristics. *** p≤0.001

| | | |
|---|---|---|
| **Age** | 28±0.7 | |
| **Sex (male:female)** | 12:18 | |
| **BMI (kg/m²)** | 22.2±0.3 | |
| **WHR** | 0.8±0.0 | |
| | **Before CE** | **After CE** |
| **Norepinephrine (ng/L)** | 188.7±15.2 | 651.7±43.2*** |

**Table S2**: Genes significantly differentially expressed between RT and 4°C in WT mice.

| gene | locus | raw p-value | Benjamini-Hochberg corrected p-value | mean fold change 4°C/RT (log2) | WT (RT) | WT (4°C) |
|---|---|---|---|---|---|---|
| Lrrc15 | chr16:30269301-30283254 | 5.00E-05 | 0.00283 | 4.049 | 0.040 | 0.661 |
| Slc25a34 | chr4:141618824-141623834 | 5.00E-05 | 0.00283 | 3.038 | 1.442 | 11.843 |
| Dio2 | chr12:90724551-90738438 | 5.00E-05 | 0.00283 | 2.916 | 2.003 | 15.121 |
| Perm1 | chr4:156215926-156221307 | 5.00E-05 | 0.00283 | 2.831 | 0.551 | 3.923 |
| Gyk | chrX:85701936-85776819 | 5.00E-05 | 0.00283 | 2.715 | 7.591 | 49.841 |
| Ucp1 | chr8:83290347-83298456 | 5.00E-05 | 0.00283 | 2.558 | 388.023 | 2284.810 |
| Cend1 | chr7:141426450-141429420 | 5.00E-05 | 0.00283 | 2.538 | 0.433 | 2.517 |
| Fabp3 | chr4:130308777-130315463 | 5.00E-05 | 0.00283 | 2.525 | 24.023 | 138.259 |
| Elovl3 | chr19:46131898-46148325 | 5.00E-05 | 0.00283 | 2.381 | 19.321 | 100.628 |
| Cyp2b10 | chr7:25897657-25926624 | 5.00E-05 | 0.00283 | 2.364 | 0.866 | 4.460 |
| Cthrc1 | chr15:39076931-39087119 | 5.00E-05 | 0.00283 | 2.315 | 0.957 | 4.763 |
| Slco4a1 | chr2:180460977-180474853 | 5.00E-05 | 0.00283 | 2.201 | 0.153 | 0.704 |
| Phospho1 | chr11:95824499-95845731 | 5.00E-05 | 0.00283 | 2.101 | 45.722 | 196.192 |
| Cdkn1a | chr17:29090985-29100722 | 5.00E-05 | 0.00283 | 1.968 | 11.321 | 44.277 |
| Mmp12 | chr9:7347373-7360461 | 5.00E-05 | 0.00283 | 1.906 | 0.480 | 1.799 |
| Gmpr | chr13:45507443-45546386 | 5.00E-05 | 0.00283 | 1.827 | 9.205 | 32.664 |
| Ankrd9 | chr12:110975352-110979021 | 5.00E-05 | 0.00283 | 1.678 | 5.139 | 16.449 |
| Pdk4 | chr6:5483350-5496278 | 5.00E-05 | 0.00283 | 1.562 | 75.162 | 221.926 |
| Adcy3 | chr12:4133396-4240123 | 5.00E-05 | 0.00283 | 1.544 | 13.107 | 38.224 |
| Nr4a1 | chr15:101266845-101274794 | 5.00E-05 | 0.00283 | 1.536 | 2.894 | 8.392 |
| Got1 | chr19:43499752-43524605 | 5.00E-05 | 0.00283 | 1.504 | 6.270 | 17.784 |
| Tuba8 | chr6:121210769-121226097 | 5.00E-05 | 0.00283 | 1.496 | 6.861 | 19.356 |
| Hpdl | chr4:116819906-116821508 | 5.00E-05 | 0.00283 | 1.463 | 2.380 | 6.562 |

| | | | | | | |
|---|---|---|---|---|---|---|
| Gadd45g | chr13:51846674-51848474 | 5.00E-05 | 0.00283 | 1.441 | 28.663 | 77.801 |
| Arntl | chr7:113207464-113314126 | 5.00E-05 | 0.00283 | 1.424 | 2.377 | 6.379 |
| Cpn2 | chr16:30256378-30267532 | 5.00E-05 | 0.00283 | 1.352 | 1.756 | 4.484 |
| Cyp26b1 | chr6:84571413-84593908 | 5.00E-05 | 0.00283 | 1.323 | 1.569 | 3.925 |
| Ccrn4l | chr3:51224446-51251654 | 5.00E-05 | 0.00283 | 1.297 | 11.056 | 27.172 |
| Slc27a2 | chr2:126553023-126588243 | 5.00E-05 | 0.00283 | 1.254 | 5.012 | 11.955 |
| Slc16a1 | chr3:104638663-104658462 | 5.00E-05 | 0.00283 | 1.248 | 15.684 | 37.263 |
| Pank1 | chr19:34806935-34879455 | 5.00E-05 | 0.00283 | 1.195 | 7.063 | 16.166 |
| Kcnk3 | chr5:30573986-30625270 | 5.00E-05 | 0.00283 | 1.151 | 29.863 | 66.300 |
| Ppif | chr14:25694169-25701282 | 5.00E-05 | 0.00283 | 1.150 | 21.727 | 48.202 |
| Pim1 | chr17:29491044-29495459 | 5.00E-05 | 0.00283 | 1.139 | 14.664 | 32.294 |
| Slc25a20 | chr9:108662097-108684641 | 5.00E-05 | 0.00283 | 1.124 | 95.643 | 208.479 |
| 6430571L13Rik | chr9:107340639-107349683 | 5.00E-05 | 0.00283 | 1.097 | 3.391 | 7.252 |
| Dtx4 | chr19:12466335-12501996 | 5.00E-05 | 0.00283 | 1.092 | 5.248 | 11.188 |
| Paqr9 | chr9:95559816-95562121 | 5.00E-05 | 0.00283 | 1.079 | 34.183 | 72.219 |
| Glrx | chr13:75839885-75850151 | 5.00E-05 | 0.00283 | 1.066 | 11.280 | 23.618 |
| Fam195a | chr17:25863697-25868738 | 5.00E-05 | 0.00283 | 1.062 | 80.428 | 167.916 |
| Traf4 | chr11:78158422-78165550 | 5.00E-05 | 0.00283 | 1.037 | 11.339 | 23.268 |
| Plin5 | chr17:56111600-56117548 | 5.00E-05 | 0.00283 | 1.033 | 32.612 | 66.731 |
| Cebpb | chr2:167688914-167690432 | 5.00E-05 | 0.00283 | 1.019 | 86.172 | 174.650 |
| Slc25a17 | chr15:81318920-81360765 | 5.00E-05 | 0.00283 | 1.009 | 53.266 | 107.175 |
| Igfbp3 | chr11:7206090-7213923 | 5.00E-05 | 0.00283 | 1.006 | 11.657 | 23.418 |
| Klb | chr5:65348410-65384003 | 5.00E-05 | 0.00283 | 0.991 | 16.343 | 32.477 |
| Impa2 | chr18:67289222-67318841 | 5.00E-05 | 0.00283 | 0.985 | 10.522 | 20.823 |
| Slmo2 | chr2:174465090-174472941 | 5.00E-05 | 0.00283 | 0.962 | 47.485 | 92.512 |
| Ucp3 | chr7:100472990-100486432 | 5.00E-05 | 0.00283 | 0.947 | 31.915 | 61.548 |
| Dusp10 | chr1:184034460-184075636 | 5.00E-05 | 0.00283 | 0.945 | 9.423 | 18.137 |
| Mtfp1 | chr11:4091480-4095431 | 5.00E-05 | 0.00283 | 0.944 | 14.051 | 27.034 |

| | | | | | | |
|---|---|---|---|---|---|---|
| Gys2 | chr6:142422612-142473109 | 5.00E-05 | 0.00283 | 0.938 | 32.135 | 61.575 |
| Ppargc1a | chr5:51454248-51553921 | 5.00E-05 | 0.00283 | 0.938 | 3.251 | 6.229 |
| Clstn3 | chr6:124430755-124464784 | 5.00E-05 | 0.00283 | 0.930 | 20.796 | 39.630 |
| Nfil3 | chr13:52967208-52981039 | 5.00E-05 | 0.00283 | 0.929 | 5.097 | 9.703 |
| Tmem37 | chr1:120067376-120073780 | 5.00E-05 | 0.00283 | 0.922 | 14.252 | 27.009 |
| Sik1 | chr17:31844249-31855792 | 5.00E-05 | 0.00283 | 0.919 | 4.856 | 9.179 |
| Ccl8 | chr11:82115184-82116799 | 5.00E-05 | 0.00283 | 0.915 | 87.231 | 164.471 |
| Ciapin1 | chr8:94819817-94838340 | 5.00E-05 | 0.00283 | 0.914 | 9.705 | 18.286 |
| Cox10 | chr11:63962626-64079472 | 5.00E-05 | 0.00283 | 0.909 | 8.632 | 16.208 |
| Adra1a | chr14:66635250-66771168 | 5.00E-05 | 0.00283 | 0.901 | 4.609 | 8.608 |
| 4931406C07Rik | chr9:15283336-15357788 | 5.00E-05 | 0.00283 | 0.896 | 127.805 | 237.810 |
| Klhl25 | chr7:75848337-75874130 | 5.00E-05 | 0.00283 | 0.893 | 4.283 | 7.953 |
| Serpina1b | chr12:103728155-103738189 | 5.00E-05 | 0.00283 | 0.886 | 28.648 | 52.950 |
| Cd1d2 | chr3:86986585-86989532 | 5.00E-05 | 0.00283 | 0.886 | 12.610 | 23.301 |
| Chchd10 | chr10:75935572-75940672 | 5.00E-05 | 0.00283 | 0.871 | 252.071 | 460.872 |
| Otop1 | chr5:38277403-38304217 | 5.00E-05 | 0.00283 | 0.864 | 26.233 | 47.760 |
| Ehhadh | chr16:21761284-21787834 | 5.00E-05 | 0.00283 | 0.863 | 37.107 | 67.500 |
| Gsta3 | chr1:21240584-21265575 | 5.00E-05 | 0.00283 | 0.849 | 83.137 | 149.791 |
| Cycs | chr6:50562562-50566474 | 5.00E-05 | 0.00283 | 0.847 | 22.512 | 40.496 |
| Hccs | chrX:169311530-169320343 | 5.00E-05 | 0.00283 | 0.847 | 12.401 | 22.300 |
| Ndufaf4 | chr4:24898082-24905001 | 5.00E-05 | 0.00283 | 0.836 | 9.038 | 16.136 |
| Dhrs11 | chr11:84820727-84829003 | 5.00E-05 | 0.00283 | 0.836 | 12.545 | 22.391 |
| Slc25a25 | chr2:32414486-32455476 | 5.00E-05 | 0.00283 | 0.822 | 13.556 | 23.965 |
| Mrpl47 | chr3:32727496-32736755 | 5.00E-05 | 0.00283 | 0.816 | 20.513 | 36.111 |
| Mid1ip1 | chrX:10717364-10719702 | 5.00E-05 | 0.00283 | 0.816 | 65.258 | 114.875 |
| Ppp1r3b | chr8:35375740-35388137 | 5.00E-05 | 0.00283 | 0.815 | 30.154 | 53.059 |
| Timm10 | chr2:84827020-84830213 | 5.00E-05 | 0.00283 | 0.805 | 41.204 | 71.989 |
| Apex2 | chrX:150571506-150588149 | 5.00E-05 | 0.00283 | 0.804 | 6.148 | 10.734 |

| | | | | | | |
|---|---|---|---|---|---|---|
| Tbrg4 | chr11:6615597-6626067 | 5.00E-05 | 0.00283 | 0.803 | 19.190 | 33.489 |
| Itpk1 | chr12:102568582-102704869 | 5.00E-05 | 0.00283 | 0.793 | 21.573 | 37.376 |
| Hacd2 | chr16:35022420-35109175 | 5.00E-05 | 0.00283 | 0.792 | 49.380 | 85.478 |
| Ldhb | chr6:142490248-142507957 | 5.00E-05 | 0.00283 | 0.785 | 102.223 | 176.162 |
| Mrps2 | chr2:28468065-28471177 | 5.00E-05 | 0.00283 | 0.782 | 20.802 | 35.778 |
| Timm23 | chr14:32180165-32201891 | 5.00E-05 | 0.00283 | 0.781 | 18.299 | 31.450 |
| Adrbk2 | chr5:112910477-113015538 | 5.00E-05 | 0.00283 | 0.777 | 6.384 | 10.937 |
| Pla2g12a | chr3:129878605-129895825 | 5.00E-05 | 0.00283 | 0.776 | 43.351 | 74.225 |
| Adam12 | chr7:133883198-134225097 | 5.00E-05 | 0.00283 | 0.775 | 5.561 | 9.514 |
| Asns | chr6:7675170-7693182 | 5.00E-05 | 0.00283 | 0.761 | 44.335 | 75.144 |
| Mlycd | chr8:119394891-119411088 | 5.00E-05 | 0.00283 | 0.761 | 35.637 | 60.396 |
| Hdhd3 | chr4:62499053-62502200 | 5.00E-05 | 0.00283 | 0.748 | 26.678 | 44.794 |
| Letm1 | chr5:33741351-33782704 | 5.00E-05 | 0.00283 | 0.747 | 45.947 | 77.117 |
| Mrs2 | chr13:24992294-25020317 | 5.00E-05 | 0.00283 | 0.741 | 12.097 | 20.221 |
| Tmed5 | chr5:108121646-108132591 | 5.00E-05 | 0.00283 | 0.739 | 36.340 | 60.664 |
| Khdrbs3 | chr15:68928419-69093518 | 5.00E-05 | 0.00283 | 0.738 | 10.326 | 17.221 |
| Syt12 | chr19:4425456-4477143 | 5.00E-05 | 0.00283 | 0.737 | 20.381 | 33.978 |
| Letmd1 | chr15:100469033-100479252 | 5.00E-05 | 0.00283 | 0.735 | 54.995 | 91.545 |
| Mrps18b | chr17:35910384-35916369 | 5.00E-05 | 0.00283 | 0.726 | 23.433 | 38.768 |
| Mrpl12 | chr11:120484668-120488754 | 5.00E-05 | 0.00283 | 0.724 | 139.112 | 229.847 |
| Etfdh | chr3:79603787-79628767 | 5.00E-05 | 0.00283 | 0.724 | 114.364 | 188.906 |
| 2010003K11Rik | chr19:4496787-4498583 | 5.00E-05 | 0.00283 | 0.722 | 19.985 | 32.955 |
| Pgam1 | chr19:41911870-41918665 | 5.00E-05 | 0.00283 | 0.717 | 23.380 | 38.423 |
| Gmps | chr3:63976142-64019078 | 5.00E-05 | 0.00283 | 0.716 | 8.837 | 14.513 |
| Tomm5 | chr4:45105209-45108113 | 5.00E-05 | 0.00283 | 0.715 | 39.754 | 65.234 |
| Fuom | chr7:140097814-140102441 | 5.00E-05 | 0.00283 | 0.714 | 8.514 | 13.969 |
| Tpcn2 | chr7:145253922-145283927 | 5.00E-05 | 0.00283 | 0.713 | 4.965 | 8.139 |
| Tmem147 | chr7:30727700-30729534 | 5.00E-05 | 0.00283 | 0.709 | 99.624 | 162.833 |

| | | | | | | |
|---|---|---|---|---|---|---|
| Slc25a51 | chr4:45395923-45408766 | 5.00E-05 | 0.00283 | 0.708 | 44.652 | 72.939 |
| Rxrg | chr1:167598361-167639623 | 5.00E-05 | 0.00283 | 0.707 | 17.030 | 27.793 |
| Trim67 | chr8:124793018-124834704 | 5.00E-05 | 0.00283 | 0.706 | 2.915 | 4.756 |
| Apmap | chr2:150583080-150608523 | 5.00E-05 | 0.00283 | 0.706 | 161.231 | 263.052 |
| Car13 | chr3:14641726-14663002 | 5.00E-05 | 0.00283 | 0.703 | 13.944 | 22.698 |
| Tmem79 | chr3:88328652-88334433 | 5.00E-05 | 0.00283 | 0.702 | 22.268 | 36.228 |
| Gtf2ird1 | chr5:134357660-134456716 | 5.00E-05 | 0.00283 | 0.701 | 7.689 | 12.502 |
| Dock8 | chr19:24999528-25202432 | 5.00E-05 | 0.00283 | 0.701 | 4.430 | 7.201 |
| Pdp2 | chr8:104591467-104596849 | 5.00E-05 | 0.00283 | 0.698 | 8.386 | 13.608 |
| 2300009A05Rik | chr9:63394446-63399244 | 5.00E-05 | 0.00283 | 0.693 | 35.391 | 57.213 |
| Esrra | chr19:6909697-6921808 | 5.00E-05 | 0.00283 | 0.692 | 44.365 | 71.677 |
| Hspa8 | chr9:40801272-40805199 | 5.00E-05 | 0.00283 | 0.691 | 135.586 | 218.832 |
| Pgk1 | chrX:106187099-106203699 | 5.00E-05 | 0.00283 | 0.690 | 51.261 | 82.703 |
| Dnajc11 | chr4:151933719-151981959 | 5.00E-05 | 0.00283 | 0.688 | 21.646 | 34.879 |
| Tomm40 | chr7:19701312-19715429 | 5.00E-05 | 0.00283 | 0.687 | 49.320 | 79.388 |
| Aspa | chr11:73304987-73324637 | 5.00E-05 | 0.00283 | 0.679 | 52.020 | 83.258 |
| Slc4a4 | chr5:88887259-89239656 | 5.00E-05 | 0.00283 | 0.675 | 6.699 | 10.694 |
| Mocs2 | chr13:114818236-114829420 | 5.00E-05 | 0.00283 | 0.675 | 73.861 | 117.887 |
| Atad3a | chr4:155740639-155761098 | 5.00E-05 | 0.00283 | 0.671 | 33.903 | 53.989 |
| Ubiad1 | chr4:148434496-148444751 | 5.00E-05 | 0.00283 | 0.671 | 7.103 | 11.310 |
| Ptges | chr2:30889470-30903297 | 5.00E-05 | 0.00283 | 0.669 | 29.413 | 46.765 |
| Slc25a42 | chr8:70184339-70212281 | 5.00E-05 | 0.00283 | 0.660 | 22.664 | 35.811 |
| Ndufa12 | chr10:94199008-94220948 | 5.00E-05 | 0.00283 | 0.658 | 117.947 | 186.152 |
| Mrpl20 | chr4:155803617-155808829 | 5.00E-05 | 0.00283 | 0.657 | 154.012 | 242.771 |
| Rgs2 | chr1:143999337-144004149 | 5.00E-05 | 0.00283 | 0.652 | 9.347 | 14.683 |
| Prodh | chr16:18071725-18089190 | 5.00E-05 | 0.00283 | 0.648 | 21.170 | 33.183 |
| Ifrd1 | chr12:40203128-40223189 | 5.00E-05 | 0.00283 | 0.647 | 62.703 | 98.214 |
| Sdhd | chr9:50596339-50603849 | 5.00E-05 | 0.00283 | 0.645 | 248.257 | 388.217 |

| | | | | | | |
|---|---|---|---|---|---|---|
| Tatdn2 | chr6:113697498-113711068 | 5.00E-05 | 0.00283 | 0.641 | 18.184 | 28.361 |
| Ppara | chr15:85735563-85806851 | 5.00E-05 | 0.00283 | 0.639 | 9.286 | 14.465 |
| Adig | chr2:158502611-158508198 | 5.00E-05 | 0.00283 | 0.632 | 339.815 | 526.623 |
| Spsb3 | chr17:24886673-24892147 | 5.00E-05 | 0.00283 | 0.626 | 17.057 | 26.321 |
| Pcyt2 | chr11:120610086-120617890 | 5.00E-05 | 0.00283 | 0.624 | 44.759 | 68.987 |
| Slc7a6 | chr8:106168874-106198704 | 5.00E-05 | 0.00283 | 0.622 | 10.100 | 15.538 |
| Slc25a39 | chr11:102402975-102407517 | 5.00E-05 | 0.00283 | 0.619 | 141.869 | 217.950 |
| Cars2 | chr8:11514016-11550771 | 5.00E-05 | 0.00283 | 0.619 | 11.626 | 17.853 |
| Timm17a | chr1:135301534-135313737 | 5.00E-05 | 0.00283 | 0.613 | 90.884 | 138.971 |
| Emc6 | chr11:73175502-73177042 | 5.00E-05 | 0.00283 | 0.611 | 73.523 | 112.265 |
| Dnaja3 | chr16:4684069-4707693 | 5.00E-05 | 0.00283 | 0.610 | 47.777 | 72.943 |
| Plbd1 | chr6:136612070-136661893 | 5.00E-05 | 0.00283 | 0.610 | 57.511 | 87.796 |
| Nabp1 | chr1:51469487-51478399 | 5.00E-05 | 0.00283 | 0.609 | 38.763 | 59.140 |
| 1700021F05Rik | chr10:43525120-43540994 | 5.00E-05 | 0.00283 | 0.606 | 46.538 | 70.843 |
| Mapkapk2 | chr1:131053703-131097543 | 5.00E-05 | 0.00283 | 0.605 | 52.681 | 80.128 |
| Gars | chr6:55038000-55079504 | 5.00E-05 | 0.00283 | 0.604 | 59.601 | 90.599 |
| Mrap | chr16:90738323-90749776 | 5.00E-05 | 0.00283 | 0.596 | 232.267 | 350.960 |
| Nudt19 | chr7:35547184-35555928 | 5.00E-05 | 0.00283 | 0.591 | 29.469 | 44.388 |
| Gja1 | chr10:56377299-56390419 | 5.00E-05 | 0.00283 | 0.585 | 31.392 | 47.082 |
| Pdhx | chr2:103021056-103073513 | 5.00E-05 | 0.00283 | 0.585 | 45.648 | 68.454 |
| Dcaf6 | chr1:165329500-165460463 | 5.00E-05 | 0.00283 | 0.584 | 21.054 | 31.569 |
| Ndufa11 | chr17:56717761-56724248 | 5.00E-05 | 0.00283 | 0.583 | 29.139 | 43.638 |
| Rbpms2 | chr9:65630581-65660518 | 5.00E-05 | 0.00283 | 0.582 | 24.215 | 36.260 |
| Phb2 | chr6:124712288-124716945 | 5.00E-05 | 0.00283 | 0.582 | 131.539 | 196.948 |
| Rhot2 | chr17:25838837-25861515 | 5.00E-05 | 0.00283 | 0.582 | 19.377 | 29.009 |
| Scarb2 | chr5:92443872-92505608 | 5.00E-05 | 0.00283 | 0.581 | 43.752 | 65.466 |
| Fam96a | chr9:66126610-66138968 | 5.00E-05 | 0.00283 | 0.581 | 37.433 | 55.989 |
| Pkn1 | chr8:83666639-83699179 | 5.00E-05 | 0.00283 | 0.577 | 31.050 | 46.323 |

| | | | | | | |
|---|---|---|---|---|---|---|
| Afg3l2 | chr18:67404763-67449136 | 5.00E-05 | 0.00283 | 0.576 | 41.500 | 61.869 |
| Abhd5 | chr9:122351615-122381523 | 5.00E-05 | 0.00283 | 0.576 | 59.464 | 88.634 |
| Dapk1 | chr13:60601946-60763191 | 5.00E-05 | 0.00283 | 0.574 | 14.888 | 22.161 |
| Slc3a2 | chr19:8706881-8723369 | 5.00E-05 | 0.00283 | 0.572 | 46.093 | 68.539 |
| Isca1 | chr13:59755414-59769789 | 5.00E-05 | 0.00283 | 0.566 | 44.168 | 65.402 |
| Oxa1l | chr14:54360840-54417702 | 5.00E-05 | 0.00283 | 0.565 | 29.396 | 43.492 |
| Tubb4b | chr2:25218744-25224702 | 5.00E-05 | 0.00283 | 0.563 | 85.028 | 125.575 |
| Ube2m | chr7:13035119-13038275 | 5.00E-05 | 0.00283 | 0.562 | 114.863 | 169.532 |
| Ndufa8 | chr2:36036333-36049292 | 5.00E-05 | 0.00283 | 0.558 | 221.640 | 326.267 |
| Lrrc8d | chr5:105699968-105815215 | 5.00E-05 | 0.00283 | 0.548 | 18.643 | 27.265 |
| Ddt | chr10:75771232-75773374 | 5.00E-05 | 0.00283 | 0.542 | 197.443 | 287.572 |
| BC004004 | chr17:29268787-29302887 | 5.00E-05 | 0.00283 | 0.537 | 35.686 | 51.789 |
| Ptprs | chr17:56412425-56476480 | 5.00E-05 | 0.00283 | -0.544 | 9.978 | 6.843 |
| Cd200 | chr16:45382134-45409053 | 5.00E-05 | 0.00283 | -0.598 | 22.870 | 15.105 |
| Aplnr | chr2:85136359-85139923 | 5.00E-05 | 0.00283 | -0.599 | 13.522 | 8.928 |
| Podn | chr4:108014792-108032090 | 5.00E-05 | 0.00283 | -0.617 | 67.036 | 43.718 |
| Eepd1 | chr9:25481596-25604110 | 5.00E-05 | 0.00283 | -0.625 | 75.350 | 48.852 |
| Col6a3 | chr1:90766859-90843971 | 5.00E-05 | 0.00283 | -0.631 | 50.747 | 32.760 |
| Rgs5 | chr1:169655500-169693526 | 5.00E-05 | 0.00283 | -0.637 | 57.146 | 36.756 |
| Zbtb16 | chr9:48654296-48835945 | 5.00E-05 | 0.00283 | -0.678 | 13.134 | 8.210 |
| Trp53i11 | chr2:93187583-93201757 | 5.00E-05 | 0.00283 | -0.708 | 13.301 | 8.144 |
| Bhlhe41 | chr6:145858242-145865420 | 5.00E-05 | 0.00283 | -0.726 | 13.708 | 8.285 |
| Nr1d2 | chr14:18204055-18239106 | 5.00E-05 | 0.00283 | -0.757 | 25.861 | 15.301 |
| Lctl | chr9:64117146-64172931 | 5.00E-05 | 0.00283 | -0.781 | 18.730 | 10.899 |
| Syne3 | chr12:104929932-105009809 | 5.00E-05 | 0.00283 | -0.783 | 5.512 | 3.203 |
| Hmgcr | chr13:96648961-96670936 | 5.00E-05 | 0.00283 | -0.787 | 7.987 | 4.629 |
| Adrb3 | chr8:27225775-27229588 | 5.00E-05 | 0.00283 | -0.790 | 235.673 | 136.303 |
| Fam20c | chr5:138755080-138810063 | 5.00E-05 | 0.00283 | -0.797 | 22.778 | 13.113 |

| | | | | | | |
|---|---|---|---|---|---|---|
| Per2 | chr1:91415981-91459328 | 5.00E-05 | 0.00283 | -0.846 | 6.654 | 3.703 |
| Ntrk3 | chr7:78192113-78577838 | 5.00E-05 | 0.00283 | -0.856 | 15.881 | 8.775 |
| Per3 | chr4:151003654-151044665 | 5.00E-05 | 0.00283 | -0.859 | 21.538 | 11.873 |
| Far1 | chr7:113513833-113570888 | 5.00E-05 | 0.00283 | -0.879 | 15.142 | 8.232 |
| Snhg11 | chr2:158375637-158386145 | 5.00E-05 | 0.00283 | -0.892 | 20.632 | 11.116 |
| Cd209a | chr8:3743394-3748984 | 5.00E-05 | 0.00283 | -0.908 | 13.683 | 7.293 |
| Kcnh2 | chr5:24319588-24351604 | 5.00E-05 | 0.00283 | -0.936 | 4.973 | 2.599 |
| Idi1 | chr13:8885605-8892396 | 5.00E-05 | 0.00283 | -0.943 | 5.108 | 2.657 |
| Rprml | chr11:103649508-103650580 | 5.00E-05 | 0.00283 | -1.063 | 13.081 | 6.260 |
| Anxa8 | chr14:34051129-34102754 | 5.00E-05 | 0.00283 | -1.089 | 15.142 | 7.119 |
| Celsr2 | chr3:108390847-108415494 | 5.00E-05 | 0.00283 | -1.151 | 4.603 | 2.073 |
| Rps3a1 | chr3:86137939-86142668 | 5.00E-05 | 0.00283 | -1.238 | 152.827 | 64.777 |
| Rab6b | chr9:103112073-103185270 | 5.00E-05 | 0.00283 | -1.249 | 4.909 | 2.066 |
| Crtac1 | chr19:42283036-42431783 | 5.00E-05 | 0.00283 | -1.480 | 11.007 | 3.947 |
| Dbp | chr7:45705246-45718002 | 5.00E-05 | 0.00283 | -1.489 | 143.170 | 51.007 |
| Pnpla5 | chr15:84112620-84123175 | 5.00E-05 | 0.00283 | -1.773 | 2.253 | 0.659 |
| Erdr1 | chrY:90785441-90816465 | 5.00E-05 | 0.00283 | -1.934 | 11.525 | 3.016 |
| Fabp1 | chr6:71199887-71205023 | 5.00E-05 | 0.00283 | -1.999 | 40.635 | 10.166 |
| Gbp11 | chr5:105323025-105346476 | 5.00E-05 | 0.00283 | -2.003 | 8.287 | 2.067 |
| Gdpd3 | chr7:126766413-126775645 | 5.00E-05 | 0.00283 | -2.177 | 35.579 | 7.868 |
| Otof | chr5:30367065-30461932 | 5.00E-05 | 0.00283 | -2.226 | 0.598 | 0.128 |
| Fam84a | chr12:14147597-14152038 | 5.00E-05 | 0.00283 | -2.312 | 1.485 | 0.299 |
| Calml3 | chr13:3802892-3804318 | 5.00E-05 | 0.00283 | -2.733 | 2.994 | 0.450 |
| Krt5 | chr15:101707069-101712891 | 5.00E-05 | 0.00283 | -2.945 | 1.740 | 0.226 |
| Trim29 | chr9:43310762-43336125 | 5.00E-05 | 0.00283 | -3.050 | 1.689 | 0.204 |
| Perp | chr10:18845070-18857072 | 5.00E-05 | 0.00283 | -3.135 | 5.147 | 0.586 |
| S100a14 | chr3:90526848-90528835 | 5.00E-05 | 0.00283 | -3.503 | 2.481 | 0.219 |
| Pkp1 | chr1:135871394-135919024 | 5.00E-05 | 0.00283 | -3.911 | 0.851 | 0.057 |

| | | | | | | |
|---|---|---|---|---|---|---|
| Asprv1 | chr6:86628173-86629704 | 5.00E-05 | 0.00283 | -4.420 | 4.187 | 0.196 |
| Klk10 | chr7:43781053-43785410 | 5.00E-05 | 0.00283 | -4.458 | 6.878 | 0.313 |
| Dsp | chr13:38151293-38198577 | 5.00E-05 | 0.00283 | -4.794 | 1.392 | 0.050 |
| Krt14 | chr11:100203161-100207510 | 5.00E-05 | 0.00283 | -5.843 | 7.741 | 0.135 |
| Awat2 | chrX:100402221-100442717 | 5.00E-05 | 0.00283 | -6.197 | 15.965 | 0.218 |
| Cst6 | chr19:5344704-5349574 | 5.00E-05 | 0.00283 | -7.845 | 15.023 | 0.065 |
| Idh3a | chr9:54586510-54604662 | 1.00E-04 | 0.00503 | 0.878 | 252.354 | 463.852 |
| Cidea | chr18:67321208-67367794 | 1.00E-04 | 0.00503 | 0.806 | 315.951 | 552.463 |
| Rnmtl1 | chr11:76243735-76250622 | 1.00E-04 | 0.00503 | 0.745 | 9.053 | 15.172 |
| Mpc2 | chr1:165461207-165481214 | 1.00E-04 | 0.00503 | 0.703 | 485.695 | 790.553 |
| Pmf1 | chr3:88394142-88410316 | 1.00E-04 | 0.00503 | 0.685 | 19.847 | 31.919 |
| Hsd17b7 | chr1:169949536-169969205 | 1.00E-04 | 0.00503 | 0.671 | 3.734 | 5.945 |
| Mrpl15 | chr1:4773199-4785726 | 1.00E-04 | 0.00503 | 0.623 | 11.299 | 17.401 |
| Gpam | chr19:55069733-55099447 | 1.00E-04 | 0.00503 | 0.607 | 54.733 | 83.339 |
| Ppp1r1a | chr15:103530278-103537992 | 1.00E-04 | 0.00503 | 0.604 | 73.646 | 111.918 |
| Alas1 | chr9:106233454-106247954 | 1.00E-04 | 0.00503 | 0.603 | 97.653 | 148.303 |
| Akirin1 | chr4:123735194-123750299 | 1.00E-04 | 0.00503 | 0.586 | 20.622 | 30.954 |
| Xrcc6 | chr15:82016368-82040084 | 1.00E-04 | 0.00503 | 0.573 | 21.764 | 32.378 |
| Txn2 | chr15:77915050-77928994 | 1.00E-04 | 0.00503 | 0.573 | 126.314 | 187.882 |
| Cd59a | chr2:104095800-104115410 | 1.00E-04 | 0.00503 | 0.567 | 29.918 | 44.329 |
| Ptch2 | chr4:117096355-117114831 | 1.00E-04 | 0.00503 | 0.562 | 10.463 | 15.443 |
| Lamtor1 | chr7:101899807-101911903 | 1.00E-04 | 0.00503 | 0.561 | 108.961 | 160.756 |
| Rbm3 | chrX:8138974-8147963 | 1.00E-04 | 0.00503 | 0.558 | 65.451 | 96.344 |
| Naa20 | chr2:145903240-145916425 | 1.00E-04 | 0.00503 | 0.551 | 34.663 | 50.788 |
| Snx10 | chr6:51523902-51590670 | 1.00E-04 | 0.00503 | 0.551 | 45.821 | 67.123 |
| 1110008F13Rik | chr2:156863121-156887078 | 1.00E-04 | 0.00503 | 0.550 | 144.486 | 211.526 |
| Pmm1 | chr15:81951105-81960930 | 1.00E-04 | 0.00503 | 0.547 | 76.658 | 112.038 |
| Inmt | chr6:55170626-55174990 | 1.00E-04 | 0.00503 | 0.543 | 128.417 | 187.090 |

| | | | | | | |
|---|---|---|---|---|---|---|
| Dusp1 | chr17:26505590-26508472 | 1.00E-04 | 0.00503 | 0.540 | 22.672 | 32.957 |
| Inhbb | chr1:119415464-119422248 | 1.00E-04 | 0.00503 | -0.536 | 20.251 | 13.965 |
| Prps1 | chrX:140456602-140476140 | 1.00E-04 | 0.00503 | -0.556 | 58.541 | 39.811 |
| Mgl2 | chr11:70130356-70137542 | 1.00E-04 | 0.00503 | -0.579 | 48.804 | 32.678 |
| Fam102b | chr3:108970996-109027607 | 1.00E-04 | 0.00503 | -0.583 | 19.952 | 13.319 |
| Ankrd12 | chr17:65967500-66077046 | 1.00E-04 | 0.00503 | -0.637 | 3.930 | 2.527 |
| Dock10 | chr1:80501067-80758553 | 1.00E-04 | 0.00503 | -0.678 | 4.910 | 3.069 |
| Krt17 | chr11:100256216-100260989 | 1.00E-04 | 0.00503 | -5.667 | 7.179 | 0.141 |
| Arhgef37 | chr18:61493793-61536536 | 1.50E-04 | 0.00683 | 1.250 | 1.290 | 3.068 |
| Cxcl13 | chr5:95956938-95961068 | 1.50E-04 | 0.00683 | 0.814 | 8.943 | 15.724 |
| Mir6236 | chr9:110281286-110281409 | 1.50E-04 | 0.00683 | 0.731 | 4041.790 | 6710.550 |
| Mgst2 | chr3:51661192-51682675 | 1.50E-04 | 0.00683 | 0.704 | 51.791 | 84.391 |
| Cluh | chr11:74649494-74670847 | 1.50E-04 | 0.00683 | 0.695 | 89.937 | 145.549 |
| Slc35g1 | chr19:38395979-38405607 | 1.50E-04 | 0.00683 | 0.656 | 5.546 | 8.739 |
| Tigar | chr6:127085115-127109552 | 1.50E-04 | 0.00683 | 0.636 | 5.920 | 9.202 |
| Leo1 | chr9:75441523-75466432 | 1.50E-04 | 0.00683 | 0.623 | 10.211 | 15.728 |
| Acsl5 | chr19:55253368-55296628 | 1.50E-04 | 0.00683 | 0.621 | 38.970 | 59.955 |
| Kcnb1 | chr2:167095968-167188818 | 1.50E-04 | 0.00683 | 0.588 | 6.783 | 10.199 |
| Timm22 | chr11:76406924-76416313 | 1.50E-04 | 0.00683 | 0.588 | 11.702 | 17.591 |
| Dusp6 | chr10:99263230-99267489 | 1.50E-04 | 0.00683 | 0.578 | 12.862 | 19.205 |
| Mrpl34 | chr8:71464925-71465753 | 1.50E-04 | 0.00683 | 0.576 | 72.196 | 107.591 |
| Lrrc59 | chr11:94629823-94653754 | 1.50E-04 | 0.00683 | 0.570 | 66.964 | 99.395 |
| Phb | chr11:95666956-95680773 | 1.50E-04 | 0.00683 | 0.561 | 33.981 | 50.131 |
| Akap1 | chr11:88830791-88881395 | 1.50E-04 | 0.00683 | 0.546 | 36.238 | 52.926 |
| Rn45s | chr17:39842996-39848829 | 1.50E-04 | 0.00683 | 0.530 | 16.329 | 23.576 |
| Ptpmt1 | chr2:90910712-90918050 | 1.50E-04 | 0.00683 | 0.512 | 46.012 | 65.630 |
| Pyurf | chr6:57684738-57692078 | 1.50E-04 | 0.00683 | 0.498 | 10.383 | 14.668 |
| Rcn3 | chr7:45082913-45092213 | 1.50E-04 | 0.00683 | -0.554 | 48.773 | 33.216 |

| | | | | | | |
|---|---|---|---|---|---|---|
| Rcn1 | chr2:105385947-105399319 | 1.50E-04 | 0.00683 | -0.557 | 18.305 | 12.441 |
| Fry | chr5:150259929-150497753 | 1.50E-04 | 0.00683 | -0.595 | 22.445 | 14.855 |
| S100b | chr10:76253835-76261319 | 1.50E-04 | 0.00683 | -0.668 | 16.147 | 10.164 |
| Dhcr24 | chr4:106561037-106589113 | 1.50E-04 | 0.00683 | -0.675 | 13.496 | 8.455 |
| Fads2 | chr19:10064163-10101503 | 1.50E-04 | 0.00683 | -0.802 | 12.585 | 7.218 |
| Ampd1 | chr3:103074013-103099720 | 1.50E-04 | 0.00683 | -1.228 | 4.440 | 1.896 |
| Ehf | chr2:103263432-103303196 | 1.50E-04 | 0.00683 | -2.565 | 0.478 | 0.081 |
| Dsc2 | chr18:20030797-20059505 | 1.50E-04 | 0.00683 | -4.174 | 0.676 | 0.037 |
| Dhrs9 | chr2:69380461-69403086 | 2.00E-04 | 0.00889 | 1.543 | 0.707 | 2.060 |
| Lncbate1 | chr8:108553251-108584677 | 2.00E-04 | 0.00889 | 0.803 | 14.630 | 25.519 |
| Ptcd2 | chr13:99319648-99344678 | 2.00E-04 | 0.00889 | 0.586 | 20.732 | 31.118 |
| Manf | chr9:106887414-106891938 | 2.00E-04 | 0.00889 | 0.535 | 22.879 | 33.156 |
| Per1 | chr11:69098955-69109957 | 2.00E-04 | 0.00889 | -0.582 | 13.158 | 8.792 |
| Aldh1a3 | chr7:66390892-66427477 | 2.00E-04 | 0.00889 | -0.589 | 9.312 | 6.190 |
| Apoa1 | chr9:46228629-46230469 | 2.00E-04 | 0.00889 | -0.854 | 29.569 | 16.358 |
| Cpt1b | chr15:89416404-89429927 | 2.50E-04 | 0.01055 | 1.276 | 45.667 | 110.559 |
| Osgin1 | chr8:119437161-119446256 | 2.50E-04 | 0.01055 | 1.033 | 2.775 | 5.678 |
| Smyd4 | chr11:75348432-75405705 | 2.50E-04 | 0.01055 | 0.748 | 3.441 | 5.777 |
| Cd44 | chr2:102811141-102901665 | 2.50E-04 | 0.01055 | 0.742 | 3.136 | 5.244 |
| Timm9 | chr12:71111427-71136675 | 2.50E-04 | 0.01055 | 0.670 | 26.019 | 41.396 |
| Acot11 | chr4:106733914-106799831 | 2.50E-04 | 0.01055 | 0.644 | 20.104 | 31.420 |
| Spry4 | chr18:38586264-38601268 | 2.50E-04 | 0.01055 | 0.609 | 5.619 | 8.569 |
| Sod2 | chr17:13007838-13018119 | 2.50E-04 | 0.01055 | 0.601 | 69.417 | 105.286 |
| Pde4a | chr9:21165713-21226281 | 2.50E-04 | 0.01055 | 0.594 | 5.119 | 7.728 |
| Ndufb10 | chr17:24722066-24724388 | 2.50E-04 | 0.01055 | 0.553 | 351.421 | 515.532 |
| Sbk1 | chr7:126272618-126294999 | 2.50E-04 | 0.01055 | 0.544 | 27.537 | 40.144 |
| Memo1 | chr17:74200699-74294863 | 2.50E-04 | 0.01055 | 0.542 | 31.898 | 46.457 |
| 2010107E04Rik | chr12:111961375-111966977 | 2.50E-04 | 0.01055 | 0.528 | 526.784 | 759.530 |

| | | | | | | |
|---|---|---|---|---|---|---|
| Atxn2 | chr5:121711608-121814950 | 2.50E-04 | 0.01055 | 0.519 | 13.710 | 19.640 |
| Cadm3 | chr1:173334253-173367695 | 2.50E-04 | 0.01055 | -0.545 | 14.366 | 9.849 |
| Ptprf | chr4:118208212-118291397 | 2.50E-04 | 0.01055 | -0.616 | 4.215 | 2.751 |
| Timm8b | chr9:50603900-50625000 | 3.00E-04 | 0.01232 | 0.596 | 142.707 | 215.659 |
| Slc19a1 | chr10:77032738-77050432 | 3.00E-04 | 0.01232 | 0.531 | 16.154 | 23.346 |
| Slc43a2 | chr11:75531693-75577572 | 3.00E-04 | 0.01232 | 0.519 | 5.785 | 8.290 |
| Dusp4 | chr8:34807609-34819894 | 3.00E-04 | 0.01232 | 0.517 | 27.235 | 38.982 |
| Mtx2 | chr2:74825811-74876748 | 3.00E-04 | 0.01232 | 0.513 | 64.103 | 91.485 |
| Armc1 | chr3:19132143-19163065 | 3.00E-04 | 0.01232 | 0.500 | 19.204 | 27.165 |
| Fbxo45 | chr16:32230111-32247025 | 3.00E-04 | 0.01232 | -0.489 | 18.969 | 13.515 |
| Krt79 | chr15:101929331-101940324 | 3.00E-04 | 0.01232 | -0.797 | 12.896 | 7.423 |
| Mycl | chr4:122995930-123002480 | 3.00E-04 | 0.01232 | -1.048 | 3.549 | 1.716 |
| Mtch2 | chr2:90847154-90866634 | 3.50E-04 | 0.01398 | 0.608 | 134.774 | 205.372 |
| Cd320 | chr17:33843090-33849774 | 3.50E-04 | 0.01398 | 0.591 | 12.006 | 18.090 |
| Sqle | chr15:59315091-59331193 | 3.50E-04 | 0.01398 | 0.587 | 10.769 | 16.176 |
| Abhd6 | chr14:8002901-8056555 | 3.50E-04 | 0.01398 | 0.577 | 13.493 | 20.129 |
| G0s2 | chr1:193272159-193273188 | 3.50E-04 | 0.01398 | 0.550 | 381.002 | 557.748 |
| Mpc1 | chr17:8283812-8327442 | 3.50E-04 | 0.01398 | 0.543 | 157.004 | 228.773 |
| Mtpap | chr18:4375591-4397330 | 3.50E-04 | 0.01398 | 0.533 | 16.640 | 24.070 |
| Gde1 | chr7:118688557-118705738 | 3.50E-04 | 0.01398 | 0.521 | 104.892 | 150.462 |
| Gpr153 | chr4:152274361-152285337 | 3.50E-04 | 0.01398 | -0.659 | 7.054 | 4.469 |
| Apln | chrX:48025145-48034852 | 4.00E-04 | 0.01548 | 0.874 | 2.226 | 4.079 |
| Egln3 | chr12:54178980-54203874 | 4.00E-04 | 0.01548 | 0.829 | 3.129 | 5.557 |
| Rdh13 | chr7:4425664-4445657 | 4.00E-04 | 0.01548 | 0.567 | 9.215 | 13.648 |
| Jagn1 | chr6:113442516-113448229 | 4.00E-04 | 0.01548 | 0.540 | 30.354 | 44.149 |
| Arfgef2 | chr2:166805580-166898051 | 4.00E-04 | 0.01548 | 0.520 | 9.986 | 14.321 |
| Atpaf2 | chr11:60400623-60417099 | 4.00E-04 | 0.01548 | 0.512 | 56.691 | 80.869 |
| Atp6v0a2 | chr5:124629051-124724455 | 4.00E-04 | 0.01548 | 0.496 | 16.885 | 23.819 |

| | | | | | | |
|---|---|---|---|---|---|---|
| Efr3a | chr15:65787040-65873812 | 4.00E-04 | 0.01548 | 0.484 | 18.722 | 26.186 |
| Fam114a1 | chr5:64970074-65041901 | 4.00E-04 | 0.01548 | -0.542 | 16.315 | 11.205 |
| Sult1e1 | chr5:87575967-87591611 | 4.00E-04 | 0.01548 | -0.760 | 16.867 | 9.958 |
| Btla | chr16:45224336-45252895 | 4.00E-04 | 0.01548 | -1.469 | 1.064 | 0.384 |
| A530050N04Rik | chr18:61470224-61484607 | 4.50E-04 | 0.01664 | 4.225 | 0.144 | 2.702 |
| Trp53inp2 | chr2:155381855-155389847 | 4.50E-04 | 0.01664 | 0.661 | 110.918 | 175.410 |
| N6amt1 | chr16:87354184-87368649 | 4.50E-04 | 0.01664 | 0.555 | 13.961 | 20.508 |
| Mrps16 | chr14:20391230-20393555 | 4.50E-04 | 0.01664 | 0.552 | 81.884 | 120.044 |
| Abcd3 | chr3:121758909-121815215 | 4.50E-04 | 0.01664 | 0.544 | 54.180 | 78.993 |
| Agpat1 | chr17:34605860-34615971 | 4.50E-04 | 0.01664 | 0.524 | 44.101 | 63.425 |
| 2310061I04Rik | chr17:35892676-35897378 | 4.50E-04 | 0.01664 | 0.521 | 34.952 | 50.148 |
| Fam210a | chr18:68260184-68300333 | 4.50E-04 | 0.01664 | 0.507 | 8.515 | 12.103 |
| Tmem14c | chr13:41016249-41022582 | 4.50E-04 | 0.01664 | 0.507 | 139.273 | 197.865 |
| Slc25a11 | chr11:70644026-70647039 | 4.50E-04 | 0.01664 | 0.493 | 63.194 | 88.943 |
| Rpl7l1 | chr17:46773906-46782656 | 4.50E-04 | 0.01664 | 0.481 | 26.419 | 36.872 |
| Bpnt1 | chr1:185332158-185357769 | 4.50E-04 | 0.01664 | 0.471 | 30.822 | 42.726 |
| Tln2 | chr9:67217084-67559703 | 4.50E-04 | 0.01664 | -0.484 | 8.727 | 6.240 |
| Hlf | chr11:90336534-90390917 | 4.50E-04 | 0.01664 | -0.507 | 9.946 | 7.000 |
| Clmn | chr12:104763113-104865076 | 4.50E-04 | 0.01664 | -0.514 | 8.684 | 6.081 |
| Tef | chr15:81802672-81826863 | 4.50E-04 | 0.01664 | -0.551 | 73.896 | 50.444 |
| Cyp2u1 | chr3:131290490-131303227 | 5.00E-04 | 0.01766 | 0.866 | 3.285 | 5.988 |
| Atg4d | chr9:21265284-21287969 | 5.00E-04 | 0.01766 | 0.768 | 9.795 | 16.685 |
| Gpd2 | chr2:57237677-57370719 | 5.00E-04 | 0.01766 | 0.625 | 58.863 | 90.763 |
| 2310039H08Rik | chr17:46772634-46773407 | 5.00E-04 | 0.01766 | 0.563 | 36.634 | 54.107 |
| Ndufv2 | chr17:66078794-66101559 | 5.00E-04 | 0.01766 | 0.533 | 160.422 | 232.151 |
| Usmg5 | chr19:47067747-47090625 | 5.00E-04 | 0.01766 | 0.523 | 311.623 | 447.663 |
| Tmem11 | chr11:60864451-60879038 | 5.00E-04 | 0.01766 | 0.510 | 41.537 | 59.136 |
| Rmdn3 | chr2:119136997-119157034 | 5.00E-04 | 0.01766 | 0.508 | 63.220 | 89.923 |

| | | | | | | |
|---|---|---|---|---|---|---|
| Ndufb6 | chr4:40270662-40279368 | 5.00E-04 | 0.01766 | 0.501 | 359.590 | 509.037 |
| Poldip2 | chr11:78512295-78522736 | 5.00E-04 | 0.01766 | 0.500 | 82.454 | 116.618 |
| Glrx5 | chr12:105032688-105040910 | 5.00E-04 | 0.01766 | 0.500 | 135.525 | 191.614 |
| 0610012G03Rik | chr16:31947050-31948521 | 5.00E-04 | 0.01766 | 0.486 | 36.346 | 50.916 |
| Stk40 | chr4:126103956-126141029 | 5.00E-04 | 0.01766 | 0.482 | 24.349 | 33.999 |
| Notch3 | chr17:32120892-32166852 | 5.00E-04 | 0.01766 | -0.486 | 11.417 | 8.150 |
| Net1 | chr13:3882017-3918220 | 5.00E-04 | 0.01766 | -0.531 | 73.357 | 50.754 |
| Tuba1a | chr15:98949846-98953501 | 5.00E-04 | 0.01766 | -0.561 | 163.160 | 110.603 |
| Soat1 | chr1:156428107-156474328 | 5.00E-04 | 0.01766 | -0.827 | 3.129 | 1.764 |
| Tmem82 | chr4:141614232-141618633 | 5.50E-04 | 0.01884 | 1.966 | 0.295 | 1.152 |
| Bhmt | chr13:93616890-93637758 | 5.50E-04 | 0.01884 | 1.564 | 1.244 | 3.679 |
| St3gal5 | chr6:72097607-72154570 | 5.50E-04 | 0.01884 | 0.777 | 5.114 | 8.762 |
| Cxadr | chr16:78301670-78359785 | 5.50E-04 | 0.01884 | 0.718 | 2.719 | 4.474 |
| Atf5 | chr7:44812255-44849079 | 5.50E-04 | 0.01884 | 0.692 | 98.436 | 159.080 |
| Ppargc1b | chr18:61298135-61400431 | 5.50E-04 | 0.01884 | 0.666 | 3.514 | 5.575 |
| Dlat | chr9:50634632-50659780 | 5.50E-04 | 0.01884 | 0.657 | 103.918 | 163.809 |
| Chchd4 | chr6:91464275-91473423 | 5.50E-04 | 0.01884 | 0.543 | 23.917 | 34.857 |
| Ube2f | chr1:91250318-91286025 | 5.50E-04 | 0.01884 | 0.525 | 24.869 | 35.785 |
| Srxn1 | chr2:152105523-152111376 | 5.50E-04 | 0.01884 | 0.523 | 19.835 | 28.504 |
| Dnajb9 | chr12:44205896-44210068 | 5.50E-04 | 0.01884 | 0.492 | 25.784 | 36.266 |
| Tmem56 | chr3:121202009-121263316 | 5.50E-04 | 0.01884 | -0.891 | 2.087 | 1.126 |
| Uck1 | chr2:32236682-32260105 | 6.00E-04 | 0.01999 | 0.613 | 91.823 | 140.408 |
| Mrps22 | chr9:98588729-98601679 | 6.00E-04 | 0.01999 | 0.542 | 24.599 | 35.815 |
| Atp2a2 | chr5:122453512-122502225 | 6.00E-04 | 0.01999 | 0.531 | 51.564 | 74.521 |
| Tbc1d20 | chr2:152293871-152312590 | 6.00E-04 | 0.01999 | 0.523 | 75.594 | 108.641 |
| Megf9 | chr4:70431926-70534928 | 6.00E-04 | 0.01999 | 0.514 | 10.383 | 14.829 |
| Fastk | chr5:24441039-24445235 | 6.00E-04 | 0.01999 | 0.513 | 51.131 | 72.959 |
| Ptcd3 | chr6:71880637-71908762 | 6.00E-04 | 0.01999 | 0.493 | 20.867 | 29.360 |

| | | | | | | |
|---|---|---|---|---|---|---|
| Gsto1 | chr19:47854988-47864788 | 6.00E-04 | 0.01999 | 0.491 | 55.249 | 77.633 |
| Mtor | chr4:148448581-148557685 | 6.00E-04 | 0.01999 | 0.488 | 12.610 | 17.689 |
| Tomm70a | chr16:57121713-57154530 | 6.00E-04 | 0.01999 | 0.473 | 28.717 | 39.846 |
| Clpx | chr9:65294259-65330658 | 6.00E-04 | 0.01999 | 0.472 | 42.919 | 59.546 |
| Cox5a | chr9:57521231-57532426 | 6.50E-04 | 0.02118 | 0.549 | 541.244 | 791.900 |
| Pmm2 | chr16:8637706-8657524 | 6.50E-04 | 0.02118 | 0.496 | 27.615 | 38.951 |
| Mcee | chr7:64392770-64412119 | 6.50E-04 | 0.02118 | 0.495 | 83.407 | 117.577 |
| Pak1ip1 | chr13:41001009-41013033 | 6.50E-04 | 0.02118 | 0.489 | 32.858 | 46.120 |
| Aatk | chr11:120007315-120047145 | 6.50E-04 | 0.02118 | 0.486 | 11.463 | 16.059 |
| C7 | chr15:4988761-5063773 | 6.50E-04 | 0.02118 | -0.500 | 57.523 | 40.669 |
| Nrep | chr18:33437018-33464029 | 6.50E-04 | 0.02118 | -0.517 | 47.083 | 32.911 |
| Neb | chr2:52136646-52338798 | 6.50E-04 | 0.02118 | -0.918 | 1.377 | 0.729 |
| Rdh9 | chr10:127776404-127792697 | 6.50E-04 | 0.02118 | -1.923 | 0.772 | 0.204 |
| Uqcrfs1 | chr13:30540311-30545316 | 7.00E-04 | 0.02227 | 0.608 | 333.784 | 508.657 |
| Gramd1b | chr9:40297906-40455764 | 7.00E-04 | 0.02227 | 0.593 | 6.517 | 9.828 |
| Deb1 | chr9:121710388-121712921 | 7.00E-04 | 0.02227 | 0.592 | 52.646 | 79.377 |
| Fastkd1 | chr2:69686823-69712606 | 7.00E-04 | 0.02227 | 0.583 | 7.232 | 10.831 |
| Cgrrf1 | chr14:46832243-46854190 | 7.00E-04 | 0.02227 | 0.541 | 32.959 | 47.962 |
| Gfm1 | chr3:67430114-67475068 | 7.00E-04 | 0.02227 | 0.505 | 38.343 | 54.425 |
| Irak2 | chr6:113638466-113695027 | 7.00E-04 | 0.02227 | 0.497 | 15.427 | 21.776 |
| Ptges2 | chr2:32395889-32402740 | 7.00E-04 | 0.02227 | 0.490 | 33.940 | 47.654 |
| Trdn | chr10:33083482-33476709 | 7.00E-04 | 0.02227 | -1.149 | 1.689 | 0.762 |
| Gm14085 | chr2:122484940-122528040 | 7.00E-04 | 0.02227 | -1.152 | 2.018 | 0.908 |
| Tuba4a | chr1:75210828-75219253 | 7.50E-04 | 0.02353 | 0.934 | 14.069 | 26.875 |
| Qrsl1 | chr10:43874189-43901736 | 7.50E-04 | 0.02353 | 0.696 | 5.814 | 9.419 |
| Hmgcs1 | chr13:119690350-119708260 | 7.50E-04 | 0.02353 | 0.630 | 109.935 | 170.154 |
| Ndufc2 | chr7:97400002-97407800 | 7.50E-04 | 0.02353 | 0.493 | 285.614 | 401.920 |
| Scn7a | chr2:66673425-66784910 | 7.50E-04 | 0.02353 | -0.505 | 35.021 | 24.671 |

| | | | | | | |
|---|---|---|---|---|---|---|
| Ly6d | chr15:74762055-74763567 | 7.50E-04 | 0.02353 | -1.846 | 5.886 | 1.637 |
| Pam16 | chr16:4616465-4624946 | 8.00E-04 | 0.02452 | 0.639 | 37.549 | 58.467 |
| Shb | chr4:45423275-45530828 | 8.00E-04 | 0.02452 | 0.546 | 10.266 | 14.991 |
| Mrpl53 | chr6:83101515-83109932 | 8.00E-04 | 0.02452 | 0.531 | 97.839 | 141.328 |
| Chchd3 | chr6:32792226-33060152 | 8.00E-04 | 0.02452 | 0.514 | 112.956 | 161.297 |
| Cdc34 | chr10:79682194-79688398 | 8.00E-04 | 0.02452 | 0.492 | 63.970 | 89.981 |
| Fam73b | chr2:30364232-30385519 | 8.00E-04 | 0.02452 | 0.490 | 51.008 | 71.622 |
| Mrpl30 | chr1:37890552-37898333 | 8.00E-04 | 0.02452 | 0.478 | 171.126 | 238.356 |
| Atp6v0e | chr17:26676395-26699646 | 8.00E-04 | 0.02452 | 0.470 | 187.983 | 260.465 |
| Azin2 | chr4:128930232-128962455 | 8.00E-04 | 0.02452 | -0.621 | 12.232 | 7.951 |
| Has3 | chr8:106870241-106882902 | 8.00E-04 | 0.02452 | -1.225 | 1.172 | 0.502 |
| Taco1 | chr11:106066106-106073612 | 8.50E-04 | 0.02559 | 0.547 | 17.501 | 25.561 |
| Pla2g7 | chr17:43568450-43612201 | 8.50E-04 | 0.02559 | 0.531 | 16.286 | 23.526 |
| Timm10b | chr7:105640539-105641845 | 8.50E-04 | 0.02559 | 0.509 | 37.597 | 53.516 |
| Pptc7 | chr5:122284397-122324281 | 8.50E-04 | 0.02559 | 0.491 | 11.161 | 15.681 |
| Mlx | chr11:101087289-101095435 | 8.50E-04 | 0.02559 | 0.486 | 24.999 | 35.004 |
| Chst1 | chr2:92599706-92615252 | 8.50E-04 | 0.02559 | -0.481 | 73.798 | 52.868 |
| Sema5a | chr15:32244812-32696341 | 8.50E-04 | 0.02559 | -0.562 | 2.622 | 1.776 |
| Mboat2 | chr12:24831598-24960299 | 8.50E-04 | 0.02559 | -1.538 | 1.243 | 0.428 |
| Cox8b | chr7:140898941-140900446 | 9.00E-04 | 0.02673 | 0.535 | 1104.400 | 1600.630 |
| Slc35f6 | chr5:30647935-30659729 | 9.00E-04 | 0.02673 | 0.488 | 12.422 | 17.422 |
| Coq7 | chr7:118509658-118533356 | 9.00E-04 | 0.02673 | 0.479 | 64.520 | 89.910 |
| Prelp | chr1:133910303-133921401 | 9.00E-04 | 0.02673 | -0.576 | 157.845 | 105.863 |
| Kcnk2 | chr1:189207929-189402273 | 9.00E-04 | 0.02673 | -0.627 | 6.789 | 4.397 |
| Xirp2 | chr2:67446001-67526606 | 9.00E-04 | 0.02673 | -1.002 | 1.105 | 0.552 |
| Coa7 | chr4:108328151-108340718 | 9.50E-04 | 0.02779 | 0.590 | 6.121 | 9.213 |
| Sdhaf4 | chr1:23995938-24005640 | 9.50E-04 | 0.02779 | 0.551 | 41.666 | 61.036 |
| Ric8b | chr10:84917612-85018337 | 9.50E-04 | 0.02779 | 0.481 | 7.828 | 10.926 |

| | | | | | | |
|---|---|---|---|---|---|---|
| Trib1 | chr15:59648653-59657099 | 9.50E-04 | 0.02779 | 0.470 | 12.819 | 17.760 |
| Tcn2 | chr11:3917077-3932078 | 9.50E-04 | 0.02779 | 0.466 | 60.597 | 83.673 |
| Mpz | chr1:171150712-171161123 | 9.50E-04 | 0.02779 | -0.466 | 50.719 | 36.713 |
| Gjb2 | chr14:57098601-57104702 | 9.50E-04 | 0.02779 | -1.772 | 1.573 | 0.461 |
| Angptl4 | chr17:33774899-33781575 | 1.00E-03 | 0.02875 | 0.595 | 131.196 | 198.172 |
| Grpel1 | chr5:36465184-36484285 | 1.00E-03 | 0.02875 | 0.566 | 38.412 | 56.854 |
| Pvrl2 | chr7:19716643-19749573 | 1.00E-03 | 0.02875 | 0.523 | 15.077 | 21.665 |
| Tmem126a | chr7:90450711-90457208 | 1.00E-03 | 0.02875 | 0.517 | 46.136 | 66.010 |
| Mrpl55 | chr11:59202485-59206135 | 1.00E-03 | 0.02875 | 0.504 | 38.367 | 54.412 |
| C1qbp | chr11:70970199-70983026 | 1.00E-03 | 0.02875 | 0.491 | 66.824 | 93.916 |
| Larp1b | chr3:40950630-40977793 | 1.00E-03 | 0.02875 | 0.473 | 25.221 | 35.014 |
| Tspan4 | chr7:141475235-141539857 | 1.00E-03 | 0.02875 | -0.583 | 71.417 | 47.685 |
| Dnajc25 | chr4:59003192-59023398 | 1.05E-03 | 0.02993 | 0.534 | 11.364 | 16.451 |
| Opa3 | chr7:19228388-19246817 | 1.05E-03 | 0.02993 | 0.517 | 12.891 | 18.447 |
| Oxnad1 | chr14:32085727-32103203 | 1.05E-03 | 0.02993 | 0.507 | 17.224 | 24.469 |
| Ankrd29 | chr18:12252356-12305720 | 1.05E-03 | 0.02993 | -0.900 | 3.626 | 1.943 |
| Poln | chr5:34007178-34169526 | 1.10E-03 | 0.03096 | 0.788 | 5.448 | 9.409 |
| Cmc2 | chr8:116888684-116921436 | 1.10E-03 | 0.03096 | 0.632 | 9.695 | 15.024 |
| Armt1 | chr10:4432604-4455140 | 1.10E-03 | 0.03096 | 0.573 | 7.701 | 11.456 |
| Abhd11 | chr5:135009151-135013157 | 1.10E-03 | 0.03096 | 0.475 | 53.722 | 74.645 |
| Ndrg1 | chr15:66929317-66969641 | 1.10E-03 | 0.03096 | -0.459 | 50.222 | 36.540 |
| Slc1a3 | chr15:8634123-8710807 | 1.10E-03 | 0.03096 | -0.514 | 56.887 | 39.846 |
| Bcl6 | chr16:23965051-23988612 | 1.15E-03 | 0.03203 | 0.507 | 16.246 | 23.079 |
| Kcmf1 | chr6:72841113-72899979 | 1.15E-03 | 0.03203 | 0.464 | 29.088 | 40.132 |
| Ndufb7 | chr8:83566757-83571623 | 1.15E-03 | 0.03203 | 0.460 | 292.469 | 402.202 |
| Pgls | chr8:71592183-71596267 | 1.15E-03 | 0.03203 | 0.453 | 93.785 | 128.356 |
| Vcan | chr13:89655309-89742512 | 1.15E-03 | 0.03203 | -0.661 | 4.049 | 2.561 |
| Tmem69 | chr4:116551527-116555943 | 1.20E-03 | 0.03281 | 0.526 | 10.262 | 14.781 |

| | | | | | | |
|---|---|---|---|---|---|---|
| Rilpl2 | chr5:124463264-124478235 | 1.20E-03 | 0.03281 | 0.498 | 26.048 | 36.798 |
| Loxl1 | chr9:58287722-58313212 | 1.20E-03 | 0.03281 | 0.482 | 48.396 | 67.571 |
| Pex14 | chr4:148960534-149099812 | 1.20E-03 | 0.03281 | 0.472 | 44.878 | 62.268 |
| Mrpl51 | chr6:125192199-125194392 | 1.20E-03 | 0.03281 | 0.472 | 33.829 | 46.927 |
| Dnajc15 | chr14:77826216-77874917 | 1.20E-03 | 0.03281 | 0.462 | 225.643 | 310.812 |
| Timm44 | chr8:4259730-4275905 | 1.20E-03 | 0.03281 | 0.452 | 40.993 | 56.079 |
| Fam132a | chr4:155962311-155966629 | 1.20E-03 | 0.03281 | 0.449 | 56.657 | 77.323 |
| Tjp1 | chr7:65296164-65371244 | 1.20E-03 | 0.03281 | -0.467 | 19.519 | 14.119 |
| Pwwp2b | chr7:139248481-139267253 | 1.25E-03 | 0.03363 | 0.642 | 4.977 | 7.767 |
| Pafah2 | chr4:134396319-134427412 | 1.25E-03 | 0.03363 | 0.541 | 8.043 | 11.705 |
| Pgp | chr17:24470472-24471596 | 1.25E-03 | 0.03363 | 0.484 | 39.053 | 54.630 |
| Tesk1 | chr4:43442276-43454626 | 1.25E-03 | 0.03363 | 0.475 | 16.992 | 23.615 |
| Mrpl37 | chr4:107055873-107066866 | 1.25E-03 | 0.03363 | 0.453 | 40.801 | 55.836 |
| Psmc4 | chr7:28041701-28050092 | 1.25E-03 | 0.03363 | 0.451 | 63.793 | 87.217 |
| Mtss1l | chr8:110721483-110741400 | 1.25E-03 | 0.03363 | -0.505 | 8.254 | 5.818 |
| Plxna3 | chrX:74329065-74344689 | 1.25E-03 | 0.03363 | -0.806 | 1.610 | 0.921 |
| Hoxa10 | chr6:52231196-52240854 | 1.30E-03 | 0.03436 | 0.587 | 7.221 | 10.845 |
| H2-Q10 | chr17:35470088-35474563 | 1.30E-03 | 0.03436 | 0.530 | 92.952 | 134.169 |
| Rpia | chr6:70765719-70792175 | 1.30E-03 | 0.03436 | 0.525 | 16.667 | 23.978 |
| Lzic | chr4:149485332-149496667 | 1.30E-03 | 0.03436 | 0.508 | 16.045 | 22.813 |
| Gbe1 | chr16:70313948-70569720 | 1.30E-03 | 0.03436 | 0.491 | 76.602 | 107.620 |
| Iars | chr13:49682129-49734267 | 1.30E-03 | 0.03436 | 0.462 | 8.897 | 12.254 |
| Mgat4b | chr11:50225334-50235103 | 1.30E-03 | 0.03436 | 0.438 | 63.091 | 85.457 |
| Aldh3a2 | chr11:61244751-61267186 | 1.30E-03 | 0.03436 | -0.483 | 28.581 | 20.446 |
| Pzp | chr6:128438756-128526720 | 1.30E-03 | 0.03436 | -1.139 | 2.581 | 1.172 |
| Atp6v1d | chr12:78842981-78861638 | 1.35E-03 | 0.03547 | 0.450 | 54.286 | 74.160 |
| Rasl11b | chr5:74195325-74199477 | 1.35E-03 | 0.03547 | -0.886 | 6.298 | 3.407 |
| Fcer2a | chr8:3681736-3694174 | 1.35E-03 | 0.03547 | -1.611 | 0.537 | 0.176 |

| | | | | | | |
|---|---|---|---|---|---|---|
| Npnt | chr3:132881744-132950291 | 1.40E-03 | 0.03657 | 1.064 | 0.844 | 1.764 |
| F3 | chr3:121723536-121735052 | 1.40E-03 | 0.03657 | 0.463 | 35.835 | 49.390 |
| Zfp697 | chr3:98382480-98431949 | 1.40E-03 | 0.03657 | -0.651 | 3.142 | 2.001 |
| Cth | chr3:157894247-157925063 | 1.50E-03 | 0.03873 | 0.587 | 8.126 | 12.207 |
| Tysnd1 | chr10:61695513-61702773 | 1.50E-03 | 0.03873 | 0.545 | 38.551 | 56.256 |
| Atp5o | chr16:91925222-91931630 | 1.50E-03 | 0.03873 | 0.496 | 420.917 | 593.799 |
| Nudt7 | chr8:114133573-114152312 | 1.50E-03 | 0.03873 | 0.483 | 128.285 | 179.251 |
| Mrpl35 | chr6:71812996-71823784 | 1.50E-03 | 0.03873 | 0.481 | 8.237 | 11.499 |
| Sec16b | chr1:157506795-157568424 | 1.50E-03 | 0.03873 | -0.490 | 7.937 | 5.651 |
| Ndufa6 | chr15:82350138-82354291 | 1.55E-03 | 0.03971 | 0.456 | 302.067 | 414.474 |
| Tmem43 | chr6:91473750-91488458 | 1.55E-03 | 0.03971 | -0.494 | 120.180 | 85.342 |
| Mov10 | chr3:104794833-104818563 | 1.55E-03 | 0.03971 | -0.544 | 7.513 | 5.153 |
| Caprin2 | chr6:148842491-148896237 | 1.55E-03 | 0.03971 | -0.723 | 3.759 | 2.277 |
| Fahd1 | chr17:24848895-24850302 | 1.60E-03 | 0.04060 | 0.505 | 19.130 | 27.149 |
| Egln1 | chr8:124908586-124949254 | 1.60E-03 | 0.04060 | 0.464 | 43.321 | 59.766 |
| Hba-a1,Hba-a2 | chr11:32283671-32284493 | 1.60E-03 | 0.04060 | 0.457 | 299.015 | 416.582 |
| Timm50 | chr7:28305825-28312046 | 1.60E-03 | 0.04060 | 0.445 | 60.609 | 82.507 |
| Fgb | chr3:83042304-83049790 | 1.60E-03 | 0.04060 | -0.988 | 6.174 | 3.114 |
| Endog | chr2:30171523-30178459 | 1.65E-03 | 0.04140 | 0.584 | 35.643 | 53.418 |
| Trmu | chr15:85879326-85897393 | 1.65E-03 | 0.04140 | 0.536 | 13.511 | 19.586 |
| Ncbp1 | chr4:46138510-46172402 | 1.65E-03 | 0.04140 | 0.438 | 17.040 | 23.091 |
| Mturn | chr6:54681623-54703855 | 1.65E-03 | 0.04140 | -0.450 | 23.871 | 17.477 |
| Pcdh20 | chr14:88464746-88471396 | 1.65E-03 | 0.04140 | -0.775 | 2.307 | 1.348 |
| Tacstd2 | chr6:67534058-67535822 | 1.65E-03 | 0.04140 | -2.186 | 0.938 | 0.206 |
| Psmb5 | chr14:54614119-54617995 | 1.70E-03 | 0.04242 | 0.481 | 64.372 | 89.856 |
| Uqcr10 | chr11:4701967-4704344 | 1.70E-03 | 0.04242 | 0.465 | 689.144 | 951.530 |
| Efs | chr14:54916542-54926788 | 1.70E-03 | 0.04242 | -0.477 | 12.061 | 8.664 |
| Bcl3 | chr7:19808461-19822755 | 1.75E-03 | 0.04335 | 0.658 | 5.611 | 8.853 |

| | | | | | | |
|---|---|---|---|---|---|---|
| Cox11 | chr11:90638183-90687601 | 1.75E-03 | 0.04335 | 0.600 | 9.430 | 14.295 |
| Rbp7 | chr4:149449701-149454968 | 1.75E-03 | 0.04335 | 0.533 | 58.644 | 84.838 |
| Fabp5 | chr3:10012584-10016610 | 1.75E-03 | 0.04335 | 0.476 | 240.069 | 333.888 |
| Narfl | chr17:25773775-25785586 | 1.80E-03 | 0.04418 | 0.543 | 12.630 | 18.402 |
| Pfkl | chr10:77986948-78009796 | 1.80E-03 | 0.04418 | 0.540 | 72.746 | 105.801 |
| Cpt1a | chr19:3323300-3385733 | 1.80E-03 | 0.04418 | -0.431 | 22.627 | 16.787 |
| Thbd | chr2:148404470-148408188 | 1.80E-03 | 0.04418 | -0.482 | 74.741 | 53.512 |
| Acot5 | chr12:84069324-84076019 | 1.80E-03 | 0.04418 | -1.712 | 2.105 | 0.643 |
| Ppp1r3d | chr2:178411205-178414463 | 1.85E-03 | 0.04524 | 0.464 | 10.257 | 14.152 |
| Gsk3a | chr7:25228258-25237851 | 1.85E-03 | 0.04524 | 0.451 | 38.211 | 52.244 |
| Mdh1 | chr11:21556691-21571934 | 1.90E-03 | 0.04629 | 0.682 | 403.091 | 646.889 |
| Atp6v0e2 | chr6:48537568-48541800 | 1.90E-03 | 0.04629 | -0.599 | 12.346 | 8.153 |
| Dolk | chr2:30284228-30286354 | 1.95E-03 | 0.04734 | 0.425 | 49.735 | 66.775 |
| Clcn2 | chr16:20695056-20716636 | 1.95E-03 | 0.04734 | -0.715 | 4.187 | 2.551 |
| Prdx3 | chr19:60864065-60874538 | 2.00E-03 | 0.04820 | 0.461 | 147.820 | 203.413 |
| Slc25a22 | chr7:141429748-141437874 | 2.00E-03 | 0.04820 | 0.439 | 52.649 | 71.386 |
| Slco2b1 | chr7:99657803-99711340 | 2.00E-03 | 0.04820 | -0.435 | 25.685 | 19.001 |
| Ift122 | chr6:115853527-115926699 | 2.00E-03 | 0.04820 | -0.689 | 3.509 | 2.177 |
| Ruvbl2 | chr7:45421897-45434464 | 2.05E-03 | 0.04880 | 0.469 | 22.277 | 30.831 |
| Mrpl4 | chr9:21002736-21008837 | 2.05E-03 | 0.04880 | 0.439 | 46.078 | 62.448 |
| Rcl1 | chr19:29101374-29143843 | 2.05E-03 | 0.04880 | 0.438 | 27.353 | 37.047 |
| Cpped1 | chr16:11803720-11930594 | 2.05E-03 | 0.04880 | 0.422 | 28.568 | 38.276 |
| Emc3 | chr6:113514886-113531638 | 2.05E-03 | 0.04880 | 0.418 | 56.235 | 75.110 |
| Agps | chr2:75832176-75931350 | 2.05E-03 | 0.04880 | -0.504 | 11.304 | 7.969 |
| Mt2 | chr8:94172617-94173567 | 2.05E-03 | 0.04880 | -0.513 | 90.278 | 63.278 |
| Lipt1 | chr1:37872205-37876298 | 2.10E-03 | 0.04955 | 0.556 | 11.441 | 16.826 |
| Cnst | chr1:179546528-179627473 | 2.10E-03 | 0.04955 | 0.440 | 19.311 | 26.204 |
| 1110008P14Rik | chr2:32379100-32381915 | 2.10E-03 | 0.04955 | 0.430 | 143.908 | 193.885 |

| | | | | | | |
|---|---|---|---|---|---|---|
| Mfng | chr15:78755882-78773445 | 2.10E-03 | 0.04955 | -0.444 | 103.564 | 76.135 |
| Aqp1 | chr6:55336298-55348555 | 2.10E-03 | 0.04955 | -0.510 | 184.353 | 129.416 |

**Table S3**: Genes significantly differentially expressed between RT and 4°C in Rbp$^{-/-}$ mice.

| gene | locus | raw p-value | Benjamini Hochbreg corrected p-value | mean fold change 4°C/RT (log2) | KO (RT) | KO (4°C) |
|---|---|---|---|---|---|---|
| Perm1 | chr4:156215926-156221307 | 5.00E-05 | 0.00335 | 3.416 | 0.283 | 3.025 |
| Dio2 | chr12:90724551-90738438 | 5.00E-05 | 0.00335 | 3.318 | 1.451 | 14.474 |
| Cyp2b10 | chr7:25897657-25926624 | 5.00E-05 | 0.00335 | 2.481 | 0.652 | 3.638 |
| Gyk | chrX:85701936-85776819 | 5.00E-05 | 0.00335 | 2.417 | 6.891 | 36.807 |
| Tmem82 | chr4:141614232-141618633 | 5.00E-05 | 0.00335 | 2.023 | 0.333 | 1.352 |
| Slc25a34 | chr4:141618824-141623834 | 5.00E-05 | 0.00335 | 2.013 | 2.109 | 8.512 |
| 9030619P08Rik | chr15:75427605-75431829 | 5.00E-05 | 0.00335 | 1.900 | 1.206 | 4.500 |
| Fabp3 | chr4:130308777-130315463 | 5.00E-05 | 0.00335 | 1.889 | 21.313 | 78.916 |
| Cyp1a1 | chr9:57687927-57703824 | 5.00E-05 | 0.00335 | 1.761 | 1.430 | 4.844 |
| Dhrs9 | chr2:69380461-69403086 | 5.00E-05 | 0.00335 | 1.747 | 0.519 | 1.743 |
| Elovl3 | chr19:46131898-46148325 | 5.00E-05 | 0.00335 | 1.718 | 26.665 | 87.704 |
| Cyp2f2 | chr7:27119954-27133660 | 5.00E-05 | 0.00335 | 1.473 | 88.635 | 246.034 |
| Cdkn1a | chr17:29090985-29100722 | 5.00E-05 | 0.00335 | 1.463 | 16.229 | 44.742 |
| Ankrd9 | chr12:110975352-110979021 | 5.00E-05 | 0.00335 | 1.438 | 4.658 | 12.625 |
| Serpina1b | chr12:103728155-103738189 | 5.00E-05 | 0.00335 | 1.438 | 30.828 | 83.507 |
| Fam110c | chr12:31073967-31079940 | 5.00E-05 | 0.00335 | 1.396 | 3.863 | 10.165 |
| Gmpr | chr13:45507443-45546386 | 5.00E-05 | 0.00335 | 1.378 | 8.749 | 22.737 |
| Sugct | chr13:16857474-17694765 | 5.00E-05 | 0.00335 | 1.298 | 2.610 | 6.417 |
| Wfdc21 | chr11:83746939-83752646 | 5.00E-05 | 0.00335 | 1.232 | 141.996 | 333.492 |
| Bcl6 | chr16:23965051-23988612 | 5.00E-05 | 0.00335 | 1.203 | 18.342 | 42.224 |
| Ppargc1a | chr5:51454248-51553921 | 5.00E-05 | 0.00335 | 1.188 | 2.718 | 6.193 |
| Ldlr | chr9:21723575-21749918 | 5.00E-05 | 0.00335 | 1.088 | 10.628 | 22.597 |

| | | | | | | |
|---|---|---|---|---|---|---|
| Nr4a1 | chr15:101266845-101274794 | 5.00E-05 | 0.00335 | 1.074 | 2.480 | 5.223 |
| Apex2 | chrX:150571506-150588149 | 5.00E-05 | 0.00335 | 1.050 | 5.836 | 12.081 |
| Slc6a13 | chr6:121300295-121337718 | 5.00E-05 | 0.00335 | 1.048 | 10.700 | 22.122 |
| Trhde | chr10:114398820-114801370 | 5.00E-05 | 0.00335 | 1.020 | 1.243 | 2.520 |
| Dusp10 | chr1:184034460-184075636 | 5.00E-05 | 0.00335 | 1.000 | 9.255 | 18.513 |
| Paqr9 | chr9:95559816-95562121 | 5.00E-05 | 0.00335 | 0.978 | 36.396 | 71.714 |
| Slc25a25 | chr2:32414486-32455476 | 5.00E-05 | 0.00335 | 0.975 | 14.100 | 27.719 |
| Gsta3 | chr1:21240584-21265575 | 5.00E-05 | 0.00335 | 0.966 | 107.814 | 210.625 |
| Hfe | chr13:23703840-23710811 | 5.00E-05 | 0.00335 | 0.952 | 34.665 | 67.077 |
| Got1 | chr19:43499752-43524605 | 5.00E-05 | 0.00335 | 0.927 | 7.347 | 13.967 |
| G0s2 | chr1:193272159-193273188 | 5.00E-05 | 0.00335 | 0.925 | 320.466 | 608.324 |
| Ndufaf4 | chr4:24898082-24905001 | 5.00E-05 | 0.00335 | 0.924 | 8.571 | 16.268 |
| Mapk6 | chr9:75386781-75410016 | 5.00E-05 | 0.00335 | 0.908 | 25.813 | 48.445 |
| Syt12 | chr19:4425456-4477143 | 5.00E-05 | 0.00335 | 0.905 | 16.917 | 31.671 |
| Kcnj15 | chr16:95257557-95300258 | 5.00E-05 | 0.00335 | 0.900 | 22.012 | 41.064 |
| Tmed5 | chr5:108121646-108132591 | 5.00E-05 | 0.00335 | 0.896 | 33.325 | 62.019 |
| Tubb4b | chr2:25218744-25224702 | 5.00E-05 | 0.00335 | 0.893 | 75.016 | 139.348 |
| Atl2 | chr17:79848389-79896123 | 5.00E-05 | 0.00335 | 0.888 | 32.206 | 59.613 |
| Gadd45g | chr13:51846674-51848474 | 5.00E-05 | 0.00335 | 0.886 | 32.505 | 60.064 |
| Ccrn4l | chr3:51224446-51251654 | 5.00E-05 | 0.00335 | 0.882 | 10.922 | 20.129 |
| Cth | chr3:157894247-157925063 | 5.00E-05 | 0.00335 | 0.862 | 8.435 | 15.337 |
| Prodh | chr16:18071725-18089190 | 5.00E-05 | 0.00335 | 0.856 | 19.861 | 35.961 |
| Zfp503 | chr14:21983961-21989601 | 5.00E-05 | 0.00335 | 0.845 | 12.497 | 22.446 |
| Traf4 | chr11:78158422-78165550 | 5.00E-05 | 0.00335 | 0.843 | 11.742 | 21.061 |
| Hccs | chrX:169311530-169320343 | 5.00E-05 | 0.00335 | 0.839 | 10.725 | 19.189 |
| Nnmt | chr9:48591889-48605077 | 5.00E-05 | 0.00335 | 0.837 | 175.675 | 313.751 |
| Arxes1 | chrX:136033366-136034946 | 5.00E-05 | 0.00335 | 0.836 | 38.527 | 68.780 |
| Igfbp3 | chr11:7206090-7213923 | 5.00E-05 | 0.00335 | 0.836 | 12.454 | 22.225 |

| | | | | | | |
|---|---|---|---|---|---|---|
| Adam12 | chr7:133883198-134225097 | 5.00E-05 | 0.00335 | 0.835 | 5.152 | 9.189 |
| Ptch2 | chr4:117096355-117114831 | 5.00E-05 | 0.00335 | 0.813 | 5.804 | 10.196 |
| Slc25a17 | chr15:81318920-81360765 | 5.00E-05 | 0.00335 | 0.772 | 58.446 | 99.794 |
| Dusp1 | chr17:26505590-26508472 | 5.00E-05 | 0.00335 | 0.767 | 19.923 | 33.912 |
| Tpcn2 | chr7:145253922-145283927 | 5.00E-05 | 0.00335 | 0.761 | 4.791 | 8.121 |
| Ciapin1 | chr8:94819817-94838340 | 5.00E-05 | 0.00335 | 0.744 | 9.867 | 16.522 |
| Tatdn2 | chr6:113697498-113711068 | 5.00E-05 | 0.00335 | 0.739 | 16.206 | 27.054 |
| Ppp1r3d | chr2:178411205-178414463 | 5.00E-05 | 0.00335 | 0.739 | 8.473 | 14.140 |
| Mid1ip1 | chrX:10717364-10719702 | 5.00E-05 | 0.00335 | 0.734 | 61.157 | 101.750 |
| Ifrd1 | chr12:40203128-40223189 | 5.00E-05 | 0.00335 | 0.734 | 65.821 | 109.506 |
| Hdhd3 | chr4:62499053-62502200 | 5.00E-05 | 0.00335 | 0.734 | 21.914 | 36.445 |
| Inmt | chr6:55170626-55174990 | 5.00E-05 | 0.00335 | 0.731 | 130.134 | 216.034 |
| Slmo2 | chr2:174465090-174472941 | 5.00E-05 | 0.00335 | 0.726 | 51.258 | 84.811 |
| Fam213b | chr4:154896429-154899043 | 5.00E-05 | 0.00335 | 0.724 | 127.716 | 210.993 |
| 5730508B09Rik | chr3:127869687-127896323 | 5.00E-05 | 0.00335 | 0.722 | 22.559 | 37.200 |
| 3110043O21Rik | chr4:35191281-35225880 | 5.00E-05 | 0.00335 | 0.716 | 35.020 | 57.533 |
| Timm9 | chr12:71111427-71136675 | 5.00E-05 | 0.00335 | 0.712 | 26.301 | 43.095 |
| Arxes2 | chrX:135993819-135995359 | 5.00E-05 | 0.00335 | 0.711 | 135.690 | 222.158 |
| Ppif | chr14:25694169-25701282 | 5.00E-05 | 0.00335 | 0.705 | 24.987 | 40.744 |
| Mocs2 | chr13:114818236-114829420 | 5.00E-05 | 0.00335 | 0.703 | 75.747 | 123.274 |
| Gmps | chr3:63976142-64019078 | 5.00E-05 | 0.00335 | 0.680 | 8.246 | 13.209 |
| Car13 | chr3:14641726-14663002 | 5.00E-05 | 0.00335 | 0.670 | 14.834 | 23.607 |
| Rbm3 | chrX:8138974-8147963 | 5.00E-05 | 0.00335 | 0.657 | 61.805 | 97.462 |
| Akirin1 | chr4:123735194-123750299 | 5.00E-05 | 0.00335 | 0.657 | 20.130 | 31.731 |
| Nabp1 | chr1:51469487-51478399 | 5.00E-05 | 0.00335 | 0.653 | 36.309 | 57.109 |
| Pla2g12a | chr3:129878605-129895825 | 5.00E-05 | 0.00335 | 0.651 | 46.019 | 72.270 |
| Rcl1 | chr19:29101374-29143843 | 5.00E-05 | 0.00335 | 0.651 | 25.234 | 39.624 |
| Slc38a2 | chr15:96687391-96699698 | 5.00E-05 | 0.00335 | 0.602 | 14.185 | 21.530 |

| | | | | | | |
|---|---|---|---|---|---|---|
| Scn1b | chr7:31116523-31126945 | 5.00E-05 | 0.00335 | -0.586 | 46.721 | 31.124 |
| Atp8a1 | chr5:67618138-67847431 | 5.00E-05 | 0.00335 | -0.646 | 5.530 | 3.534 |
| Thbs2 | chr17:14665499-14694262 | 5.00E-05 | 0.00335 | -0.668 | 14.606 | 9.194 |
| Podn | chr4:108014792-108032090 | 5.00E-05 | 0.00335 | -0.686 | 58.732 | 36.494 |
| Aplnr | chr2:85136359-85139923 | 5.00E-05 | 0.00335 | -0.726 | 13.042 | 7.887 |
| Tln2 | chr9:67217084-67559703 | 5.00E-05 | 0.00335 | -0.737 | 8.532 | 5.119 |
| Fmo5 | chr3:97628803-97655287 | 5.00E-05 | 0.00335 | -0.766 | 12.119 | 7.127 |
| Fam20c | chr5:138755080-138810063 | 5.00E-05 | 0.00335 | -0.775 | 17.729 | 10.362 |
| Slc43a1 | chr2:84839407-84863586 | 5.00E-05 | 0.00335 | -0.807 | 21.174 | 12.102 |
| Fbn1 | chr2:125300593-125506438 | 5.00E-05 | 0.00335 | -0.809 | 23.552 | 13.442 |
| Grik5 | chr7:25009850-25072369 | 5.00E-05 | 0.00335 | -0.817 | 4.567 | 2.591 |
| Aldh1a3 | chr7:66390892-66427477 | 5.00E-05 | 0.00335 | -0.820 | 7.990 | 4.527 |
| Rcn3 | chr7:45082913-45092213 | 5.00E-05 | 0.00335 | -0.821 | 48.627 | 27.530 |
| Trp53i11 | chr2:93187583-93201757 | 5.00E-05 | 0.00335 | -0.825 | 17.392 | 9.818 |
| Synpo2 | chr3:123076518-123236149 | 5.00E-05 | 0.00335 | -0.829 | 4.326 | 2.436 |
| Tppp | chr13:74009418-74035753 | 5.00E-05 | 0.00335 | -0.834 | 8.973 | 5.034 |
| Rnf144b | chr13:47122719-47247991 | 5.00E-05 | 0.00335 | -0.838 | 6.351 | 3.553 |
| Cpe | chr8:64592550-64693040 | 5.00E-05 | 0.00335 | -0.879 | 39.158 | 21.292 |
| Kit | chr5:75574986-75656721 | 5.00E-05 | 0.00335 | -0.891 | 5.176 | 2.792 |
| Dhcr24 | chr4:106561037-106589113 | 5.00E-05 | 0.00335 | -0.897 | 13.157 | 7.063 |
| Col14a1 | chr15:55307749-55520803 | 5.00E-05 | 0.00335 | -0.900 | 13.042 | 6.987 |
| Oxtr | chr6:112473683-112489808 | 5.00E-05 | 0.00335 | -0.902 | 9.281 | 4.967 |
| Mapk8ip1 | chr2:92383675-92401263 | 5.00E-05 | 0.00335 | -0.953 | 15.095 | 7.800 |
| Serpina3k | chr12:104338485-104345739 | 5.00E-05 | 0.00335 | -0.954 | 21.695 | 11.201 |
| Bmyc | chr2:25706878-25707719 | 5.00E-05 | 0.00335 | -0.954 | 33.506 | 17.296 |
| Syne3 | chr12:104929932-105009809 | 5.00E-05 | 0.00335 | -0.965 | 5.871 | 3.007 |
| Tfr2 | chr5:137569839-137588081 | 5.00E-05 | 0.00335 | -0.966 | 4.855 | 2.486 |
| Lctl | chr9:64117146-64172931 | 5.00E-05 | 0.00335 | -0.969 | 24.355 | 12.443 |

| | | | | | | |
|---|---|---|---|---|---|---|
| Smpd3 | chr8:106252547-106337988 | 5.00E-05 | 0.00335 | -0.988 | 8.301 | 4.184 |
| Rian | chr12:109603944-109661711 | 5.00E-05 | 0.00335 | -0.999 | 4.744 | 2.373 |
| Sfrp5 | chr19:42197970-42202252 | 5.00E-05 | 0.00335 | -1.017 | 11.731 | 5.796 |
| Tnnc2 | chr2:164777161-164779734 | 5.00E-05 | 0.00335 | -1.022 | 102.689 | 50.560 |
| Casq1 | chr1:172209893-172219895 | 5.00E-05 | 0.00335 | -1.028 | 9.961 | 4.885 |
| Fads2 | chr19:10064163-10101503 | 5.00E-05 | 0.00335 | -1.033 | 14.814 | 7.240 |
| Rorc | chr3:94372793-94398274 | 5.00E-05 | 0.00335 | -1.034 | 13.809 | 6.745 |
| Mup3 | chr4:62083475-62087312 | 5.00E-05 | 0.00335 | -1.041 | 36.261 | 17.626 |
| Mybpc2 | chr7:44501698-44524669 | 5.00E-05 | 0.00335 | -1.044 | 9.009 | 4.368 |
| Mylpf | chr7:127211607-127214287 | 5.00E-05 | 0.00335 | -1.048 | 68.332 | 33.038 |
| Ttn | chr2:76703983-76982547 | 5.00E-05 | 0.00335 | -1.067 | 0.606 | 0.289 |
| Pvalb | chr15:78191117-78206351 | 5.00E-05 | 0.00335 | -1.086 | 70.387 | 33.166 |
| Fn3k | chr11:121434952-121450490 | 5.00E-05 | 0.00335 | -1.091 | 13.174 | 6.184 |
| Ddo | chr10:40630010-40649931 | 5.00E-05 | 0.00335 | -1.091 | 11.391 | 5.346 |
| Aspg | chr12:112106682-112127573 | 5.00E-05 | 0.00335 | -1.095 | 20.752 | 9.718 |
| Actn3 | chr19:4861215-4877909 | 5.00E-05 | 0.00335 | -1.095 | 25.878 | 12.112 |
| Neb | chr2:52136646-52338798 | 5.00E-05 | 0.00335 | -1.099 | 0.900 | 0.420 |
| Cish | chr9:107296688-107301961 | 5.00E-05 | 0.00335 | -1.102 | 12.123 | 5.646 |
| Celsr2 | chr3:108390847-108415494 | 5.00E-05 | 0.00335 | -1.118 | 3.755 | 1.730 |
| Lama3 | chr18:12334023-12583012 | 5.00E-05 | 0.00335 | -1.125 | 0.853 | 0.391 |
| Pth1r | chr9:110722084-110747145 | 5.00E-05 | 0.00335 | -1.130 | 26.533 | 12.126 |
| Fam167a | chr14:63436393-63465502 | 5.00E-05 | 0.00335 | -1.205 | 1.956 | 0.848 |
| Prune2 | chr19:16956117-17223932 | 5.00E-05 | 0.00335 | -1.208 | 4.334 | 1.876 |
| Acsm3 | chr7:119760922-119794058 | 5.00E-05 | 0.00335 | -1.251 | 34.288 | 14.407 |
| Apob | chr12:7977676-8016839 | 5.00E-05 | 0.00335 | -1.251 | 1.100 | 0.462 |
| Hdac11 | chr6:91156814-91174683 | 5.00E-05 | 0.00335 | -1.253 | 3.527 | 1.480 |
| Nav2 | chr7:48959072-49610088 | 5.00E-05 | 0.00335 | -1.261 | 0.611 | 0.255 |
| Vnn1 | chr10:23894687-23905343 | 5.00E-05 | 0.00335 | -1.299 | 4.659 | 1.893 |

| | | | | | | |
|---|---|---|---|---|---|---|
| Tmem56 | chr3:121202009-121263316 | 5.00E-05 | 0.00335 | -1.302 | 2.172 | 0.881 |
| Cyp3a11 | chr5:145854606-145879854 | 5.00E-05 | 0.00335 | -1.307 | 11.938 | 4.825 |
| Hmgcs2 | chr3:98280430-98310738 | 5.00E-05 | 0.00335 | -1.336 | 16.451 | 6.518 |
| Ttr | chr18:20665249-20674326 | 5.00E-05 | 0.00335 | -1.336 | 28.496 | 11.290 |
| Ampd1 | chr3:103074013-103099720 | 5.00E-05 | 0.00335 | -1.356 | 3.403 | 1.330 |
| Golga7b | chr19:42247577-42270348 | 5.00E-05 | 0.00335 | -1.381 | 8.380 | 3.218 |
| Mycl | chr4:122995930-123002480 | 5.00E-05 | 0.00335 | -1.433 | 3.848 | 1.426 |
| Slc38a3 | chr9:107651154-107668968 | 5.00E-05 | 0.00335 | -1.472 | 4.036 | 1.455 |
| Gc | chr5:89417510-89457898 | 5.00E-05 | 0.00335 | -1.489 | 15.287 | 5.447 |
| Ces3a | chr8:105048598-105058414 | 5.00E-05 | 0.00335 | -1.507 | 3.856 | 1.356 |
| Ambp | chr4:63143278-63154142 | 5.00E-05 | 0.00335 | -1.522 | 7.331 | 2.553 |
| Peg3 | chr7:6705959-6730419 | 5.00E-05 | 0.00335 | -1.549 | 4.671 | 1.597 |
| Krt79 | chr15:101929331-101940324 | 5.00E-05 | 0.00335 | -1.555 | 19.226 | 6.541 |
| Ntrk3 | chr7:78192113-78577838 | 5.00E-05 | 0.00335 | -1.601 | 15.028 | 4.954 |
| Uox | chr3:146581372-146631483 | 5.00E-05 | 0.00335 | -1.644 | 5.483 | 1.755 |
| Apoa1 | chr9:46228629-46230469 | 5.00E-05 | 0.00335 | -1.645 | 45.988 | 14.707 |
| Kng1 | chr16:23058299-23082078 | 5.00E-05 | 0.00335 | -1.656 | 4.602 | 1.461 |
| Plg | chr17:12378608-12419384 | 5.00E-05 | 0.00335 | -1.674 | 3.573 | 1.119 |
| Mug1 | chr6:121838540-121889057 | 5.00E-05 | 0.00335 | -1.702 | 2.984 | 0.917 |
| Slc2a5 | chr4:150119343-150144168 | 5.00E-05 | 0.00335 | -1.715 | 22.911 | 6.979 |
| Thbs1 | chr2:118111921-118127133 | 5.00E-05 | 0.00335 | -1.810 | 4.470 | 1.275 |
| Krtdap | chr7:30787904-30791083 | 5.00E-05 | 0.00335 | -1.900 | 24.683 | 6.614 |
| Atp1a3 | chr7:24978168-25005937 | 5.00E-05 | 0.00335 | -1.902 | 24.897 | 6.661 |
| Fgb | chr3:83042304-83049790 | 5.00E-05 | 0.00335 | -1.943 | 9.807 | 2.551 |
| Ddah1 | chr3:145758691-145894277 | 5.00E-05 | 0.00335 | -1.949 | 3.459 | 0.896 |
| Fcgbp | chr7:28071235-28120864 | 5.00E-05 | 0.00335 | -2.000 | 0.641 | 0.160 |
| Fgg | chr3:83007895-83015049 | 5.00E-05 | 0.00335 | -2.011 | 9.096 | 2.257 |
| Gbp11 | chr5:105323025-105346476 | 5.00E-05 | 0.00335 | -2.027 | 5.390 | 1.322 |

| | | | | | | |
|---|---|---|---|---|---|---|
| Pzp | chr6:128438756-128526720 | 5.00E-05 | 0.00335 | -2.028 | 3.825 | 0.938 |
| Fga | chr3:83026152-83033617 | 5.00E-05 | 0.00335 | -2.102 | 6.431 | 1.498 |
| Igfbp2 | chr1:72824479-72852471 | 5.00E-05 | 0.00335 | -2.125 | 5.267 | 1.208 |
| Pkp3 | chr7:141078228-141090510 | 5.00E-05 | 0.00335 | -2.183 | 1.356 | 0.298 |
| Otof | chr5:30367065-30461932 | 5.00E-05 | 0.00335 | -2.232 | 0.866 | 0.184 |
| Kcng4 | chr8:119623853-119635680 | 5.00E-05 | 0.00335 | -2.295 | 1.129 | 0.230 |
| Fam84a | chr12:14147597-14152038 | 5.00E-05 | 0.00335 | -2.394 | 1.743 | 0.332 |
| Mmd2 | chr5:142563480-142608752 | 5.00E-05 | 0.00335 | -2.422 | 1.043 | 0.195 |
| Foxq1 | chr13:31558169-31560974 | 5.00E-05 | 0.00335 | -2.449 | 0.856 | 0.157 |
| AA467197 | chr2:122637886-122641076 | 5.00E-05 | 0.00335 | -2.562 | 10.824 | 1.833 |
| Rdh9 | chr10:127776404-127792697 | 5.00E-05 | 0.00335 | -2.596 | 0.746 | 0.123 |
| Elovl4 | chr9:83778691-83806305 | 5.00E-05 | 0.00335 | -2.750 | 1.051 | 0.156 |
| Fabp1 | chr6:71199887-71205023 | 5.00E-05 | 0.00335 | -3.124 | 56.505 | 6.482 |
| Trim29 | chr9:43310762-43336125 | 5.00E-05 | 0.00335 | -3.335 | 2.891 | 0.287 |
| Ly6d | chr15:74762055-74763567 | 5.00E-05 | 0.00335 | -3.364 | 14.491 | 1.408 |
| Tacstd2 | chr6:67534058-67535822 | 5.00E-05 | 0.00335 | -3.430 | 1.697 | 0.157 |
| Perp | chr10:18845070-18857072 | 5.00E-05 | 0.00335 | -3.643 | 7.165 | 0.574 |
| Ehf | chr2:103263432-103303196 | 5.00E-05 | 0.00335 | -3.821 | 1.052 | 0.074 |
| S100a14 | chr3:90526848-90528835 | 5.00E-05 | 0.00335 | -3.958 | 3.289 | 0.212 |
| Calml3 | chr13:3802892-3804318 | 5.00E-05 | 0.00335 | -3.960 | 6.865 | 0.441 |
| Pkp1 | chr1:135871394-135919024 | 5.00E-05 | 0.00335 | -4.545 | 1.839 | 0.079 |
| Asprv1 | chr6:86628173-86629704 | 5.00E-05 | 0.00335 | -4.553 | 5.383 | 0.229 |
| Gbp2b | chr3:142594846-142619176 | 5.00E-05 | 0.00335 | -4.641 | 5.074 | 0.203 |
| Klk10 | chr7:43781053-43785410 | 5.00E-05 | 0.00335 | -5.361 | 17.513 | 0.426 |
| Krt14 | chr11:100203161-100207510 | 5.00E-05 | 0.00335 | -5.862 | 13.655 | 0.235 |
| Awat2 | chrX:100402221-100442717 | 5.00E-05 | 0.00335 | -6.062 | 15.732 | 0.235 |
| Dsp | chr13:38151293-38198577 | 5.00E-05 | 0.00335 | -6.175 | 3.191 | 0.044 |
| Cst6 | chr19:5344704-5349574 | 5.00E-05 | 0.00335 | -8.806 | 39.099 | 0.087 |

| | | | | | | |
|---|---|---|---|---|---|---|
| Dfna5 | chr6:50207402-50261769 | 1.00E-04 | 0.00601 | 0.897 | 4.697 | 8.745 |
| Them4 | chr3:94310131-94332532 | 1.00E-04 | 0.00601 | 0.865 | 4.779 | 8.705 |
| Adcy3 | chr12:4133396-4240123 | 1.00E-04 | 0.00601 | 0.857 | 15.816 | 28.652 |
| Spon1 | chr7:113765997-114043375 | 1.00E-04 | 0.00601 | 0.722 | 3.594 | 5.927 |
| Tubb5 | chr17:35833919-35838301 | 1.00E-04 | 0.00601 | 0.719 | 64.428 | 106.051 |
| Aspa | chr11:73304987-73324637 | 1.00E-04 | 0.00601 | 0.702 | 50.953 | 82.896 |
| Slc3a2 | chr19:8706881-8723369 | 1.00E-04 | 0.00601 | 0.653 | 48.139 | 75.671 |
| Slc25a42 | chr8:70184339-70212281 | 1.00E-04 | 0.00601 | 0.641 | 20.595 | 32.122 |
| Dusp6 | chr10:99263230-99267489 | 1.00E-04 | 0.00601 | 0.620 | 13.534 | 20.807 |
| Atf4 | chr15:80255183-80257545 | 1.00E-04 | 0.00601 | 0.617 | 78.450 | 120.284 |
| Ptprs | chr17:56412425-56476480 | 1.00E-04 | 0.00601 | -0.667 | 9.747 | 6.137 |
| Il1r2 | chr1:40076211-40125226 | 1.00E-04 | 0.00601 | -0.727 | 31.961 | 19.314 |
| Rdh11 | chr12:79175550-79191819 | 1.00E-04 | 0.00601 | -0.776 | 16.376 | 9.563 |
| Marc1 | chr1:184786766-184811300 | 1.00E-04 | 0.00601 | -0.800 | 25.558 | 14.684 |
| Clec12a | chr6:129350243-129365303 | 1.00E-04 | 0.00601 | -1.097 | 4.812 | 2.250 |
| Col6a6 | chr9:105689416-105828085 | 1.00E-04 | 0.00601 | -1.191 | 1.260 | 0.552 |
| Serpinc1 | chr1:160978605-161003014 | 1.00E-04 | 0.00601 | -1.247 | 4.745 | 1.999 |
| Akr1c6 | chr13:4434342-4457530 | 1.00E-04 | 0.00601 | -1.885 | 4.747 | 1.285 |
| Rgn | chrX:20549817-20562087 | 1.00E-04 | 0.00601 | -2.045 | 4.358 | 1.056 |
| Lrrc8e | chr8:4226826-4237470 | 1.00E-04 | 0.00601 | -2.069 | 0.621 | 0.148 |
| Bdh1 | chr16:31422296-31458901 | 1.00E-04 | 0.00601 | -2.227 | 0.704 | 0.150 |
| Krt17 | chr11:100256216-100260989 | 1.00E-04 | 0.00601 | -7.460 | 23.790 | 0.135 |
| 4833423E24Rik | chr2:85484091-85518935 | 1.00E-04 | 0.00601 | -8.013 | 31.881 | 0.123 |
| Gm9696 | chr3:59952308-59973594 | 1.50E-04 | 0.00844 | 2.139 | 0.353 | 1.555 |
| Nfil3 | chr13:52967208-52981039 | 1.50E-04 | 0.00844 | 0.920 | 3.999 | 7.564 |
| Nop2 | chr6:125131882-125144753 | 1.50E-04 | 0.00844 | 0.655 | 10.349 | 16.298 |
| Hoxa9 | chr6:52223096-52227370 | 1.50E-04 | 0.00844 | 0.635 | 7.071 | 10.983 |
| Cd320 | chr17:33843090-33849774 | 1.50E-04 | 0.00844 | 0.608 | 12.584 | 19.178 |

| | | | | | | |
|---|---|---|---|---|---|---|
| Rcn1 | chr2:105385947-105399319 | 1.50E-04 | 0.00844 | -0.633 | 16.581 | 10.691 |
| Ano1 | chr7:144588548-144738592 | 1.50E-04 | 0.00844 | -0.750 | 5.454 | 3.242 |
| Col16a1 | chr4:130047839-130099277 | 1.50E-04 | 0.00844 | -0.752 | 4.032 | 2.394 |
| Crtac1 | chr19:42283036-42431783 | 1.50E-04 | 0.00844 | -0.802 | 9.010 | 5.168 |
| Pygm | chr19:6384428-6398459 | 1.50E-04 | 0.00844 | -0.928 | 22.541 | 11.845 |
| Apoc3 | chr9:46233049-46235636 | 1.50E-04 | 0.00844 | -1.046 | 40.421 | 19.575 |
| Apoh | chr11:108395296-108414396 | 1.50E-04 | 0.00844 | -1.300 | 9.726 | 3.949 |
| F2 | chr2:91625319-91636457 | 1.50E-04 | 0.00844 | -1.428 | 3.864 | 1.436 |
| Cyp2c29 | chr19:39287084-39330713 | 1.50E-04 | 0.00844 | -1.441 | 4.693 | 1.729 |
| Proc | chr18:32123125-32139570 | 1.50E-04 | 0.00844 | -1.935 | 1.631 | 0.427 |
| Rpl29 | chr9:106429538-106431567 | 2.00E-04 | 0.01043 | 0.704 | 40.011 | 65.191 |
| Clstn3 | chr6:124430755-124464784 | 2.00E-04 | 0.01043 | 0.687 | 26.166 | 42.135 |
| Timm10 | chr2:84827020-84830213 | 2.00E-04 | 0.01043 | 0.667 | 43.979 | 69.841 |
| Coq10b | chr1:55052769-55072702 | 2.00E-04 | 0.01043 | 0.649 | 17.082 | 26.778 |
| Slc20a1 | chr2:129198772-129211612 | 2.00E-04 | 0.01043 | 0.627 | 11.463 | 17.698 |
| Rdh13 | chr7:4425664-4445657 | 2.00E-04 | 0.01043 | 0.606 | 8.865 | 13.489 |
| Abhd15 | chr11:77515116-77520628 | 2.00E-04 | 0.01043 | 0.592 | 20.112 | 30.319 |
| Cd248 | chr19:5068077-5070639 | 2.00E-04 | 0.01043 | -0.590 | 51.630 | 34.309 |
| Tmsb10 | chr6:72957346-72958748 | 2.00E-04 | 0.01043 | -0.601 | 118.821 | 78.346 |
| Clmn | chr12:104763113-104865076 | 2.00E-04 | 0.01043 | -0.625 | 9.249 | 5.996 |
| Eno3 | chr11:70657175-70662513 | 2.00E-04 | 0.01043 | -0.769 | 30.196 | 17.722 |
| Gpr153 | chr4:152274361-152285337 | 2.00E-04 | 0.01043 | -0.799 | 5.958 | 3.423 |
| Ckm | chr7:19411093-19421583 | 2.00E-04 | 0.01043 | -0.994 | 106.701 | 53.572 |
| B3gnt3 | chr8:71691719-71701800 | 2.00E-04 | 0.01043 | -1.136 | 3.988 | 1.814 |
| 1700029J07Rik | chr8:45953605-45975252 | 2.00E-04 | 0.01043 | -1.725 | 1.567 | 0.474 |
| Cdh1 | chr8:106603367-106670246 | 2.00E-04 | 0.01043 | -1.885 | 1.208 | 0.327 |
| Gjb2 | chr14:57098601-57104702 | 2.00E-04 | 0.01043 | -2.127 | 2.031 | 0.465 |
| Slco1a1 | chr6:141907280-141946962 | 2.00E-04 | 0.01043 | -2.178 | 0.490 | 0.108 |

| | | | | | | |
|---|---|---|---|---|---|---|
| Gsg1l | chr7:125878418-126082411 | 2.00E-04 | 0.01043 | -3.248 | 0.569 | 0.060 |
| Lrrc59 | chr11:94629823-94653754 | 2.50E-04 | 0.01228 | 0.626 | 71.027 | 109.610 |
| Lamtor3 | chr3:137918554-137928762 | 2.50E-04 | 0.01228 | 0.613 | 22.211 | 33.960 |
| Tbrg4 | chr11:6615597-6626067 | 2.50E-04 | 0.01228 | 0.607 | 20.782 | 31.644 |
| Mir22hg | chr11:75461538-75466690 | 2.50E-04 | 0.01228 | 0.572 | 33.708 | 50.097 |
| Trabd2b | chr4:114406723-114615098 | 2.50E-04 | 0.01228 | 0.568 | 10.718 | 15.889 |
| Macrod1 | chr19:7056767-7198062 | 2.50E-04 | 0.01228 | -0.603 | 44.199 | 29.095 |
| Fam114a1 | chr5:64970074-65041901 | 2.50E-04 | 0.01228 | -0.606 | 14.865 | 9.766 |
| H2-Q10 | chr17:35470088-35474563 | 2.50E-04 | 0.01228 | -0.684 | 134.147 | 83.496 |
| Pla2g2d | chr4:138775734-138782143 | 2.50E-04 | 0.01228 | -0.687 | 9.511 | 5.907 |
| Xaf1 | chr11:72301628-72313733 | 2.50E-04 | 0.01228 | -0.790 | 8.353 | 4.832 |
| Ptgfr | chr3:151798609-151837528 | 2.50E-04 | 0.01228 | -1.030 | 1.976 | 0.968 |
| Cml2 | chr6:85865421-85869137 | 2.50E-04 | 0.01228 | -1.451 | 1.493 | 0.546 |
| Myo5b | chr18:74442618-74771477 | 2.50E-04 | 0.01228 | -1.536 | 0.960 | 0.331 |
| Rdh7 | chr10:127884026-127888733 | 2.50E-04 | 0.01228 | -1.560 | 3.294 | 1.117 |
| Col17a1 | chr19:47646340-47692042 | 2.50E-04 | 0.01228 | -1.678 | 0.510 | 0.159 |
| Cyp2c44 | chr19:44005021-44029247 | 2.50E-04 | 0.01228 | -1.833 | 1.387 | 0.389 |
| Rnf11 | chr4:109452856-109476505 | 3.00E-04 | 0.01421 | 0.676 | 138.914 | 221.929 |
| Adh1 | chr3:138277644-138290691 | 3.00E-04 | 0.01421 | 0.637 | 135.063 | 210.073 |
| Cachd1 | chr4:100776678-101003748 | 3.00E-04 | 0.01421 | 0.614 | 9.205 | 14.091 |
| Larp1b | chr3:40950630-40977793 | 3.00E-04 | 0.01421 | 0.612 | 23.797 | 36.365 |
| Pcolce | chr5:137605106-137611404 | 3.00E-04 | 0.01421 | -0.565 | 83.563 | 56.471 |
| P2rx5 | chr11:73160529-73172687 | 3.00E-04 | 0.01421 | -0.641 | 14.834 | 9.515 |
| Hal | chr10:93488767-93516743 | 3.00E-04 | 0.01421 | -2.046 | 0.800 | 0.194 |
| Cyp1a2 | chr9:57676936-57683655 | 3.00E-04 | 0.01421 | -2.131 | 2.477 | 0.565 |
| Lad1 | chr1:135818597-135833341 | 3.00E-04 | 0.01421 | -2.974 | 0.494 | 0.063 |
| Krt5 | chr15:101707069-101712891 | 3.00E-04 | 0.01421 | -4.013 | 3.762 | 0.233 |
| Hpdl | chr4:116819906-116821508 | 3.50E-04 | 0.01591 | 1.405 | 1.904 | 5.044 |

| | | | | | | |
|---|---|---|---|---|---|---|
| Alas1 | chr9:106233454-106247954 | 3.50E-04 | 0.01591 | 0.701 | 94.625 | 153.848 |
| Slc35g1 | chr19:38395979-38405607 | 3.50E-04 | 0.01591 | 0.652 | 6.429 | 10.102 |
| Arl4a | chr12:40033290-40037987 | 3.50E-04 | 0.01591 | 0.647 | 66.320 | 103.843 |
| Ubc | chr5:125385964-125390017 | 3.50E-04 | 0.01591 | -0.587 | 76.010 | 50.586 |
| Mturn | chr6:54681623-54703855 | 3.50E-04 | 0.01591 | -0.611 | 20.393 | 13.352 |
| Cd209g | chr8:4134735-4137707 | 3.50E-04 | 0.01591 | -0.658 | 29.195 | 18.497 |
| Gpr137b | chr13:13357619-13393624 | 3.50E-04 | 0.01591 | -0.809 | 5.835 | 3.331 |
| Col1a1 | chr11:94936269-94951856 | 3.50E-04 | 0.01591 | -0.827 | 164.616 | 92.773 |
| Aldob | chr4:49535994-49549483 | 3.50E-04 | 0.01591 | -0.962 | 7.731 | 3.968 |
| Has3 | chr8:106870241-106882902 | 3.50E-04 | 0.01591 | -1.205 | 1.084 | 0.470 |
| Slc23a3 | chr1:75125541-75133890 | 3.50E-04 | 0.01591 | -2.728 | 0.710 | 0.107 |
| Cend1 | chr7:141426450-141429420 | 4.00E-04 | 0.01759 | 1.872 | 0.491 | 1.796 |
| Tmem45b | chr9:31426195-31464238 | 4.00E-04 | 0.01759 | 0.747 | 152.681 | 256.278 |
| Cnr1 | chr4:33924631-33948831 | 4.00E-04 | 0.01759 | 0.676 | 2.835 | 4.529 |
| Cdkn2c | chr4:109660875-109666756 | 4.00E-04 | 0.01759 | 0.671 | 195.956 | 311.965 |
| Cox10 | chr11:63962626-64079472 | 4.00E-04 | 0.01759 | 0.593 | 9.717 | 14.662 |
| Tbc1d16 | chr11:119143042-119228499 | 4.00E-04 | 0.01759 | -0.622 | 3.913 | 2.543 |
| Ttc25 | chr11:100545631-100572566 | 4.00E-04 | 0.01759 | -0.849 | 5.468 | 3.036 |
| Baiap2l2 | chr15:79258194-79285509 | 4.00E-04 | 0.01759 | -1.438 | 2.548 | 0.940 |
| Phyhip | chr14:70457516-70468824 | 4.00E-04 | 0.01759 | -3.746 | 0.696 | 0.052 |
| Dsc2 | chr18:20030797-20059505 | 4.00E-04 | 0.01759 | -5.837 | 1.405 | 0.025 |
| Gpr135 | chr12:72069617-72070991 | 4.50E-04 | 0.01892 | 1.249 | 1.652 | 3.925 |
| Rrp12 | chr19:41862850-41896153 | 4.50E-04 | 0.01892 | 0.705 | 2.734 | 4.456 |
| Hspa8 | chr9:40801272-40805199 | 4.50E-04 | 0.01892 | 0.670 | 127.474 | 202.864 |
| Lmo4 | chr3:144188529-144205255 | 4.50E-04 | 0.01892 | 0.592 | 81.126 | 122.318 |
| Eif4a1 | chr11:69666935-69672423 | 4.50E-04 | 0.01892 | 0.576 | 104.659 | 156.047 |
| Cbx4 | chr11:119077570-119086237 | 4.50E-04 | 0.01892 | 0.570 | 12.166 | 18.059 |
| Alcam | chr16:52248995-52452997 | 4.50E-04 | 0.01892 | 0.566 | 11.051 | 16.365 |

| | | | | | | |
|---|---|---|---|---|---|---|
| Dolk | chr2:30284228-30286354 | 4.50E-04 | 0.01892 | 0.564 | 52.197 | 77.144 |
| Snhg11 | chr2:158375637-158386145 | 4.50E-04 | 0.01892 | -0.565 | 21.101 | 14.268 |
| Tnni2 | chr7:142442467-142444405 | 4.50E-04 | 0.01892 | -0.848 | 78.310 | 43.516 |
| Stard10 | chr7:101321318-101346312 | 4.50E-04 | 0.01892 | -0.901 | 13.253 | 7.096 |
| Mup20 | chr4:62050234-62054117 | 4.50E-04 | 0.01892 | -1.100 | 12.862 | 5.999 |
| Sec14l4 | chr11:4031781-4048009 | 4.50E-04 | 0.01892 | -1.253 | 2.236 | 0.938 |
| Abcb11 | chr2:69238281-69342616 | 4.50E-04 | 0.01892 | -1.942 | 0.708 | 0.184 |
| Rnase2a | chr14:51255261-51256112 | 5.00E-04 | 0.02045 | 1.594 | 2.443 | 7.377 |
| Abhd5 | chr9:122351615-122381523 | 5.00E-04 | 0.02045 | 0.643 | 64.491 | 100.701 |
| Pak1ip1 | chr13:41001009-41013033 | 5.00E-04 | 0.02045 | 0.562 | 35.513 | 52.431 |
| Svep1 | chr4:58042795-58206596 | 5.00E-04 | 0.02045 | -0.565 | 9.754 | 6.593 |
| Sdc1 | chr12:8771395-8793687 | 5.00E-04 | 0.02045 | -0.709 | 7.226 | 4.421 |
| Ift122 | chr6:115853527-115926699 | 5.00E-04 | 0.02045 | -0.839 | 3.468 | 1.938 |
| Myoz1 | chr14:20649101-20656540 | 5.00E-04 | 0.02045 | -0.944 | 12.626 | 6.562 |
| Alb | chr5:90460888-90476603 | 5.00E-04 | 0.02045 | -1.023 | 201.133 | 98.965 |
| Slco1b2 | chr6:141629517-141686635 | 5.00E-04 | 0.02045 | -1.691 | 1.351 | 0.418 |
| 4931406C07Rik | chr9:15283336-15357788 | 5.50E-04 | 0.02189 | 0.771 | 141.182 | 240.865 |
| Ppp1r3b | chr8:35375740-35388137 | 5.50E-04 | 0.02189 | 0.717 | 25.513 | 41.941 |
| Slc4a4 | chr5:88887259-89239656 | 5.50E-04 | 0.02189 | 0.606 | 5.201 | 7.914 |
| Nmd3 | chr3:69722054-69749046 | 5.50E-04 | 0.02189 | 0.570 | 15.168 | 22.524 |
| Mrpl20 | chr4:155803617-155808829 | 5.50E-04 | 0.02189 | 0.565 | 164.588 | 243.562 |
| Dixdc1 | chr9:50662752-50727984 | 5.50E-04 | 0.02189 | -0.637 | 4.986 | 3.205 |
| Adck3 | chr1:179961088-180196020 | 5.50E-04 | 0.02189 | -1.088 | 44.817 | 21.080 |
| Apoa5 | chr9:46268607-46271919 | 5.50E-04 | 0.02189 | -1.297 | 3.311 | 1.348 |
| Afm | chr5:90518948-90553544 | 5.50E-04 | 0.02189 | -2.075 | 0.844 | 0.200 |
| Rps3a1 | chr3:86137939-86142668 | 6.00E-04 | 0.02360 | 0.641 | 73.095 | 113.982 |
| Larp4 | chr15:99970064-100016358 | 6.00E-04 | 0.02360 | 0.582 | 5.344 | 7.999 |
| Lgals3 | chr14:47373859-47386167 | 6.00E-04 | 0.02360 | 0.536 | 57.513 | 83.401 |

| | | | | | | |
|---|---|---|---|---|---|---|
| Haao | chr17:83831353-83846790 | 6.00E-04 | 0.02360 | -1.673 | 2.379 | 0.746 |
| 4632428N05Rik | chr10:60302749-60696490 | 6.50E-04 | 0.02520 | -0.588 | 33.477 | 22.275 |
| Adrb3 | chr8:27225775-27229588 | 6.50E-04 | 0.02520 | -0.710 | 225.016 | 137.581 |
| Stbd1 | chr5:92603050-92606579 | 6.50E-04 | 0.02520 | -0.899 | 5.684 | 3.048 |
| Cfi | chr3:129836738-129875328 | 6.50E-04 | 0.02520 | -1.837 | 1.555 | 0.435 |
| Nlrp10 | chr7:108921852-108930158 | 6.50E-04 | 0.02520 | -1.950 | 0.489 | 0.127 |
| A530050N04Rik | chr18:61470224-61484607 | 7.00E-04 | 0.02668 | 3.744 | 0.115 | 1.539 |
| Klhl25 | chr7:75848337-75874130 | 7.00E-04 | 0.02668 | 0.671 | 4.580 | 7.291 |
| Dpyd | chr3:118562177-119432918 | 7.00E-04 | 0.02668 | -0.581 | 12.123 | 8.105 |
| Col6a3 | chr1:90766859-90843971 | 7.00E-04 | 0.02668 | -0.651 | 44.558 | 28.384 |
| Pgam2 | chr11:5801636-5803796 | 7.00E-04 | 0.02668 | -1.092 | 11.079 | 5.196 |
| Ces1c | chr8:93099015-93131283 | 7.00E-04 | 0.02668 | -1.242 | 4.918 | 2.080 |
| Cpeb2 | chr5:43151685-43289724 | 7.50E-04 | 0.02787 | 0.562 | 5.496 | 8.116 |
| Tomm70a | chr16:57121713-57154530 | 7.50E-04 | 0.02787 | 0.542 | 27.437 | 39.957 |
| Irf4 | chr13:30749257-30766927 | 7.50E-04 | 0.02787 | -0.632 | 5.065 | 3.270 |
| Cmya5 | chr13:93040714-93144724 | 7.50E-04 | 0.02787 | -0.834 | 1.169 | 0.655 |
| Itih4 | chr14:30886475-30901986 | 7.50E-04 | 0.02787 | -1.004 | 3.175 | 1.583 |
| Cyp2d9 | chr15:82452376-82456827 | 7.50E-04 | 0.02787 | -1.349 | 3.820 | 1.500 |
| Sult2a8 | chr7:14410685-14446587 | 7.50E-04 | 0.02787 | -1.535 | 1.791 | 0.618 |
| Esrp1 | chr4:11331932-11386783 | 7.50E-04 | 0.02787 | -2.091 | 0.560 | 0.131 |
| Pdzk1ip1 | chr4:115088707-115093894 | 7.50E-04 | 0.02787 | -4.778 | 8.182 | 0.298 |
| Cd164l2 | chr4:133220808-133224554 | 8.00E-04 | 0.02933 | 1.338 | 2.570 | 6.498 |
| Nolc1 | chr19:46075846-46085543 | 8.00E-04 | 0.02933 | 0.542 | 10.549 | 15.356 |
| Ryr1 | chr7:29003339-29125151 | 8.00E-04 | 0.02933 | -0.800 | 1.738 | 0.999 |
| Acox2 | chr14:8225510-8259019 | 8.00E-04 | 0.02933 | -1.575 | 1.726 | 0.579 |
| Bbox1 | chr2:110265082-110305725 | 8.00E-04 | 0.02933 | -2.836 | 0.844 | 0.118 |
| Ccl8 | chr11:82115184-82116799 | 8.50E-04 | 0.03041 | 0.664 | 62.243 | 98.643 |
| Megf9 | chr4:70431926-70534928 | 8.50E-04 | 0.03041 | 0.565 | 8.841 | 13.083 |

| | | | | | | |
|---|---|---|---|---|---|---|
| Asns | chr6:7675170-7693182 | 8.50E-04 | 0.03041 | 0.539 | 49.808 | 72.370 |
| Ftsj3 | chr11:106249143-106255802 | 8.50E-04 | 0.03041 | 0.526 | 14.891 | 21.445 |
| Atrn | chr2:130906495-131030326 | 8.50E-04 | 0.03041 | 0.518 | 9.852 | 14.105 |
| Slco2b1 | chr7:99657803-99711340 | 8.50E-04 | 0.03041 | -0.557 | 27.839 | 18.928 |
| Wscd2 | chr5:113550419-113589725 | 8.50E-04 | 0.03041 | -0.618 | 6.627 | 4.319 |
| Atp1b2 | chr11:69599749-69605960 | 8.50E-04 | 0.03041 | -0.649 | 7.914 | 5.047 |
| Alpk1 | chr3:127670309-127780527 | 8.50E-04 | 0.03041 | -0.872 | 2.644 | 1.445 |
| Map3k6 | chr4:133240817-133252928 | 9.00E-04 | 0.03178 | 0.553 | 7.081 | 10.391 |
| Tspan18 | chr2:93201759-93334487 | 9.00E-04 | 0.03178 | -0.565 | 33.218 | 22.459 |
| Tlr5 | chr1:182954787-182976044 | 9.00E-04 | 0.03178 | -0.688 | 5.846 | 3.629 |
| Serpina1c | chr12:103894925-103904950 | 9.00E-04 | 0.03178 | -0.859 | 9.743 | 5.372 |
| Cep83os | chr10:94673492-94688613 | 9.00E-04 | 0.03178 | -0.966 | 2.412 | 1.235 |
| Trmt10c | chr16:56033719-56037774 | 9.50E-04 | 0.03269 | 0.561 | 13.281 | 19.598 |
| Cycs | chr6:50562562-50566474 | 9.50E-04 | 0.03269 | 0.554 | 24.593 | 36.117 |
| Fam195a | chr17:25863697-25868738 | 9.50E-04 | 0.03269 | 0.541 | 102.218 | 148.721 |
| Tomm40 | chr7:19701312-19715429 | 9.50E-04 | 0.03269 | 0.505 | 52.925 | 75.112 |
| Mgl2 | chr11:70130356-70137542 | 9.50E-04 | 0.03269 | -0.522 | 52.160 | 36.319 |
| Gpc3 | chrX:52272426-52619047 | 9.50E-04 | 0.03269 | -0.586 | 13.847 | 9.227 |
| Srpx | chrX:10037976-10117661 | 9.50E-04 | 0.03269 | -0.615 | 10.407 | 6.795 |
| Comp | chr8:70373547-70382066 | 9.50E-04 | 0.03269 | -1.345 | 1.724 | 0.679 |
| Serpinf2 | chr11:75431735-75439501 | 9.50E-04 | 0.03269 | -1.441 | 1.778 | 0.655 |
| Sorcs2 | chr5:36017180-36398139 | 9.50E-04 | 0.03269 | -2.587 | 6.663 | 1.109 |
| Rprml | chr11:103649508-103650580 | 1.00E-03 | 0.03423 | -0.962 | 9.570 | 4.912 |
| Sec14l2 | chr11:4097039-4118729 | 1.00E-03 | 0.03423 | -1.356 | 1.773 | 0.693 |
| Nol10 | chr12:17348492-17430095 | 1.05E-03 | 0.03514 | 0.661 | 4.023 | 6.362 |
| Sfrp2 | chr3:83766320-83774314 | 1.05E-03 | 0.03514 | -0.513 | 39.938 | 27.988 |
| Dhrs3 | chr4:144892826-144927645 | 1.05E-03 | 0.03514 | -0.534 | 105.898 | 73.117 |
| Tmem43 | chr6:91473750-91488458 | 1.05E-03 | 0.03514 | -0.571 | 116.750 | 78.572 |

| | | | | | | |
|---|---|---|---|---|---|---|
| 2810474O19Rik | chr6:149309413-149335663 | 1.05E-03 | 0.03514 | -0.628 | 4.145 | 2.682 |
| Lrrc17 | chr5:21483846-21645605 | 1.05E-03 | 0.03514 | -0.638 | 10.544 | 6.776 |
| Isoc2b | chr7:4844959-4866179 | 1.05E-03 | 0.03514 | -0.691 | 20.413 | 12.648 |
| Maff | chr15:79346620-79359076 | 1.05E-03 | 0.03514 | -0.962 | 5.034 | 2.585 |
| Ces3b | chr8:105083754-105093591 | 1.05E-03 | 0.03514 | -1.835 | 0.925 | 0.259 |
| Pla2g2e | chr4:138877941-138882814 | 1.10E-03 | 0.03654 | 1.016 | 5.933 | 11.997 |
| Ppp1r10 | chr17:35917195-35932283 | 1.10E-03 | 0.03654 | 0.551 | 6.408 | 9.391 |
| Dnajb9 | chr12:44205896-44210068 | 1.10E-03 | 0.03654 | 0.530 | 28.759 | 41.531 |
| Apln | chrX:48025145-48034852 | 1.15E-03 | 0.03783 | 0.881 | 1.700 | 3.131 |
| Cadm3 | chr1:173334253-173367695 | 1.15E-03 | 0.03783 | -0.529 | 12.516 | 8.676 |
| Sgms2 | chr3:131318985-131344923 | 1.15E-03 | 0.03783 | -0.617 | 3.508 | 2.287 |
| Camk2b | chr11:5969665-6065748 | 1.15E-03 | 0.03783 | -1.209 | 1.143 | 0.494 |
| Cdkn2b | chr4:89306288-89311032 | 1.20E-03 | 0.03881 | 0.689 | 8.184 | 13.193 |
| Abcd3 | chr3:121758909-121815215 | 1.20E-03 | 0.03881 | 0.599 | 53.106 | 80.427 |
| Fam83a | chr15:57985902-58010702 | 1.20E-03 | 0.03881 | 0.583 | 9.545 | 14.296 |
| Mkks | chr2:136873780-137069778 | 1.20E-03 | 0.03881 | 0.545 | 13.712 | 20.009 |
| Rhot1 | chr11:80209054-80267907 | 1.20E-03 | 0.03881 | 0.497 | 16.783 | 23.681 |
| Bgn | chrX:73483634-73495936 | 1.20E-03 | 0.03881 | -0.535 | 81.050 | 55.954 |
| Sdr42e1 | chr8:117661398-117671515 | 1.20E-03 | 0.03881 | -0.657 | 7.715 | 4.892 |
| Usp2 | chr9:44067020-44095627 | 1.25E-03 | 0.04014 | 0.627 | 4.661 | 7.201 |
| Cyp27a1 | chr1:74713573-74737890 | 1.25E-03 | 0.04014 | -0.501 | 32.654 | 23.081 |
| Ckmt1 | chr2:121358640-121380940 | 1.25E-03 | 0.04014 | -4.378 | 2.500 | 0.120 |
| Tmem11 | chr11:60864451-60879038 | 1.30E-03 | 0.04145 | 0.530 | 41.160 | 59.419 |
| Tbc1d2 | chr4:46604389-46650199 | 1.30E-03 | 0.04145 | -0.588 | 5.837 | 3.882 |
| Clca3a2 | chr3:144796558-144819494 | 1.30E-03 | 0.04145 | -2.897 | 0.556 | 0.075 |
| Rxrg | chr1:167598361-167639623 | 1.35E-03 | 0.04244 | 0.539 | 19.539 | 28.395 |
| Snx10 | chr6:51523902-51590670 | 1.35E-03 | 0.04244 | 0.509 | 48.016 | 68.311 |
| Chrdl1 | chrX:143285673-143394262 | 1.35E-03 | 0.04244 | -0.551 | 20.075 | 13.701 |

| | | | | | | |
|---|---|---|---|---|---|---|
| Ugt2b34 | chr5:86889769-86906937 | 1.35E-03 | 0.04244 | -1.725 | 0.706 | 0.214 |
| Hsd17b13 | chr5:103955441-103977388 | 1.35E-03 | 0.04244 | -1.821 | 1.285 | 0.364 |
| Cdkl4 | chr17:80523549-80563834 | 1.35E-03 | 0.04244 | -1.929 | 0.469 | 0.123 |
| Idi1 | chr13:8885605-8892396 | 1.40E-03 | 0.04380 | -0.787 | 4.450 | 2.579 |
| Hfe2 | chr3:96525184-96529216 | 1.40E-03 | 0.04380 | -1.275 | 2.843 | 1.175 |
| Col6a2 | chr10:76595755-76623404 | 1.45E-03 | 0.04516 | -0.629 | 111.648 | 72.186 |
| Adprhl1 | chr8:13235661-13254162 | 1.45E-03 | 0.04516 | -1.715 | 3.200 | 0.975 |
| Rnmtl1 | chr11:76243735-76250622 | 1.50E-03 | 0.04629 | 0.637 | 9.289 | 14.443 |
| Mfng | chr15:78755882-78773445 | 1.50E-03 | 0.04629 | -0.512 | 103.827 | 72.829 |
| Spint2 | chr7:29256329-29281977 | 1.50E-03 | 0.04629 | -0.825 | 9.990 | 5.641 |
| Mboat2 | chr12:24831598-24960299 | 1.50E-03 | 0.04629 | -1.452 | 1.046 | 0.382 |
| Aldh1a1 | chr19:20601981-20643462 | 1.55E-03 | 0.04750 | 0.520 | 41.199 | 59.075 |
| Tsc22d3 | chrX:140539528-140600522 | 1.55E-03 | 0.04750 | 0.509 | 36.398 | 51.796 |
| Anxa8 | chr14:34051129-34102754 | 1.55E-03 | 0.04750 | -0.686 | 10.499 | 6.528 |
| Retn | chr8:3655769-3659818 | 1.60E-03 | 0.04816 | 1.042 | 2262.020 | 4659.250 |
| Hsd17b7 | chr1:169949536-169969205 | 1.60E-03 | 0.04816 | 0.590 | 3.789 | 5.703 |
| Ptgfrn | chr3:101040235-101110166 | 1.60E-03 | 0.04816 | -0.519 | 9.991 | 6.973 |
| Sema3b | chr9:107597673-107609241 | 1.60E-03 | 0.04816 | -0.520 | 13.385 | 9.334 |
| Ptafr | chr4:132564066-132580866 | 1.60E-03 | 0.04816 | -0.700 | 8.903 | 5.479 |
| Hrg | chr16:22951071-22961659 | 1.60E-03 | 0.04816 | -1.446 | 2.108 | 0.774 |
| Sbk2 | chr7:4957080-4964348 | 1.60E-03 | 0.04816 | -1.744 | 1.887 | 0.563 |
| Klk13 | chr7:43712566-43726758 | 1.60E-03 | 0.04816 | -3.407 | 1.432 | 0.135 |
| Ube3a | chr7:59228749-59306727 | 1.65E-03 | 0.04933 | 0.535 | 5.501 | 7.970 |
| Efemp2 | chr19:5474689-5481854 | 1.65E-03 | 0.04933 | -0.563 | 19.538 | 13.224 |
| Cfhr2 | chr1:139810291-139858699 | 1.65E-03 | 0.04933 | -1.670 | 1.312 | 0.412 |

**Table S4**: GSEA analysis of browning related gene sets in sWAT of WT and Rbp$^{-/-}$ mice gene sets, including Hallmark, Reactome and GO pathways.

**(i)** **HALLMARK_OXIDATIVE_PHOSPHORYLATION** (Size: 192; ES: 0.60; NES: 2.43; NOM p-value: 0.000; FDR q-value: 0.000)

| Probe | Rank in Gene List | Rank Metric Score | Running ES | Core Enrichment |
|---|---|---|---|---|
| ATP1B1 | 792 | 0.77297 | -0.03057 | Yes |
| BCKDHA | 814 | 0.76055 | -0.02056 | Yes |
| LDHB | 885 | 0.73395 | -0.01354 | Yes |
| PDHB | 930 | 0.72104 | -0.00533 | Yes |
| ECI1 | 939 | 0.71859 | 0.00475 | Yes |
| NDUFA8 | 1173 | 0.65806 | 0.00205 | Yes |
| GPX4 | 1311 | 0.62479 | 0.00394 | Yes |
| COX6A1 | 1326 | 0.62148 | 0.01228 | Yes |
| NDUFS7 | 1337 | 0.61927 | 0.02080 | Yes |
| SDHB | 1398 | 0.60668 | 0.02649 | Yes |
| GOT2 | 1421 | 0.60064 | 0.03411 | Yes |
| ACO2 | 1425 | 0.60040 | 0.04272 | Yes |
| AFG3L2 | 1441 | 0.59757 | 0.05066 | Yes |
| NDUFA3 | 1445 | 0.59686 | 0.05922 | Yes |
| NDUFV2 | 1466 | 0.59397 | 0.06684 | Yes |
| NDUFB8 | 1470 | 0.59359 | 0.07536 | Yes |
| UQCRH | 1530 | 0.58188 | 0.08074 | Yes |
| UQCRQ | 1542 | 0.57977 | 0.08863 | Yes |
| CYC1 | 1555 | 0.57708 | 0.09643 | Yes |
| UQCR10 | 1557 | 0.57677 | 0.10481 | Yes |
| ATP5H | 1621 | 0.56505 | 0.10973 | Yes |
| MRPL34 | 1636 | 0.56131 | 0.11719 | Yes |
| NDUFA6 | 1657 | 0.55869 | 0.12430 | Yes |
| IDH3B | 1667 | 0.55643 | 0.13196 | Yes |
| NDUFV1 | 1705 | 0.55164 | 0.13806 | Yes |
| ETFB | 1728 | 0.54857 | 0.14491 | Yes |
| SDHD | 1735 | 0.54724 | 0.15259 | Yes |
| COX5B | 1757 | 0.54331 | 0.15942 | Yes |
| UQCRC2 | 1760 | 0.54307 | 0.16725 | Yes |
| PDK4 | 1762 | 0.54301 | 0.17513 | Yes |
| NDUFS8 | 1790 | 0.53833 | 0.18157 | Yes |
| NDUFS3 | 1791 | 0.53820 | 0.18943 | Yes |
| ATP5J2 | 1822 | 0.53297 | 0.19564 | Yes |
| COX5A | 1832 | 0.53190 | 0.20293 | Yes |
| UQCR11 | 1835 | 0.53169 | 0.21060 | Yes |
| SLC25A20 | 1902 | 0.52215 | 0.21474 | Yes |
| GRPEL1 | 1919 | 0.51966 | 0.22149 | Yes |
| COX4I1 | 1922 | 0.51948 | 0.22897 | Yes |
| CS | 1962 | 0.51432 | 0.23442 | Yes |
| PHYH | 2002 | 0.50711 | 0.23977 | Yes |
| DECR1 | 2007 | 0.50687 | 0.24697 | Yes |
| NDUFA4 | 2008 | 0.50684 | 0.25437 | Yes |
| IDH3G | 2018 | 0.50537 | 0.26128 | Yes |
| UQCRC1 | 2043 | 0.50269 | 0.26736 | Yes |
| SDHC | 2054 | 0.50142 | 0.27416 | Yes |
| NDUFA7 | 2061 | 0.50077 | 0.28116 | Yes |
| ATP5D | 2074 | 0.49957 | 0.28782 | Yes |
| COX6B1 | 2080 | 0.49895 | 0.29485 | Yes |
| SDHA | 2081 | 0.49873 | 0.30214 | Yes |
| ATP5A1 | 2089 | 0.49804 | 0.30904 | Yes |
| PDHX | 2107 | 0.49467 | 0.31537 | Yes |
| HADHA | 2138 | 0.49148 | 0.32097 | Yes |
| NDUFAB1 | 2142 | 0.49114 | 0.32799 | Yes |

| | | | | |
|---|---|---|---|---|
| UQCRFS1 | 2146 | 0.49090 | 0.33500 | Yes |
| ATP5O | 2154 | 0.48965 | 0.34179 | Yes |
| PHB2 | 2212 | 0.48141 | 0.34581 | Yes |
| DLST | 2224 | 0.47969 | 0.35224 | Yes |
| NDUFB2 | 2225 | 0.47950 | 0.35924 | Yes |
| ISCU | 2227 | 0.47921 | 0.36619 | Yes |
| NDUFC1 | 2238 | 0.47774 | 0.37264 | Yes |
| NDUFS2 | 2239 | 0.47765 | 0.37962 | Yes |
| MRPL15 | 2266 | 0.47517 | 0.38519 | Yes |
| RETSAT | 2283 | 0.47356 | 0.39127 | Yes |
| ISCA1 | 2312 | 0.47022 | 0.39666 | Yes |
| NDUFS6 | 2317 | 0.46973 | 0.40331 | Yes |
| HADHB | 2326 | 0.46847 | 0.40973 | Yes |
| OGDH | 2334 | 0.46756 | 0.41619 | Yes |
| COX7A2L | 2437 | 0.45540 | 0.41746 | Yes |
| SLC25A11 | 2527 | 0.44714 | 0.41929 | Yes |
| NDUFA9 | 2541 | 0.44611 | 0.42512 | Yes |
| ETFDH | 2544 | 0.44590 | 0.43153 | Yes |
| COX10 | 2549 | 0.44535 | 0.43782 | Yes |
| NDUFC2 | 2555 | 0.44514 | 0.44406 | Yes |
| NDUFB6 | 2559 | 0.44459 | 0.45040 | Yes |
| COX7B | 2718 | 0.42880 | 0.44831 | Yes |
| MDH2 | 2722 | 0.42871 | 0.45442 | Yes |
| PDHA1 | 2727 | 0.42774 | 0.46046 | Yes |
| CYB5R3 | 2733 | 0.42726 | 0.46644 | Yes |
| ACADVL | 2763 | 0.42470 | 0.47111 | Yes |
| TIMM8B | 2776 | 0.42298 | 0.47666 | Yes |
| SUCLG1 | 2786 | 0.42214 | 0.48235 | Yes |
| ATP5B | 2873 | 0.41343 | 0.48385 | Yes |
| FXN | 2889 | 0.41209 | 0.48907 | Yes |
| DLAT | 2908 | 0.41111 | 0.49413 | Yes |
| OXA1L | 2934 | 0.40914 | 0.49879 | Yes |
| ECH1 | 2937 | 0.40902 | 0.50466 | Yes |
| TIMM13 | 2956 | 0.40713 | 0.50965 | Yes |
| ATP5F1 | 2977 | 0.40538 | 0.51452 | Yes |
| COX8A | 3015 | 0.40149 | 0.51843 | Yes |
| PRDX3 | 3022 | 0.40100 | 0.52397 | Yes |
| NDUFB5 | 3030 | 0.40032 | 0.52945 | Yes |
| RHOT2 | 3100 | 0.39475 | 0.53157 | Yes |
| COX7C | 3164 | 0.38996 | 0.53394 | Yes |
| MRPS12 | 3176 | 0.38879 | 0.53904 | Yes |
| COX7A2 | 3241 | 0.38343 | 0.54126 | Yes |
| ATP5C1 | 3298 | 0.37814 | 0.54383 | Yes |
| MRPS15 | 3307 | 0.37715 | 0.54892 | Yes |
| TOMM22 | 3355 | 0.37360 | 0.55189 | Yes |
| HSPA9 | 3374 | 0.37216 | 0.55638 | Yes |
| COX6C | 3385 | 0.37129 | 0.56127 | Yes |
| CYCS | 3399 | 0.37034 | 0.56600 | Yes |
| ACAT1 | 3424 | 0.36748 | 0.57010 | Yes |
| ATP5G2 | 3437 | 0.36644 | 0.57482 | Yes |
| ACAA2 | 3649 | 0.34925 | 0.56877 | Yes |
| NDUFA5 | 3672 | 0.34736 | 0.57268 | Yes |
| COX17 | 3713 | 0.34397 | 0.57560 | Yes |
| NDUFB4 | 3721 | 0.34356 | 0.58025 | Yes |
| NDUFA1 | 3797 | 0.33689 | 0.58120 | Yes |
| VDAC1 | 3837 | 0.33307 | 0.58401 | Yes |
| HSD17B10 | 3922 | 0.32537 | 0.58432 | Yes |
| MFN2 | 3972 | 0.32216 | 0.58644 | Yes |
| NDUFB7 | 3979 | 0.32162 | 0.59082 | Yes |
| MDH1 | 4004 | 0.31968 | 0.59423 | Yes |
| NDUFS1 | 4014 | 0.31893 | 0.59841 | Yes |
| IDH2 | 4039 | 0.31755 | 0.60178 | Yes |

| | | | | |
|---|---|---|---|---|
| IMMT | 4195 | 0.30606 | 0.59806 | No |
| NDUFS4 | 4268 | 0.30075 | 0.59865 | No |
| POLR2F | 4409 | 0.29103 | 0.59550 | No |
| TIMM17A | 4518 | 0.28332 | 0.59394 | No |
| LDHA | 4594 | 0.27905 | 0.59405 | No |
| ECHS1 | 4609 | 0.27808 | 0.59737 | No |
| MRPL11 | 4667 | 0.27430 | 0.59837 | No |
| ATP6V0C | 4837 | 0.26244 | 0.59327 | No |
| ATP6V0B | 4860 | 0.26111 | 0.59592 | No |
| UQCRB | 4950 | 0.25566 | 0.59495 | No |
| ACADM | 4974 | 0.25496 | 0.59746 | No |
| PMPCA | 5044 | 0.25030 | 0.59747 | No |
| TIMM10 | 5314 | 0.23275 | 0.58666 | No |
| IDH3A | 5350 | 0.23049 | 0.58818 | No |
| OPA1 | 5362 | 0.22955 | 0.59095 | No |
| SUPV3L1 | 5491 | 0.22297 | 0.58744 | No |
| ETFA | 5494 | 0.22288 | 0.59059 | No |
| MRPL35 | 5514 | 0.22225 | 0.59284 | No |
| MTX2 | 5554 | 0.22015 | 0.59399 | No |
| SLC25A5 | 5652 | 0.21508 | 0.59201 | No |
| TIMM50 | 5662 | 0.21451 | 0.59467 | No |
| ATP6V1D | 5687 | 0.21300 | 0.59651 | No |
| ATP5E | 5730 | 0.21029 | 0.59736 | No |
| COX15 | 5754 | 0.20913 | 0.59920 | No |
| NDUFA2 | 5828 | 0.20544 | 0.59835 | No |
| AIFM1 | 5976 | 0.19683 | 0.59345 | No |
| MRPS22 | 6098 | 0.18973 | 0.58983 | No |
| VDAC2 | 6108 | 0.18908 | 0.59212 | No |
| FDX1 | 6340 | 0.17627 | 0.58248 | No |
| SURF1 | 6487 | 0.16745 | 0.57721 | No |
| DLD | 6535 | 0.16489 | 0.57714 | No |
| COX11 | 6556 | 0.16379 | 0.57847 | No |
| MAOB | 6680 | 0.15683 | 0.57426 | No |
| MGST3 | 6740 | 0.15346 | 0.57339 | No |
| MRPS30 | 6932 | 0.14437 | 0.56540 | No |
| ATP5G1 | 6963 | 0.14294 | 0.56591 | No |
| LRPPRC | 6986 | 0.14215 | 0.56682 | No |
| BDH2 | 7123 | 0.13572 | 0.56162 | No |
| ATP6V1E1 | 7233 | 0.13066 | 0.55776 | No |
| MRPS11 | 7283 | 0.12806 | 0.55704 | No |
| SUCLA2 | 7299 | 0.12738 | 0.55811 | No |
| ALDH6A1 | 7336 | 0.12579 | 0.55805 | No |
| SLC25A3 | 7411 | 0.12190 | 0.55592 | No |
| SLC25A4 | 7976 | 0.09535 | 0.52750 | No |
| VDAC3 | 8258 | 0.08313 | 0.51386 | No |
| CASP7 | 8355 | 0.07845 | 0.50994 | No |
| NDUFB3 | 8701 | 0.06244 | 0.49261 | No |
| BAX | 8817 | 0.05656 | 0.48736 | No |
| ATP6V1F | 9140 | 0.04285 | 0.47097 | No |
| ATP6AP1 | 9255 | 0.03747 | 0.46549 | No |
| ATP6V1G1 | 9500 | 0.02716 | 0.45299 | No |
| ATP6V1H | 9508 | 0.02695 | 0.45302 | No |
| HTRA2 | 9542 | 0.02568 | 0.45165 | No |
| OAT | 9799 | 0.01496 | 0.43834 | No |
| HCCS | 9848 | 0.01248 | 0.43598 | No |
| ATP6V1C1 | 10098 | 0.00200 | 0.42285 | No |
| TCIRG1 | 10135 | 0.00049 | 0.42095 | No |
| SLC25A12 | 10334 | -0.00928 | 0.41062 | No |
| ATP5J | 10339 | -0.00943 | 0.41055 | No |
| ATP5G3 | 11084 | -0.04418 | 0.37187 | No |
| NNT | 11360 | -0.05772 | 0.35818 | No |
| TIMM9 | 11640 | -0.07029 | 0.34446 | No |

| | | | | |
|---|---|---|---|---|
| ABCB7 | 11828 | -0.07813 | 0.33572 | No |
| NQO2 | 12076 | -0.08965 | 0.32397 | No |
| ACADSB | 12130 | -0.09293 | 0.32253 | No |
| IDH1 | 12233 | -0.09786 | 0.31857 | No |
| TOMM70A | 12695 | -0.12086 | 0.29597 | No |
| RHOT1 | 12854 | -0.12848 | 0.28950 | No |
| MTRF1 | 13362 | -0.15361 | 0.26494 | No |
| ALAS1 | 13640 | -0.16709 | 0.25275 | No |
| POR | 14095 | -0.19372 | 0.23158 | No |
| GLUD1 | 14239 | -0.20186 | 0.22697 | No |
| CYB5A | 14491 | -0.21789 | 0.21689 | No |
| MTRR | 16079 | -0.33287 | 0.13787 | No |
| CPT1A | 16211 | -0.34491 | 0.13599 | No |
| PDP1 | 17559 | -0.50610 | 0.07219 | No |
| ATP5L | 18267 | -0.66981 | 0.04461 | No |

**(ii) REACTOME_TCA_CYCLE_AND_RESPIRATORY_ELECTRON_TRANSPORT**
(Size: 112; ES: 0.67; NES: 2.53; NOM p-value: 0.000; FDR q-value: 0.000)

| Probe | Rank In Gene List | Rank Metric Score | Running Es | Core Enrichment |
|---|---|---|---|---|
| UCP3 | 326 | 1.14052 | 0.00552 | Yes |
| NDUFA12 | 338 | 1.11885 | 0.02719 | Yes |
| PDK1 | 738 | 0.80232 | 0.02214 | Yes |
| LDHB | 885 | 0.73395 | 0.02905 | Yes |
| PDHB | 930 | 0.72104 | 0.04107 | Yes |
| NDUFB9 | 962 | 0.71030 | 0.05356 | Yes |
| NDUFA10 | 1082 | 0.67688 | 0.06076 | Yes |
| PDK2 | 1153 | 0.66090 | 0.07021 | Yes |
| NDUFA8 | 1173 | 0.65806 | 0.08230 | Yes |
| SLC16A1 | 1241 | 0.64120 | 0.09152 | Yes |
| COX6A1 | 1326 | 0.62148 | 0.09946 | Yes |
| NDUFS7 | 1337 | 0.61927 | 0.11124 | Yes |
| SDHB | 1398 | 0.60668 | 0.12015 | Yes |
| ACO2 | 1425 | 0.60040 | 0.13072 | Yes |
| NDUFA3 | 1445 | 0.59686 | 0.14158 | Yes |
| NDUFV2 | 1466 | 0.59397 | 0.15234 | Yes |
| NDUFB8 | 1470 | 0.59359 | 0.16399 | Yes |
| UQCRH | 1530 | 0.58188 | 0.17245 | Yes |
| UQCRQ | 1542 | 0.57977 | 0.18340 | Yes |
| CYC1 | 1555 | 0.57708 | 0.19424 | Yes |
| ATP5H | 1621 | 0.56505 | 0.20205 | Yes |
| NDUFB10 | 1633 | 0.56200 | 0.21265 | Yes |
| NDUFA6 | 1657 | 0.55869 | 0.22255 | Yes |
| IDH3B | 1667 | 0.55643 | 0.23314 | Yes |
| NDUFV1 | 1705 | 0.55164 | 0.24216 | Yes |
| ETFB | 1728 | 0.54857 | 0.25191 | Yes |
| SDHD | 1735 | 0.54724 | 0.26247 | Yes |
| COX5B | 1757 | 0.54331 | 0.27217 | Yes |
| UQCRC2 | 1760 | 0.54307 | 0.28286 | Yes |
| PDK4 | 1762 | 0.54301 | 0.29361 | Yes |
| NDUFS8 | 1790 | 0.53833 | 0.30289 | Yes |
| NDUFS3 | 1791 | 0.53820 | 0.31359 | Yes |
| ATP5J2 | 1822 | 0.53297 | 0.32261 | Yes |
| COX5A | 1832 | 0.53190 | 0.33271 | Yes |
| UQCR11 | 1835 | 0.53169 | 0.34318 | Yes |
| COX4I1 | 1922 | 0.51948 | 0.34898 | Yes |
| CS | 1962 | 0.51432 | 0.35715 | Yes |
| NDUFA4 | 2008 | 0.50684 | 0.36486 | Yes |
| IDH3G | 2018 | 0.50537 | 0.37444 | Yes |
| UQCRC1 | 2043 | 0.50269 | 0.38317 | Yes |

| | | | | |
|---|---|---|---|---|
| SDHC | 2054 | 0.50142 | 0.39261 | Yes |
| NDUFA7 | 2061 | 0.50077 | 0.40225 | Yes |
| ATP5D | 2074 | 0.49957 | 0.41156 | Yes |
| COX6B1 | 2080 | 0.49895 | 0.42121 | Yes |
| SDHA | 2081 | 0.49873 | 0.43113 | Yes |
| ATP5A1 | 2089 | 0.49804 | 0.44066 | Yes |
| PDHX | 2107 | 0.49467 | 0.44961 | Yes |
| NDUFAB1 | 2142 | 0.49114 | 0.45758 | Yes |
| UQCRFS1 | 2146 | 0.49090 | 0.46718 | Yes |
| ATP5O | 2154 | 0.48965 | 0.47655 | Yes |
| NDUFA11 | 2189 | 0.48486 | 0.48440 | Yes |
| DLST | 2224 | 0.47969 | 0.49215 | Yes |
| NDUFB2 | 2225 | 0.47950 | 0.50168 | Yes |
| NDUFC1 | 2238 | 0.47774 | 0.51055 | Yes |
| NDUFS2 | 2239 | 0.47765 | 0.52005 | Yes |
| NDUFA13 | 2287 | 0.47305 | 0.52698 | Yes |
| NDUFS6 | 2317 | 0.46973 | 0.53479 | Yes |
| OGDH | 2334 | 0.46756 | 0.54325 | Yes |
| NDUFV3 | 2401 | 0.46021 | 0.54893 | Yes |
| COX7A2L | 2437 | 0.45540 | 0.55614 | Yes |
| NDUFA9 | 2541 | 0.44611 | 0.55959 | Yes |
| ETFDH | 2544 | 0.44590 | 0.56835 | Yes |
| NDUFC2 | 2555 | 0.44514 | 0.57667 | Yes |
| NDUFB6 | 2559 | 0.44459 | 0.58535 | Yes |
| COX7B | 2718 | 0.42880 | 0.58556 | Yes |
| MDH2 | 2722 | 0.42871 | 0.59393 | Yes |
| PDHA1 | 2727 | 0.42774 | 0.60222 | Yes |
| BSG | 2741 | 0.42682 | 0.61003 | Yes |
| SUCLG1 | 2786 | 0.42214 | 0.61610 | Yes |
| ATP5B | 2873 | 0.41343 | 0.61980 | Yes |
| DLAT | 2908 | 0.41111 | 0.62618 | Yes |
| ATP5F1 | 2977 | 0.40538 | 0.63066 | Yes |
| COX8A | 3015 | 0.40149 | 0.63670 | Yes |
| NDUFB5 | 3030 | 0.40032 | 0.64392 | Yes |
| COX7C | 3164 | 0.38996 | 0.64468 | Yes |
| ATP5C1 | 3298 | 0.37814 | 0.64519 | Yes |
| COX6C | 3385 | 0.37129 | 0.64805 | Yes |
| CYCS | 3399 | 0.37034 | 0.65473 | Yes |
| NDUFA5 | 3672 | 0.34736 | 0.64732 | Yes |
| NDUFB4 | 3721 | 0.34356 | 0.65163 | Yes |
| NDUFA1 | 3797 | 0.33689 | 0.65438 | Yes |
| PDP2 | 3802 | 0.33627 | 0.66085 | Yes |
| SLC16A3 | 3894 | 0.32836 | 0.66259 | Yes |
| NDUFB7 | 3979 | 0.32162 | 0.66457 | Yes |
| NDUFS1 | 4014 | 0.31893 | 0.66912 | Yes |
| IDH2 | 4039 | 0.31755 | 0.67417 | Yes |
| NDUFS4 | 4268 | 0.30075 | 0.66815 | No |
| PDPR | 4329 | 0.29588 | 0.67087 | No |
| LDHA | 4594 | 0.27905 | 0.66253 | No |
| UCP1 | 4734 | 0.26939 | 0.66057 | No |
| UQCRB | 4950 | 0.25566 | 0.65433 | No |
| SUCLG2 | 5240 | 0.23687 | 0.64383 | No |
| SLC16A8 | 5335 | 0.23139 | 0.64349 | No |
| IDH3A | 5350 | 0.23049 | 0.64733 | No |
| ETFA | 5494 | 0.22288 | 0.64424 | No |
| ATP5E | 5730 | 0.21029 | 0.63605 | No |
| NDUFA2 | 5828 | 0.20544 | 0.63503 | No |
| DLD | 6535 | 0.16489 | 0.60115 | No |
| ADHFE1 | 6751 | 0.15295 | 0.59288 | No |
| ATP5G1 | 6963 | 0.14294 | 0.58461 | No |
| D2HGDH | 7226 | 0.13095 | 0.57343 | No |
| SUCLA2 | 7299 | 0.12738 | 0.57217 | No |

| | | | | |
|---|---|---|---|---|
| UCP2 | 7927 | 0.09762 | 0.54111 | No |
| L2HGDH | 8225 | 0.08472 | 0.52716 | No |
| NDUFB3 | 8701 | 0.06244 | 0.50341 | No |
| ATP5J | 10339 | -0.00943 | 0.41744 | No |
| NNT | 11360 | -0.05772 | 0.36490 | No |
| IDH1 | 12233 | -0.09786 | 0.32095 | No |
| PDK3 | 13977 | -0.18731 | 0.23294 | No |
| PDP1 | 17559 | -0.50610 | 0.05453 | No |
| ATP5L | 18267 | -0.66981 | 0.03063 | No |
| NDUFS5 | 18339 | -0.69073 | 0.04063 | No |

**(iii)** **GO_CELLULAR_RESPIRATION** (Size: 132; ES: 0.62; NES: 2.37; NOM p-value: 0.000; FDR q-value: 0.000)

| **Probe** | **Rank In Gene List** | **Rank Metric Score** | **Running Es** | **Core Enrichment** |
|---|---|---|---|---|
| NDUFA12 | 338 | 1.11885 | 0.00438 | Yes |
| IMMP2L | 567 | 0.89436 | 0.01011 | Yes |
| COX4I2 | 791 | 0.77303 | 0.01369 | Yes |
| PDHB | 930 | 0.72104 | 0.02072 | Yes |
| NDUFB9 | 962 | 0.71030 | 0.03317 | Yes |
| NDUFA10 | 1082 | 0.67688 | 0.04033 | Yes |
| NDUFA8 | 1173 | 0.65806 | 0.04864 | Yes |
| COX6A1 | 1326 | 0.62148 | 0.05296 | Yes |
| NDUFS7 | 1337 | 0.61927 | 0.06471 | Yes |
| SDHB | 1398 | 0.60668 | 0.07358 | Yes |
| ACO2 | 1425 | 0.60040 | 0.08412 | Yes |
| NDUFA3 | 1445 | 0.59686 | 0.09496 | Yes |
| NDUFV2 | 1466 | 0.59397 | 0.10568 | Yes |
| NDUFB8 | 1470 | 0.59359 | 0.11730 | Yes |
| NDUFB11 | 1517 | 0.58410 | 0.12646 | Yes |
| UQCRH | 1530 | 0.58188 | 0.13737 | Yes |
| UQCRQ | 1542 | 0.57977 | 0.14829 | Yes |
| CYC1 | 1555 | 0.57708 | 0.15910 | Yes |
| UQCR10 | 1557 | 0.57677 | 0.17049 | Yes |
| NDUFB10 | 1633 | 0.56200 | 0.17768 | Yes |
| NDUFA6 | 1657 | 0.55869 | 0.18755 | Yes |
| IDH3B | 1667 | 0.55643 | 0.19811 | Yes |
| CYP1A2 | 1674 | 0.55564 | 0.20881 | Yes |
| NDUFV1 | 1705 | 0.55164 | 0.21817 | Yes |
| ETFB | 1728 | 0.54857 | 0.22790 | Yes |
| SDHD | 1735 | 0.54724 | 0.23843 | Yes |
| COX5B | 1757 | 0.54331 | 0.24810 | Yes |
| UQCRC2 | 1760 | 0.54307 | 0.25877 | Yes |
| NDUFS8 | 1790 | 0.53833 | 0.26792 | Yes |
| NDUFS3 | 1791 | 0.53820 | 0.27859 | Yes |
| COX5A | 1832 | 0.53190 | 0.28703 | Yes |
| UQCR11 | 1835 | 0.53169 | 0.29747 | Yes |
| COX4I1 | 1922 | 0.51948 | 0.30324 | Yes |
| CS | 1962 | 0.51432 | 0.31139 | Yes |
| SIRT3 | 1989 | 0.50968 | 0.32013 | Yes |
| NDUFA4 | 2008 | 0.50684 | 0.32923 | Yes |
| IDH3G | 2018 | 0.50537 | 0.33878 | Yes |
| UQCRC1 | 2043 | 0.50269 | 0.34749 | Yes |
| SDHC | 2054 | 0.50142 | 0.35690 | Yes |
| NDUFA7 | 2061 | 0.50077 | 0.36652 | Yes |
| COX6B1 | 2080 | 0.49895 | 0.37547 | Yes |
| SDHA | 2081 | 0.49873 | 0.38536 | Yes |
| NDUFAB1 | 2142 | 0.49114 | 0.39194 | Yes |
| UQCRFS1 | 2146 | 0.49090 | 0.40152 | Yes |
| NDUFA11 | 2189 | 0.48486 | 0.40892 | Yes |

| | | | | |
|---|---|---|---|---|
| DLST | 2224 | 0.47969 | 0.41664 | Yes |
| NDUFB2 | 2225 | 0.47950 | 0.42615 | Yes |
| NDUFC1 | 2238 | 0.47774 | 0.43499 | Yes |
| NDUFS2 | 2239 | 0.47765 | 0.44447 | Yes |
| NDUFA13 | 2287 | 0.47305 | 0.45137 | Yes |
| NDUFS6 | 2317 | 0.46973 | 0.45916 | Yes |
| OGDH | 2334 | 0.46756 | 0.46759 | Yes |
| NDUFV3 | 2401 | 0.46021 | 0.47324 | Yes |
| PINK1 | 2412 | 0.45877 | 0.48181 | Yes |
| COX7A2L | 2437 | 0.45540 | 0.48958 | Yes |
| NDUFA9 | 2541 | 0.44611 | 0.49300 | Yes |
| ETFDH | 2544 | 0.44590 | 0.50174 | Yes |
| COX10 | 2549 | 0.44535 | 0.51036 | Yes |
| NDUFC2 | 2555 | 0.44514 | 0.51893 | Yes |
| NDUFB6 | 2559 | 0.44459 | 0.52759 | Yes |
| COX7B | 2718 | 0.42880 | 0.52777 | Yes |
| MDH2 | 2722 | 0.42871 | 0.53611 | Yes |
| PDHA1 | 2727 | 0.42774 | 0.54438 | Yes |
| SUCLG1 | 2786 | 0.42214 | 0.54970 | Yes |
| FXN | 2889 | 0.41209 | 0.55250 | Yes |
| COQ9 | 2894 | 0.41195 | 0.56046 | Yes |
| DLAT | 2908 | 0.41111 | 0.56793 | Yes |
| OXA1L | 2934 | 0.40914 | 0.57473 | Yes |
| COX8A | 3015 | 0.40149 | 0.57847 | Yes |
| NDUFB5 | 3030 | 0.40032 | 0.58568 | Yes |
| COX7C | 3164 | 0.38996 | 0.58640 | Yes |
| COX6C | 3385 | 0.37129 | 0.58217 | Yes |
| CYCS | 3399 | 0.37034 | 0.58884 | Yes |
| SCO2 | 3598 | 0.35305 | 0.58541 | Yes |
| COX19 | 3658 | 0.34831 | 0.58920 | Yes |
| NDUFA5 | 3672 | 0.34736 | 0.59541 | Yes |
| NDUFB4 | 3721 | 0.34356 | 0.59969 | Yes |
| NDUFA1 | 3797 | 0.33689 | 0.60242 | Yes |
| NDUFB7 | 3979 | 0.32162 | 0.59927 | Yes |
| MDH1 | 4004 | 0.31968 | 0.60434 | Yes |
| NDUFS1 | 4014 | 0.31893 | 0.61019 | Yes |
| IDH2 | 4039 | 0.31755 | 0.61523 | Yes |
| GPD1 | 4169 | 0.30752 | 0.61453 | Yes |
| TBRG4 | 4178 | 0.30709 | 0.62020 | Yes |
| NDUFS4 | 4268 | 0.30075 | 0.62147 | Yes |
| TAZ | 4440 | 0.28888 | 0.61819 | Yes |
| SLC25A13 | 4463 | 0.28704 | 0.62273 | Yes |
| NDUFAF2 | 4768 | 0.26716 | 0.61201 | Yes |
| PMPCB | 4854 | 0.26159 | 0.61272 | Yes |
| SNCA | 4903 | 0.25862 | 0.61532 | Yes |
| ALDH5A1 | 4926 | 0.25680 | 0.61925 | Yes |
| UQCRB | 4950 | 0.25566 | 0.62311 | Yes |
| SUCLG2 | 5240 | 0.23687 | 0.61258 | No |
| IDH3A | 5350 | 0.23049 | 0.61141 | No |
| ETFA | 5494 | 0.22288 | 0.60830 | No |
| COX15 | 5754 | 0.20913 | 0.59880 | No |
| SDHAF2 | 5774 | 0.20808 | 0.60193 | No |
| NDUFA2 | 5828 | 0.20544 | 0.60321 | No |
| CHCHD5 | 5981 | 0.19658 | 0.59910 | No |
| SURF1 | 6487 | 0.16745 | 0.57581 | No |
| DLD | 6535 | 0.16489 | 0.57661 | No |
| ME3 | 7135 | 0.13519 | 0.54773 | No |
| NDUFAF1 | 7269 | 0.12878 | 0.54327 | No |
| SUCLA2 | 7299 | 0.12738 | 0.54427 | No |
| MTFR1 | 7420 | 0.12152 | 0.54036 | No |
| NFATC4 | 7434 | 0.12108 | 0.54208 | No |
| FASTKD1 | 7855 | 0.10050 | 0.52194 | No |

| | | | | |
|---|---|---|---|---|
| FASTKD2 | 8393 | 0.07654 | 0.49517 | No |
| PDHA2 | 8431 | 0.07445 | 0.49469 | No |
| MDH1B | 8522 | 0.07088 | 0.49136 | No |
| NDUFB3 | 8701 | 0.06244 | 0.48322 | No |
| BAX | 8817 | 0.05656 | 0.47828 | No |
| POLG2 | 9313 | 0.03490 | 0.45289 | No |
| PANK2 | 9532 | 0.02611 | 0.44193 | No |
| MYBBP1A | 9558 | 0.02474 | 0.44110 | No |
| SLC25A14 | 9754 | 0.01638 | 0.43115 | No |
| SLC25A12 | 10334 | -0.00928 | 0.40083 | No |
| ACO1 | 10480 | -0.01569 | 0.39350 | No |
| MECP2 | 10862 | -0.03359 | 0.37409 | No |
| FASTKD5 | 11002 | -0.04008 | 0.36756 | No |
| NNT | 11360 | -0.05772 | 0.34990 | No |
| BLOC1S1 | 12057 | -0.08916 | 0.31500 | No |
| IDH1 | 12233 | -0.09786 | 0.30772 | No |
| CAT | 12489 | -0.11166 | 0.29650 | No |
| DHTKD1 | 13535 | -0.16206 | 0.24465 | No |
| FASTKD3 | 13577 | -0.16392 | 0.24574 | No |
| PPARGC1A | 14719 | -0.23235 | 0.19024 | No |
| SLC1A3 | 15091 | -0.25704 | 0.17579 | No |
| SLC25A25 | 15167 | -0.26191 | 0.17703 | No |
| NR4A3 | 16571 | -0.38035 | 0.11065 | No |
| OGDHL | 17369 | -0.47223 | 0.07803 | No |
| NDUFS5 | 18339 | -0.69073 | 0.04067 | No |

**Table S5**: GSEA analysis of lipid metabolism related gene sets in sWAT of WT and Rbp$^{-/-}$ mice gene sets, including Hallmark and GO pathways.

**(i)** **HALLMARK_FATTY_ACID_METABOLISM** (Size: 145; ES: 0.49; NES: 1.89; NOM p-value: 0.000; FDR q-value: 0.021)

| Probe | Rank In Gene List | Rank Metric Score | Running Es | Core Enrichment |
|---|---|---|---|---|
| HMGCS2 | 274 | 1.23308 | 0.01086 | Yes |
| RDH11 | 406 | 1.03469 | 0.02519 | Yes |
| ACSM3 | 437 | 1.00655 | 0.04427 | Yes |
| ELOVL5 | 742 | 0.79933 | 0.04465 | Yes |
| ACAT2 | 818 | 0.75763 | 0.05625 | Yes |
| CRAT | 831 | 0.75074 | 0.07102 | Yes |
| FASN | 908 | 0.72769 | 0.08195 | Yes |
| PDHB | 930 | 0.72104 | 0.09565 | Yes |
| EHHADH | 933 | 0.72053 | 0.11033 | Yes |
| ECI1 | 939 | 0.71859 | 0.12482 | Yes |
| HPGD | 1090 | 0.67532 | 0.13077 | Yes |
| CIDEA | 1188 | 0.65553 | 0.13911 | Yes |
| GRHPR | 1210 | 0.64963 | 0.15134 | Yes |
| ALDH9A1 | 1323 | 0.62160 | 0.15819 | Yes |
| ACO2 | 1425 | 0.60040 | 0.16519 | Yes |
| SLC22A5 | 1583 | 0.57107 | 0.16863 | Yes |
| ME1 | 1585 | 0.57090 | 0.18030 | Yes |
| HAO2 | 1590 | 0.57010 | 0.19179 | Yes |
| IDH3B | 1667 | 0.55643 | 0.19920 | Yes |
| MLYCD | 1695 | 0.55274 | 0.20913 | Yes |
| GPD2 | 1719 | 0.55002 | 0.21920 | Yes |
| SDHD | 1735 | 0.54724 | 0.22964 | Yes |
| AQP7 | 1858 | 0.52909 | 0.23407 | Yes |
| DECR1 | 2007 | 0.50687 | 0.23667 | Yes |
| IDH3G | 2018 | 0.50537 | 0.24652 | Yes |
| SDHC | 2054 | 0.50142 | 0.25496 | Yes |
| SDHA | 2081 | 0.49873 | 0.26383 | Yes |
| VNN1 | 2130 | 0.49235 | 0.27141 | Yes |
| HMGCL | 2141 | 0.49120 | 0.28096 | Yes |
| ALDOA | 2162 | 0.48862 | 0.28994 | Yes |
| ACADL | 2164 | 0.48844 | 0.29991 | Yes |
| DLST | 2224 | 0.47969 | 0.30664 | Yes |
| ACOT2 | 2246 | 0.47706 | 0.31533 | Yes |
| HADH | 2256 | 0.47651 | 0.32463 | Yes |
| MCEE | 2280 | 0.47401 | 0.33315 | Yes |
| RETSAT | 2283 | 0.47356 | 0.34277 | Yes |
| HADHB | 2326 | 0.46847 | 0.35017 | Yes |
| ETFDH | 2544 | 0.44590 | 0.34788 | Yes |
| MDH2 | 2722 | 0.42871 | 0.34735 | Yes |
| PDHA1 | 2727 | 0.42774 | 0.35592 | Yes |
| ACADVL | 2763 | 0.42470 | 0.36279 | Yes |
| FABP1 | 2766 | 0.42393 | 0.37138 | Yes |
| SUCLG1 | 2786 | 0.42214 | 0.37905 | Yes |
| ECH1 | 2937 | 0.40902 | 0.37953 | Yes |
| PPARA | 2996 | 0.40330 | 0.38475 | Yes |
| ODC1 | 2998 | 0.40306 | 0.39297 | Yes |
| NSDHL | 3014 | 0.40158 | 0.40043 | Yes |
| ACOT8 | 3299 | 0.37796 | 0.39321 | Yes |
| MGLL | 3315 | 0.37654 | 0.40015 | Yes |
| MIF | 3354 | 0.37362 | 0.40581 | Yes |
| ADIPOR2 | 3592 | 0.35436 | 0.40059 | Yes |
| RDH16 | 3631 | 0.35092 | 0.40579 | Yes |
| ACAA2 | 3649 | 0.34925 | 0.41206 | Yes |

| | | | | |
|---|---|---|---|---|
| ACADS | 3752 | 0.34094 | 0.41368 | Yes |
| REEP6 | 3818 | 0.33453 | 0.41712 | Yes |
| HSD17B10 | 3922 | 0.32537 | 0.41837 | Yes |
| MDH1 | 4004 | 0.31968 | 0.42066 | Yes |
| ACSL5 | 4060 | 0.31626 | 0.42425 | Yes |
| IL4I1 | 4062 | 0.31577 | 0.43068 | Yes |
| GCDH | 4079 | 0.31422 | 0.43629 | Yes |
| GLUL | 4085 | 0.31407 | 0.44247 | Yes |
| HMGCS1 | 4128 | 0.31062 | 0.44663 | Yes |
| AADAT | 4133 | 0.31010 | 0.45279 | Yes |
| GPD1 | 4169 | 0.30752 | 0.45725 | Yes |
| DHCR24 | 4286 | 0.29941 | 0.45728 | Yes |
| ECI2 | 4289 | 0.29930 | 0.46332 | Yes |
| UROS | 4310 | 0.29718 | 0.46837 | Yes |
| UROD | 4348 | 0.29499 | 0.47247 | Yes |
| HSDL2 | 4585 | 0.27969 | 0.46577 | Yes |
| LDHA | 4594 | 0.27905 | 0.47107 | Yes |
| ECHS1 | 4609 | 0.27808 | 0.47604 | Yes |
| CRYZ | 4634 | 0.27691 | 0.48046 | Yes |
| S100A10 | 4799 | 0.26483 | 0.47725 | Yes |
| ACADM | 4974 | 0.25496 | 0.47331 | Yes |
| NTHL1 | 5057 | 0.24920 | 0.47410 | Yes |
| ADSL | 5183 | 0.24104 | 0.47246 | Yes |
| SUCLG2 | 5240 | 0.23687 | 0.47437 | Yes |
| ACSS1 | 5281 | 0.23488 | 0.47708 | Yes |
| AUH | 5413 | 0.22668 | 0.47483 | Yes |
| CBR1 | 5414 | 0.22664 | 0.47948 | Yes |
| CPT2 | 5451 | 0.22466 | 0.48219 | Yes |
| XIST | 5467 | 0.22404 | 0.48600 | Yes |
| CCDC58 | 5628 | 0.21643 | 0.48201 | No |
| NCAPH2 | 5704 | 0.21206 | 0.48240 | No |
| GABARAPL1 | 5811 | 0.20643 | 0.48105 | No |
| RAP1GDS1 | 6397 | 0.17311 | 0.45376 | No |
| DLD | 6535 | 0.16489 | 0.44993 | No |
| EPHX1 | 6678 | 0.15689 | 0.44566 | No |
| SETD8 | 6817 | 0.14954 | 0.44145 | No |
| HIBCH | 6831 | 0.14881 | 0.44382 | No |
| H2AFZ | 6964 | 0.14284 | 0.43979 | No |
| ENO3 | 7000 | 0.14166 | 0.44086 | No |
| D2HGDH | 7226 | 0.13095 | 0.43168 | No |
| SUCLA2 | 7299 | 0.12738 | 0.43050 | No |
| PSME1 | 7347 | 0.12537 | 0.43059 | No |
| HSD17B7 | 7479 | 0.11908 | 0.42613 | No |
| FMO1 | 7646 | 0.11106 | 0.41966 | No |
| ACOX1 | 7678 | 0.10953 | 0.42027 | No |
| ALAD | 7917 | 0.09815 | 0.40974 | No |
| HSD17B11 | 8197 | 0.08580 | 0.39679 | No |
| ACSL1 | 8414 | 0.07531 | 0.38695 | No |
| SMS | 8494 | 0.07207 | 0.38426 | No |
| HSPH1 | 8638 | 0.06528 | 0.37806 | No |
| BPHL | 8700 | 0.06248 | 0.37613 | No |
| ERP29 | 8719 | 0.06175 | 0.37645 | No |
| UGDH | 8988 | 0.04928 | 0.36333 | No |
| OSTC | 9213 | 0.03975 | 0.35234 | No |
| HCCS | 9848 | 0.01248 | 0.31917 | No |
| TDO2 | 9852 | 0.01241 | 0.31926 | No |
| CPOX | 10097 | 0.00200 | 0.30644 | No |
| ACSL4 | 10429 | -0.01338 | 0.28926 | No |
| BCKDHB | 10714 | -0.02665 | 0.27484 | No |
| UBE2L6 | 10748 | -0.02829 | 0.27368 | No |
| GSTZ1 | 10887 | -0.03455 | 0.26711 | No |
| HSD17B4 | 11378 | -0.05880 | 0.24248 | No |

| | | | | |
|---|---|---|---|---|
| LTC4S | 11456 | -0.06189 | 0.23969 | No |
| NBN | 11593 | -0.06884 | 0.23394 | No |
| PTS | 11968 | -0.08395 | 0.21594 | No |
| IDH1 | 12233 | -0.09786 | 0.20403 | No |
| BLVRA | 12345 | -0.10483 | 0.20033 | No |
| PTPRG | 12675 | -0.11959 | 0.18544 | No |
| ENO2 | 12709 | -0.12155 | 0.18619 | No |
| ALDH3A2 | 12889 | -0.13036 | 0.17943 | No |
| BMPR1B | 12998 | -0.13576 | 0.17652 | No |
| METAP1 | 13486 | -0.15942 | 0.15412 | No |
| AOC3 | 13646 | -0.16753 | 0.14918 | No |
| HSP90AA1 | 13999 | -0.18858 | 0.13449 | No |
| PRDX6 | 14374 | -0.20985 | 0.11908 | No |
| MAOA | 14437 | -0.21405 | 0.12020 | No |
| FABP2 | 14439 | -0.21437 | 0.12455 | No |
| ALDH1A1 | 14620 | -0.22596 | 0.11970 | No |
| LGALS1 | 14683 | -0.22978 | 0.12114 | No |
| SERINC1 | 14698 | -0.23065 | 0.12514 | No |
| IDI1 | 14715 | -0.23196 | 0.12906 | No |
| INMT | 15356 | -0.27526 | 0.10096 | No |
| CD36 | 15573 | -0.29071 | 0.09554 | No |
| APEX1 | 15608 | -0.29309 | 0.09977 | No |
| PCBD1 | 15900 | -0.31791 | 0.09095 | No |
| CPT1A | 16211 | -0.34491 | 0.08168 | No |
| CYP1A1 | 16512 | -0.37433 | 0.07355 | No |
| YWHAH | 16775 | -0.40259 | 0.06800 | No |
| ALDH3A1 | 17541 | -0.50033 | 0.03794 | No |
| CBR3 | 17667 | -0.52575 | 0.04214 | No |
| G0S2 | 18190 | -0.64596 | 0.02788 | No |
| ADH7 | 18829 | -1.00506 | 0.01487 | No |

**(ii) GO_ ENERGY_DERIVATION_BY_OXIDATION_OF_ORGANIC_COMPOUNDS**
(Size: 200; ES: 0.54; NES: 2.17; NOM p-value: 0.000; FDR q-value: 0.002)

| Probe | Rank In Gene List | Rank Metric Score | Running Es | Core Enrichment |
|---|---|---|---|---|
| KL | 148 | 1.77845 | 0.01684 | Yes |
| GCK | 246 | 1.30873 | 0.02986 | Yes |
| NDUFA12 | 338 | 1.11885 | 0.04056 | Yes |
| UBC | 540 | 0.91118 | 0.04257 | Yes |
| IMMP2L | 567 | 0.89436 | 0.05360 | Yes |
| PRKAG2 | 770 | 0.78381 | 0.05379 | Yes |
| COX4I2 | 791 | 0.77303 | 0.06345 | Yes |
| PDHB | 930 | 0.72104 | 0.06615 | Yes |
| NDUFB9 | 962 | 0.71030 | 0.07436 | Yes |
| NDUFA10 | 1082 | 0.67688 | 0.07746 | Yes |
| NDUFA8 | 1173 | 0.65806 | 0.08183 | Yes |
| PYGL | 1282 | 0.63289 | 0.08489 | Yes |
| COX6A1 | 1326 | 0.62148 | 0.09124 | Yes |
| NDUFS7 | 1337 | 0.61927 | 0.09930 | Yes |
| SDHB | 1398 | 0.60668 | 0.10454 | Yes |
| ACO2 | 1425 | 0.60040 | 0.11149 | Yes |
| NDUFA3 | 1445 | 0.59686 | 0.11876 | Yes |
| NDUFV2 | 1466 | 0.59397 | 0.12594 | Yes |
| NDUFB8 | 1470 | 0.59359 | 0.13401 | Yes |
| NDUFB11 | 1517 | 0.58410 | 0.13968 | Yes |
| UQCRH | 1530 | 0.58188 | 0.14712 | Yes |
| UQCRQ | 1542 | 0.57977 | 0.15457 | Yes |
| CYC1 | 1555 | 0.57708 | 0.16194 | Yes |
| UQCR10 | 1557 | 0.57677 | 0.16989 | Yes |
| NDUFB10 | 1633 | 0.56200 | 0.17372 | Yes |

| | | | | |
|---|---|---|---|---|
| NDUFA6 | 1657 | 0.55869 | 0.18025 | Yes |
| IDH3B | 1667 | 0.55643 | 0.18749 | Yes |
| CYP1A2 | 1674 | 0.55564 | 0.19488 | Yes |
| NDUFV1 | 1705 | 0.55164 | 0.20094 | Yes |
| ETFB | 1728 | 0.54857 | 0.20739 | Yes |
| SDHD | 1735 | 0.54724 | 0.21466 | Yes |
| COX5B | 1757 | 0.54331 | 0.22108 | Yes |
| UQCRC2 | 1760 | 0.54307 | 0.22851 | Yes |
| NDUFS8 | 1790 | 0.53833 | 0.23444 | Yes |
| NDUFS3 | 1791 | 0.53820 | 0.24190 | Yes |
| COX5A | 1832 | 0.53190 | 0.24716 | Yes |
| UQCR11 | 1835 | 0.53169 | 0.25443 | Yes |
| GYS2 | 1869 | 0.52755 | 0.26000 | Yes |
| G6PC | 1885 | 0.52456 | 0.26648 | Yes |
| COX4I1 | 1922 | 0.51948 | 0.27178 | Yes |
| CS | 1962 | 0.51432 | 0.27686 | Yes |
| SIRT3 | 1989 | 0.50968 | 0.28255 | Yes |
| NDUFA4 | 2008 | 0.50684 | 0.28863 | Yes |
| IDH3G | 2018 | 0.50537 | 0.29516 | Yes |
| UQCRC1 | 2043 | 0.50269 | 0.30086 | Yes |
| SDHC | 2054 | 0.50142 | 0.30729 | Yes |
| NDUFA7 | 2061 | 0.50077 | 0.31391 | Yes |
| COX6B1 | 2080 | 0.49895 | 0.31988 | Yes |
| SDHA | 2081 | 0.49873 | 0.32680 | Yes |
| NDUFAB1 | 2142 | 0.49114 | 0.33044 | Yes |
| UQCRFS1 | 2146 | 0.49090 | 0.33708 | Yes |
| NDUFA11 | 2189 | 0.48486 | 0.34159 | Yes |
| DLST | 2224 | 0.47969 | 0.34644 | Yes |
| NDUFB2 | 2225 | 0.47950 | 0.35309 | Yes |
| NDUFC1 | 2238 | 0.47774 | 0.35908 | Yes |
| NDUFS2 | 2239 | 0.47765 | 0.36571 | Yes |
| PPP1R1A | 2262 | 0.47576 | 0.37114 | Yes |
| NDUFA13 | 2287 | 0.47305 | 0.37643 | Yes |
| NDUFS6 | 2317 | 0.46973 | 0.38141 | Yes |
| OGDH | 2334 | 0.46756 | 0.38705 | Yes |
| NDUFV3 | 2401 | 0.46021 | 0.38994 | Yes |
| PINK1 | 2412 | 0.45877 | 0.39578 | Yes |
| COX7A2L | 2437 | 0.45540 | 0.40082 | Yes |
| GAA | 2463 | 0.45197 | 0.40577 | Yes |
| UBA52 | 2534 | 0.44683 | 0.40826 | Yes |
| NDUFA9 | 2541 | 0.44611 | 0.41413 | Yes |
| ETFDH | 2544 | 0.44590 | 0.42021 | Yes |
| COX10 | 2549 | 0.44535 | 0.42618 | Yes |
| NDUFC2 | 2555 | 0.44514 | 0.43209 | Yes |
| NDUFB6 | 2559 | 0.44459 | 0.43809 | Yes |
| STBD1 | 2618 | 0.43854 | 0.44111 | Yes |
| COX7B | 2718 | 0.42880 | 0.44182 | Yes |
| MDH2 | 2722 | 0.42871 | 0.44761 | Yes |
| PDHA1 | 2727 | 0.42774 | 0.45333 | Yes |
| GNMT | 2744 | 0.42649 | 0.45839 | Yes |
| ACADVL | 2763 | 0.42470 | 0.46333 | Yes |
| SUCLG1 | 2786 | 0.42214 | 0.46802 | Yes |
| FXN | 2889 | 0.41209 | 0.46834 | Yes |
| COQ9 | 2894 | 0.41195 | 0.47385 | Yes |
| DLAT | 2908 | 0.41111 | 0.47886 | Yes |
| OXA1L | 2934 | 0.40914 | 0.48321 | Yes |
| COX8A | 3015 | 0.40149 | 0.48455 | Yes |
| NDUFB5 | 3030 | 0.40032 | 0.48936 | Yes |
| MTOR | 3042 | 0.39952 | 0.49432 | Yes |
| GNAS | 3118 | 0.39349 | 0.49581 | Yes |
| COX7C | 3164 | 0.38996 | 0.49884 | Yes |
| COX6C | 3385 | 0.37129 | 0.49236 | Yes |

| | | | | |
|---|---|---|---|---|
| CYCS | 3399 | 0.37034 | 0.49680 | Yes |
| ACSM1 | 3572 | 0.35581 | 0.49264 | Yes |
| SCO2 | 3598 | 0.35305 | 0.49622 | Yes |
| COX19 | 3658 | 0.34831 | 0.49793 | Yes |
| NDUFA5 | 3672 | 0.34736 | 0.50206 | Yes |
| GSK3A | 3679 | 0.34658 | 0.50655 | Yes |
| NDUFB4 | 3721 | 0.34356 | 0.50914 | Yes |
| NDUFA1 | 3797 | 0.33689 | 0.50985 | Yes |
| STK40 | 3949 | 0.32365 | 0.50635 | Yes |
| NDUFB7 | 3979 | 0.32162 | 0.50928 | Yes |
| UGP2 | 4001 | 0.31982 | 0.51261 | Yes |
| MDH1 | 4004 | 0.31968 | 0.51693 | Yes |
| NDUFS1 | 4014 | 0.31893 | 0.52088 | Yes |
| IDH2 | 4039 | 0.31755 | 0.52401 | Yes |
| GPD1 | 4169 | 0.30752 | 0.52146 | Yes |
| TBRG4 | 4178 | 0.30709 | 0.52529 | Yes |
| PGM1 | 4233 | 0.30371 | 0.52665 | Yes |
| NDUFS4 | 4268 | 0.30075 | 0.52902 | Yes |
| PYGB | 4291 | 0.29882 | 0.53200 | Yes |
| TAZ | 4440 | 0.28888 | 0.52818 | Yes |
| SLC25A13 | 4463 | 0.28704 | 0.53100 | Yes |
| PGM2 | 4577 | 0.28029 | 0.52891 | Yes |
| PFKM | 4604 | 0.27845 | 0.53140 | Yes |
| PID1 | 4740 | 0.26901 | 0.52799 | Yes |
| NDUFAF2 | 4768 | 0.26716 | 0.53027 | Yes |
| GFPT2 | 4826 | 0.26311 | 0.53091 | Yes |
| PMPCB | 4854 | 0.26159 | 0.53311 | Yes |
| SNCA | 4903 | 0.25862 | 0.53415 | Yes |
| ALDH5A1 | 4926 | 0.25680 | 0.53655 | Yes |
| UQCRB | 4950 | 0.25566 | 0.53888 | Yes |
| AKT2 | 5227 | 0.23768 | 0.52758 | No |
| SUCLG2 | 5240 | 0.23687 | 0.53023 | No |
| IDH3A | 5350 | 0.23049 | 0.52767 | No |
| ETFA | 5494 | 0.22288 | 0.52320 | No |
| PRKAG3 | 5753 | 0.20916 | 0.51245 | No |
| COX15 | 5754 | 0.20913 | 0.51535 | No |
| SDHAF2 | 5774 | 0.20808 | 0.51724 | No |
| NDUFA2 | 5828 | 0.20544 | 0.51728 | No |
| CHCHD5 | 5981 | 0.19658 | 0.51197 | No |
| PPP1CA | 6120 | 0.18809 | 0.50728 | No |
| PHKG1 | 6444 | 0.16992 | 0.49256 | No |
| SURF1 | 6487 | 0.16745 | 0.49266 | No |
| DLD | 6535 | 0.16489 | 0.49246 | No |
| AGL | 6580 | 0.16244 | 0.49239 | No |
| GYS1 | 6792 | 0.15102 | 0.48333 | No |
| CPS1 | 7061 | 0.13859 | 0.47108 | No |
| ME3 | 7135 | 0.13519 | 0.46909 | No |
| NDUFAF1 | 7269 | 0.12878 | 0.46385 | No |
| SUCLA2 | 7299 | 0.12738 | 0.46408 | No |
| PHKG2 | 7333 | 0.12602 | 0.46408 | No |
| MTFR1 | 7420 | 0.12152 | 0.46122 | No |
| NFATC4 | 7434 | 0.12108 | 0.46221 | No |
| FASTKD1 | 7855 | 0.10050 | 0.44140 | No |
| PYGM | 7871 | 0.10006 | 0.44199 | No |
| NR1D1 | 7990 | 0.09512 | 0.43707 | No |
| NHLRC1 | 8200 | 0.08563 | 0.42721 | No |
| PPP1R3E | 8239 | 0.08391 | 0.42636 | No |
| GBE1 | 8327 | 0.07973 | 0.42287 | No |
| FASTKD2 | 8393 | 0.07654 | 0.42049 | No |
| PPP1R3B | 8397 | 0.07639 | 0.42139 | No |
| PDHA2 | 8431 | 0.07445 | 0.42068 | No |
| MDH1B | 8522 | 0.07088 | 0.41690 | No |

| | | | | |
|---|---|---|---|---|
| PPP1R2 | 8637 | 0.06532 | 0.41178 | No |
| NDUFB3 | 8701 | 0.06244 | 0.40932 | No |
| GFPT1 | 8786 | 0.05887 | 0.40569 | No |
| BAX | 8817 | 0.05656 | 0.40489 | No |
| PHKA1 | 9066 | 0.04564 | 0.39241 | No |
| POLG2 | 9313 | 0.03490 | 0.37989 | No |
| RPS27A | 9391 | 0.03162 | 0.37625 | No |
| PANK2 | 9532 | 0.02611 | 0.36921 | No |
| MYBBP1A | 9558 | 0.02474 | 0.36823 | No |
| AKT1 | 9624 | 0.02142 | 0.36509 | No |
| SLC25A14 | 9754 | 0.01638 | 0.35850 | No |
| PPP1R3A | 10001 | 0.00657 | 0.34558 | No |
| MRAP2 | 10235 | -0.00435 | 0.33332 | No |
| PPP1CC | 10258 | -0.00555 | 0.33224 | No |
| SLC25A12 | 10334 | -0.00928 | 0.32840 | No |
| ACO1 | 10480 | -0.01569 | 0.32095 | No |
| EPM2A | 10721 | -0.02706 | 0.30863 | No |
| PHKB | 10733 | -0.02756 | 0.30844 | No |
| MECP2 | 10862 | -0.03359 | 0.30213 | No |
| FASTKD5 | 11002 | -0.04008 | 0.29534 | No |
| NNT | 11360 | -0.05772 | 0.27726 | No |
| MYC | 11471 | -0.06254 | 0.27231 | No |
| GSK3B | 12012 | -0.08663 | 0.24496 | No |
| BLOC1S1 | 12057 | -0.08916 | 0.24387 | No |
| IDH1 | 12233 | -0.09786 | 0.23598 | No |
| ADRB3 | 12244 | -0.09894 | 0.23682 | No |
| CALM3 | 12451 | -0.10998 | 0.22745 | No |
| CAT | 12489 | -0.11166 | 0.22704 | No |
| IL6ST | 12507 | -0.11234 | 0.22770 | No |
| PHKA2 | 12752 | -0.12402 | 0.21652 | No |
| PCDH12 | 13168 | -0.14351 | 0.19657 | No |
| DHTKD1 | 13535 | -0.16206 | 0.17946 | No |
| PPP1CB | 13570 | -0.16349 | 0.17993 | No |
| FASTKD3 | 13577 | -0.16392 | 0.18189 | No |
| CALM2 | 14111 | -0.19439 | 0.15640 | No |
| PPP1R3C | 14661 | -0.22847 | 0.13054 | No |
| PPARGC1A | 14719 | -0.23235 | 0.13075 | No |
| LEPR | 14936 | -0.24628 | 0.12274 | No |
| SLC1A3 | 15091 | -0.25704 | 0.11816 | No |
| SLC25A25 | 15167 | -0.26191 | 0.11783 | No |
| CALM1 | 15493 | -0.28476 | 0.10459 | No |
| UBB | 16219 | -0.34537 | 0.07105 | No |
| NR4A3 | 16571 | -0.38035 | 0.05776 | No |
| MC4R | 17331 | -0.46623 | 0.02409 | No |
| OGDHL | 17369 | -0.47223 | 0.02869 | No |
| PPP1R3D | 17377 | -0.47404 | 0.03489 | No |
| PGM5 | 17658 | -0.52391 | 0.02735 | No |
| NDUFS5 | 18339 | -0.69073 | 0.00097 | No |
| LEP | 18596 | -0.81208 | -0.00130 | No |
| PER2 | 18695 | -0.87376 | 0.00564 | No |
| MT3 | 18902 | -1.17600 | 0.01105 | No |

**(iii)** **GO_CELLULAR_LIPID_CATABOLIC_PROCESS** (Size: 139; ES: 0.46; NES: 1.78; NOM p-value: 0.000; FDR q-value: 0.156)

| Probe | Rank In Gene List | Rank Metric Score | Running Es | Core Enrichment |
|---|---|---|---|---|
| FABP6 | 65 | 2.67224 | 0.05051 | Yes |
| PLA2G4F | 293 | 1.18771 | 0.06252 | Yes |

| | | | | |
|---|---|---|---|---|
| SMPD3 | 325 | 1.14531 | 0.08400 | Yes |
| ABHD6 | 477 | 0.96452 | 0.09551 | Yes |
| NEU2 | 550 | 0.90630 | 0.11000 | Yes |
| CYP26B1 | 573 | 0.89114 | 0.12683 | Yes |
| ACOX2 | 760 | 0.79078 | 0.13299 | Yes |
| DAGLA | 828 | 0.75206 | 0.14464 | Yes |
| CRAT | 831 | 0.75074 | 0.15968 | Yes |
| ACAD10 | 848 | 0.74522 | 0.17388 | Yes |
| LIPC | 913 | 0.72588 | 0.18516 | Yes |
| EHHADH | 933 | 0.72053 | 0.19870 | Yes |
| ECI1 | 939 | 0.71859 | 0.21294 | Yes |
| LIPG | 951 | 0.71445 | 0.22678 | Yes |
| PNPLA6 | 1211 | 0.64950 | 0.22624 | Yes |
| SLC27A2 | 1444 | 0.59707 | 0.22606 | Yes |
| APOC3 | 1527 | 0.58207 | 0.23349 | Yes |
| PLA2G15 | 1613 | 0.56714 | 0.24045 | Yes |
| PLA2G4B | 1722 | 0.54910 | 0.24584 | Yes |
| ETFB | 1728 | 0.54857 | 0.25665 | Yes |
| PHYH | 2002 | 0.50711 | 0.25250 | Yes |
| DECR1 | 2007 | 0.50687 | 0.26252 | Yes |
| PNLIPRP2 | 2019 | 0.50528 | 0.27214 | Yes |
| FABP7 | 2064 | 0.50045 | 0.27992 | Yes |
| FABP12 | 2071 | 0.49984 | 0.28969 | Yes |
| HADHA | 2138 | 0.49148 | 0.29613 | Yes |
| ACADL | 2164 | 0.48844 | 0.30467 | Yes |
| HADH | 2256 | 0.47651 | 0.30949 | Yes |
| MCEE | 2280 | 0.47401 | 0.31785 | Yes |
| FABP9 | 2298 | 0.47132 | 0.32646 | Yes |
| HADHB | 2326 | 0.46847 | 0.33450 | Yes |
| MMAA | 2339 | 0.46686 | 0.34329 | Yes |
| PNPLA2 | 2368 | 0.46392 | 0.35117 | Yes |
| ETFDH | 2544 | 0.44590 | 0.35095 | Yes |
| ACADVL | 2763 | 0.42470 | 0.34803 | Yes |
| FABP1 | 2766 | 0.42393 | 0.35648 | Yes |
| CPT1B | 2899 | 0.41160 | 0.35783 | Yes |
| ECH1 | 2937 | 0.40902 | 0.36414 | Yes |
| NUDT19 | 2951 | 0.40751 | 0.37168 | Yes |
| PLA2G4E | 2974 | 0.40571 | 0.37871 | Yes |
| APOA5 | 3081 | 0.39618 | 0.38112 | Yes |
| SLC25A17 | 3147 | 0.39094 | 0.38558 | Yes |
| PCCB | 3213 | 0.38526 | 0.38993 | Yes |
| PLCG2 | 3290 | 0.37846 | 0.39356 | Yes |
| ACOT8 | 3299 | 0.37796 | 0.40077 | Yes |
| MGLL | 3315 | 0.37654 | 0.40758 | Yes |
| ACAD8 | 3332 | 0.37518 | 0.41431 | Yes |
| APOB | 3348 | 0.37426 | 0.42107 | Yes |
| GDE1 | 3403 | 0.36996 | 0.42569 | Yes |
| APOA1 | 3514 | 0.36094 | 0.42718 | Yes |
| IVD | 3548 | 0.35733 | 0.43265 | Yes |
| FABP5 | 3575 | 0.35550 | 0.43846 | Yes |
| APOA2 | 3632 | 0.35087 | 0.44259 | Yes |
| PNPLA3 | 3692 | 0.34560 | 0.44645 | Yes |
| SGPL1 | 3704 | 0.34448 | 0.45283 | Yes |
| ACADS | 3752 | 0.34094 | 0.45723 | Yes |
| AMACR | 3804 | 0.33610 | 0.46133 | Yes |
| GCDH | 4079 | 0.31422 | 0.45323 | Yes |
| ECI2 | 4289 | 0.29930 | 0.44825 | Yes |
| ECHS1 | 4609 | 0.27808 | 0.43705 | Yes |
| BCO2 | 4685 | 0.27306 | 0.43861 | Yes |
| ABHD12 | 4690 | 0.27280 | 0.44390 | Yes |
| GM2A | 4848 | 0.26192 | 0.44092 | Yes |
| ACAD11 | 4949 | 0.25566 | 0.44081 | Yes |

| | | | | |
|---|---|---|---|---|
| NEU3 | 4960 | 0.25530 | 0.44543 | Yes |
| ANGPTL3 | 4965 | 0.25516 | 0.45037 | Yes |
| ACADM | 4974 | 0.25496 | 0.45510 | Yes |
| PEX2 | 5008 | 0.25280 | 0.45846 | Yes |
| FABP3 | 5235 | 0.23713 | 0.45133 | Yes |
| GALC | 5272 | 0.23528 | 0.45418 | Yes |
| SMPDL3A | 5382 | 0.22830 | 0.45305 | Yes |
| CPT2 | 5451 | 0.22466 | 0.45400 | Yes |
| LPIN2 | 5489 | 0.22306 | 0.45655 | Yes |
| ETFA | 5494 | 0.22288 | 0.46084 | Yes |
| CYP26A1 | 5504 | 0.22247 | 0.46485 | Yes |
| HAO1 | 5728 | 0.21057 | 0.45735 | No |
| PEX7 | 5773 | 0.20809 | 0.45923 | No |
| SLC27A4 | 6182 | 0.18483 | 0.44146 | No |
| ADIPOQ | 6329 | 0.17670 | 0.43733 | No |
| SESN2 | 6364 | 0.17510 | 0.43907 | No |
| ACOX3 | 6410 | 0.17240 | 0.44018 | No |
| APOE | 6497 | 0.16697 | 0.43901 | No |
| LPIN1 | 6738 | 0.15352 | 0.42946 | No |
| PLA2G4C | 6952 | 0.14350 | 0.42113 | No |
| SMPD1 | 6970 | 0.14255 | 0.42311 | No |
| ABCD1 | 7008 | 0.14143 | 0.42402 | No |
| CPS1 | 7061 | 0.13859 | 0.42408 | No |
| BDH2 | 7123 | 0.13572 | 0.42360 | No |
| PPARD | 7221 | 0.13117 | 0.42113 | No |
| AKR1B10 | 7640 | 0.11122 | 0.40135 | No |
| HACL1 | 7645 | 0.11111 | 0.40338 | No |
| ACOX1 | 7678 | 0.10953 | 0.40390 | No |
| NAGA | 7862 | 0.10032 | 0.39628 | No |
| CRABP1 | 7936 | 0.09714 | 0.39440 | No |
| PPT1 | 7967 | 0.09570 | 0.39475 | No |
| CPT1C | 8438 | 0.07419 | 0.37147 | No |
| SIRT2 | 8689 | 0.06317 | 0.35957 | No |
| SMPD4 | 8692 | 0.06299 | 0.36074 | No |
| GBA | 8758 | 0.06011 | 0.35852 | No |
| NEU1 | 8956 | 0.05096 | 0.34917 | No |
| FAAH | 9269 | 0.03671 | 0.33347 | No |
| PCCA | 9537 | 0.02581 | 0.31991 | No |
| GLA | 9692 | 0.01905 | 0.31218 | No |
| MUT | 9973 | 0.00758 | 0.29758 | No |
| CYP1B1 | 10233 | -0.00433 | 0.28401 | No |
| FABP4 | 10393 | -0.01185 | 0.27587 | No |
| LIPE | 10757 | -0.02870 | 0.25732 | No |
| HEXB | 10909 | -0.03554 | 0.25008 | No |
| ABHD2 | 10917 | -0.03576 | 0.25043 | No |
| GBA2 | 10929 | -0.03653 | 0.25059 | No |
| ENPP2 | 11252 | -0.05235 | 0.23467 | No |
| HSD17B4 | 11378 | -0.05880 | 0.22927 | No |
| ABCD3 | 11381 | -0.05895 | 0.23036 | No |
| GPCPD1 | 11561 | -0.06726 | 0.22228 | No |
| SCP2 | 11703 | -0.07291 | 0.21632 | No |
| ACADSB | 12130 | -0.09293 | 0.19574 | No |
| PLA2G4A | 12798 | -0.12608 | 0.16313 | No |
| CROT | 12802 | -0.12624 | 0.16552 | No |
| ALDH3A2 | 12889 | -0.13036 | 0.16362 | No |
| ENPP6 | 13110 | -0.14092 | 0.15487 | No |
| PNPLA8 | 13702 | -0.16983 | 0.12715 | No |
| NAPEPLD | 13888 | -0.18181 | 0.12106 | No |
| PLCG1 | 13896 | -0.18227 | 0.12437 | No |
| SPHK1 | 14177 | -0.19766 | 0.11361 | No |
| ACAD9 | 14179 | -0.19783 | 0.11755 | No |
| FABP2 | 14439 | -0.21437 | 0.10822 | No |

| | | | | |
|---|---|---|---|---|
| LPIN3 | 14920 | -0.24517 | 0.08787 | No |
| PEX13 | 14998 | -0.24986 | 0.08886 | No |
| INPP5F | 15280 | -0.26962 | 0.07949 | No |
| ACOXL | 15712 | -0.30299 | 0.06289 | No |
| CPT1A | 16211 | -0.34491 | 0.04360 | No |
| SRD5A3 | 16578 | -0.38122 | 0.03200 | No |
| SMPDL3B | 16769 | -0.40178 | 0.03010 | No |
| CYP26C1 | 17511 | -0.49554 | 0.00104 | No |
| LPL | 17885 | -0.57049 | -0.00710 | No |
| ABCD2 | 17929 | -0.57895 | 0.00232 | No |
| LEP | 18596 | -0.81208 | -0.01639 | No |
| ACER1 | 18808 | -0.97589 | -0.00782 | No |
| MT3 | 18902 | -1.17600 | 0.01102 | No |
